# Drug development of an affinity enhanced, broadly neutralizing heavy chain-only antibody that restricts SARS-CoV-2 in rodents

**DOI:** 10.1101/2021.03.08.433449

**Authors:** Bert Schepens, Loes van Schie, Wim Nerinckx, Kenny Roose, Wander Van Breedam, Daria Fijalkowska, Simon Devos, Wannes Weyts, Sieglinde De Cae, Sandrine Vanmarcke, Chiara Lonigro, Hannah Eeckhaut, Dries Van Herpe, Jimmy Borloo, Ana Filipa Oliveira, Joao Paulo Catani, Sarah Creytens, Dorien De Vlieger, Gitte Michielsen, Jackeline Cecilia Zavala Marchan, George D. Moschonas, Iebe Rossey, Koen Sedeyn, Annelies Van Hecke, Xin Zhang, Lana Langendries, Sofie Jacobs, Sebastiaan ter Horst, Laura Seldeslachts, Laurens Liesenborghs, Robbert Boudewijns, Hendrik Jan Thibaut, Kai Dallmeier, Greetje Vande Velde, Birgit Weynand, Julius Beer, Daniel Schnepf, Annette Ohnemus, Isabel Remory, Caroline S. Foo, Rana Abdelnabi, Piet Maes, Suzanne J. F. Kaptein, Joana Rocha-Pereira, Dirk Jochmans, Leen Delang, Frank Peelman, Peter Staeheli, Martin Schwemmle, Nick Devoogdt, Dominique Tersago, Massimiliano Germani, James Heads, Alistair Henry, Andrew Popplewell, Mark Ellis, Kevin Brady, Alison Turner, Bruno Dombrecht, Catelijne Stortelers, Johan Neyts, Nico Callewaert, Xavier Saelens

**Affiliations:** VIB-UGent Center for Medical Biotechnology, VIB, Technologiepark-Zwijnaarde 75, 9052 Ghent, Belgium; Department of Biochemistry and Microbiology, Ghent University, Technologiepark-Zwijnaarde 75, 9052 Ghent, Belgium; VIB Discovery Sciences, Technologiepark 104B, 9052 Ghent, Belgium; KU Leuven Department of Microbiology, Immunology and Transplantation, Rega Institute, Laboratory of Virology and Chemotherapy, BE-3000 Leuven, Belgium; GVN, Global Virus Network; KU Leuven Department of Imaging and Pathology, Biomedical MRI and MoSAIC, B-3000 Leuven, Belgium; Molecular Vaccinology and Vaccine Discovery Group; Translational Platform Virology and Chemotherapy (TPVC); KU Leuven Department of Imaging and Pathology, Translational Cell and Tissue Research, B-3000 Leuven, Belgium; Division of Translational Cell and Tissue Research; Institute of Virology, Medical Center University Freiburg, Freiburg, Germany; Department of Medical Imaging, In vivo Cellular and Molecular Imaging laboratory, Vrije Universiteit Brussel, Laarbeeklaan 103, 1090 Brussels, Belgium; Laboratory of Clinical and Epidemiological Virology, Rega Institute, KU Leuven, Department of 15 Microbiology, Immunology and Transplantation, 3000, Leuven, Belgium; Zoonotic Infectious Diseases Unit, Leuven, Belgium; Department of Biomolecular Medicine, Ghent University, 9000 Ghent, Belgium; Faculty of Medicine, University of Freiburg, Freiburg, Germany; ExeVir, Rijvisschestraat 120, 9052 Ghent, Belgium; UCB Celltech, Slough, UK

## Abstract

We have identified camelid single-domain antibodies (VHHs) that cross-neutralize SARS-CoV-1 and −2, such as VHH72, which binds to a unique highly conserved epitope in the viral receptor-binding domain (RBD) that is difficult to access for human antibodies. Here, we establish a protein engineering path for how a stable, long-acting drug candidate can be generated out of such a VHH building block. When fused to human IgG1-Fc, the prototype VHH72 molecule prophylactically protects hamsters from SARS-CoV-2. In addition, we demonstrate that both systemic and intranasal application protects hACE-2-transgenic mice from SARS-CoV-2 induced lethal disease progression. To boost potency of the lead, we used structure-guided molecular modeling combined with rapid yeast-based Fc-fusion prototyping, resulting in the affinity-matured VHH72_S56A-Fc, with subnanomolar SARS-CoV-1 and −2 neutralizing potency. Upon humanization, VHH72_S56A was fused to a human IgG1 Fc with optimized manufacturing homogeneity and silenced effector functions for enhanced safety, and its stability as well as lack of off-target binding was extensively characterized. Therapeutic systemic administration of a low dose of VHH72_S56A-Fc antibodies strongly restricted replication of both original and D614G mutant variants of SARS-CoV-2 virus in hamsters, and minimized the development of lung damage. This work led to the selection of XVR011 for clinical development, a highly stable anti-COVID-19 biologic with excellent manufacturability. Additionally, we show that XVR011 is unaffected in its neutralizing capacity of currently rapidly spreading SARS-CoV-2 variants, and demonstrate its unique, wide scope of binding across the Sarbecovirus clades.

---

Severe acute respiratory syndrome Coronavirus 2 (SARS-CoV-2) is the causative agent of COVID-19, a disease that has rapidly spread across the planet with devastating consequences^1^. SARS-CoV-2 infections can be asymptomatic and mostly present mild to moderately severe symptoms. However, in approximately 10% of patients, COVID-19 progresses to a more severe stage that is characterized by dyspnea and hypoxemia, which may progress further to acute respiratory distress often requiring long-term intensive care and causing death in a proportion of patients. The ongoing inflammation triggered by the innate recognition of the SARS-CoV-2 virus^2^ contributes to severe disease progression.

Prophylactic vaccine candidates have been developed at an unprecedented speed and viral spike-encoding mRNA and adenovirus-vectored vaccines have already obtained emergency approval in many countries^3, 4^. COVID-19 vaccines will become the cornerstone of controlling the pandemic yet will likely leave a significant part of the population insufficiently protected. Immunity may be short-lived and vaccine efficacy may be lower in the elderly, the age group that is most at risk of developing severe COVID-19^5, 6^. Limited vaccine availability in many countries, vaccine hesitancy and viral evolution to escape human immunity^7^ are other factors of which the impact is currently uncertain. Hence, passive antibody immunotherapy with broadly neutralizing molecules, to prevent or suppress viral replication in the lower airways, will likely find an important place in rescuing COVID-19 patients. Indeed, the early development of sufficient titers of neutralizing antibodies by the patient correlates with avoidance of progression to severe disease^8^, and early administration of recombinant neutralizing antibodies or those present in high-titer convalescent plasma can avert severe disease^9–11^. An advantage of antibodies and antibody Fc-based fusions compared with small molecule drugs is their long circulatory half-life imparted by the FcRn-mediated recycling into the bloodstream, which provides for long term control of virus replication even after a single administration^12^.

Developed at an extraordinary pace, Regeneron’s REGN10933/REGN10987 pair (Casirivimab/Imdevimab) and Eli Lilly’s LyCoV555 (Bamlanivimab) recently obtained FDA emergency use approval for the treatment of mild to moderate COVID-19 in non-hospitalized adults and pediatric patients who are at risk of developing severe disease^10, 13^. Since its first emergence in humans at the end of 2019, however, SARS-CoV-2 viruses have acquired mutations that indicate further adaptation to the human host. Some of these SARS-CoV-2 variants are of concern (VOCs) because they are associated with increased transmissibility or contribute to evasion of host immunity^14^. These circulating mutant viruses often have acquired mutations in the receptor-binding motif (RBM), the region of the RBD that interacts with hACE2 that is also the target of most of the reported neutralizing antibodies in immune plasma and monoclonal antibodies derived from convalescent patients. As a consequence, these SARS-CoV-2 variants display increased resistance to neutralization by many convalescent plasma-derived monoclonal antibodies, including two of the three that have received emergency use approval in the USA^15, 16^. This reinforces the need to develop neutralizing antibodies that bind to regions of the SARS-CoV-2 spike that are under lower human immune pressure and that poorly tolerate mutations without compromising virus fitness. Such broadly neutralizing antibodies would also constitute valuable tools for our pandemic preparedness for future outbreaks of viruses in this family.

We recently reported the discovery and structural characterization of a VHH (the variable domain of a heavy chain-only antibody, also known as a Nanobody), named SARS VHH-72 (here abbreviated as VHH72), with SARS-CoV-1 and −2 neutralizing capacity^17^. VHH72-Fc was the first SARS-CoV-2 neutralizing antibody that binds to the RBD region that partially overlaps with the epitope of the anti-SARS-CoV-1 antibody CR3022, which is however not ACE2-competing and non-neutralizing against SARS-CoV-2. VHH72 binds to an epitope in the receptor-binding domain (RBD) of the spike protein that is highly conserved in members of the Sarbecovirus subgenus of the Betacoronaviruses^18, 19^, prevents the interaction of the SARS-CoV-1 and −2 RBD with ACE2, and, by trapping the RBD in the “up” conformation, presumably destabilizes the spike protein of SARS-CoV-1 and −2^17^. Prophylactic administration of a prototype VHH72-human IgG1 Fc domain fusion strongly restricted SARS-CoV-2 replication in the lungs of experimentally infected golden Syrian hamsters^20^. The VHH72 contact region is occluded in the closed spike conformation, which is the dominant one on the native virus^21^. Even in the ‘1-RBD-up’ conformation that can bind the ACE2 receptor, the epitope is positioned such that human monoclonal antibodies cannot easily reach it. Possibly because of this, amidst hundreds of antibodies against other regions of the spike, very few human antibodies have been reported that bind to an epitope that substantially overlaps the VHH72 epitope^22^. Moreover, the VHH72 contact region on the RBD is comprised of residues that form crucial packing contacts between the protomers of the trimeric spike. SARS-CoV-2 viruses with mutations in this contact region remain extremely rare. Consistently, as we also illustrate in the present paper, none of the recently emerging and rapidly spreading VOC RBD mutations are in the VHH72 epitope.

Antibodies that cross-neutralize SARS-CoV-1 and −2 and other viruses of the Sarbecovirus subgenus, are very rare. Therefore, we decided to build on the VHH72 discovery, to generate a potency-enhanced drug designed for safety and long circulatory half-life, with a target product profile suitable for both treatment and prophylaxis of infections by SARS-CoV-2 and related viruses. This resulted in the design of the fully optimized XVR011, which is presently entering clinical development. Whereas the initial purpose is systemic administration to patients at risk of aggravated disease, we also show that nasal delivery of VHH72-Fc fully protects in a mouse model against SARS-CoV-2 challenge infection.

Many laboratories now have the capability to rapidly identify and characterize camelid single-domain VHH binders, either from *in vivo* immunizations or from selections from *in vitro* displayed repertoires, and the platform has hence been widely deployed in the present SARS-CoV-2 crisis^17, 23–31^. However, a binder, no matter how potent, is not necessarily a drug. In a broader sense, our report on the entire lead molecule development track from our VHH72 pre-lead molecule to a fully optimized VHH-Fc drug substance, for the first time provides a detailed description of the development track that is needed from such prototype anti-viral VHH into a VHH-hIgG1-Fc antiviral clinical lead drug candidate, which may guide other such efforts and contribute to our pandemic readiness in the future.

## Results

### Prototype VHH72-Fc protects K18-hACE2 mice from SARS-CoV-2 induced disease

We previously demonstrated protection in the Syrian hamster SARS-CoV-2 infection model by our prototype VHH72-Fc (here referred to as WT-VHH/12GS-WT-Fc; for an overview of the purified, CHO-produced VHH72-Fc constructs described in this study we refer to Supplementary Table 1)^20^. To gain more confidence, we also explored its protective efficacy in a mouse model expressing human ACE2 under the control of the cytokeratin 18 promoter (K18-hACE2), which results in susceptibility to SARS-CoV-1 and −2 infection^32–34^. K18-hACE2 transgenic mice intraperitoneally injected with 5 mg/kg of WT-VHH/12GS-WT-Fc and challenged 7 h later with a dose of 8×10^3^ plaque forming units of a clinical isolate of SARS-CoV-2 displayed no body weight loss, survived the challenge and had significantly reduced lung virus titers on day 3 after infection compared to control treated animals (Fig. 1a-c). While systemic intravenous administration of SARS-CoV-2 neutralizing antibodies is currently the main pursued route to treat patients with a high risk of aggravating disease, in a setting such as post-exposure prophylaxis of close contacts of a patient, administration into the respiratory tract through inhalation may offer benefits such as reduced dose requirement and avoiding the need for intravenous injections. Such inhaled administration can be mimicked by intranasal delivery in the sedated mouse model, and we tested this for our prototype WT_VHH/12GS-WT-Fc molecule. Intranasal administration of WT-VHH/12GS-WT-Fc at a dose of 20 µg per mouse (approximately 1 mg/kg) provided full protection against SARS-CoV-2 induced disease (Fig. 1d-f). Viral titers in the lungs were also significantly reduced in WT-VHH/12GS-WT-Fc-treated mice when compared with the controls (Fig. 1d-f). These data and our earlier report^20^ indicated that the pre-lead prototype molecule with VHH72 specificity in the context of the half-life prolonging and bivalency-imparting hIgG1 Fc fusion had the potential to robustly protect against SARS-CoV-2 virus challenge in a prophylactic setting in two different animal models following parenteral and intranasal administration.

**Figure 1.**
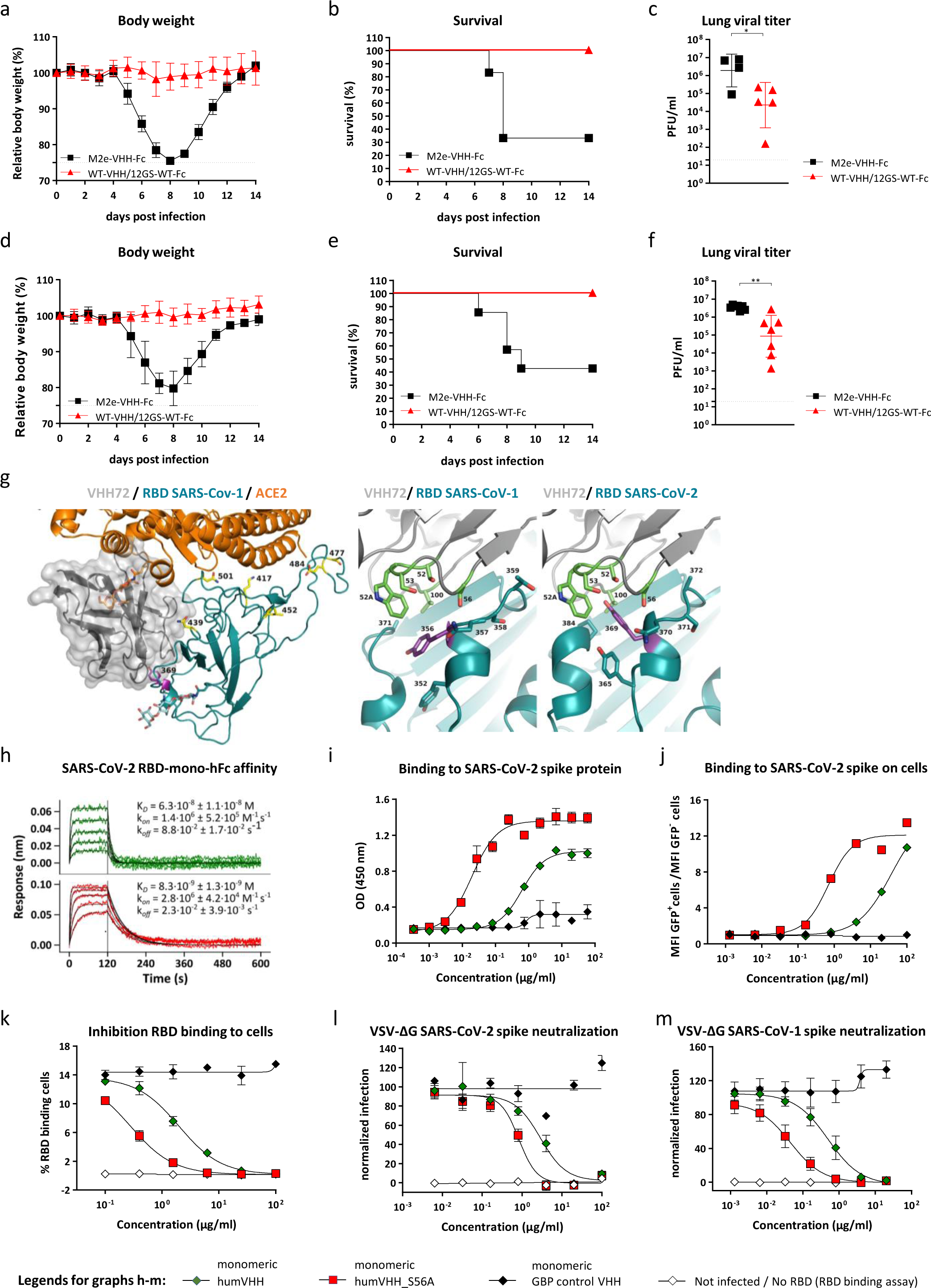
Enhanced affinity and neutralizing activity of computationally predicted VHH72 variant. **a-c.** Prototype WT-VHH/12GS-WT-Fc protects K18-hACE2 transgenic mice against SARS-CoV-2 challenge. Mice were injected intraperitoneally with 5 mg/kg of either WT-VHH/12GS-WT-Fc (n = 17) or control M2e-VHH-Fc (directed against the matrix protein 2 ectodomain of influenza A; n = 10). Seven hours later, the mice were challenged with 8 x 10^3^ PFU of a clinical SARS-CoV-2 isolate. (**a**) Body weight (symbols represent means ± SD, p < 0.0001, 2-way ANOVA), (**b**) survival over time after challenge (p = 0.0015, Mantel-Cox), and (**c**) lung viral loads determined on day 3 after infection from 4 or 5 mice sacrificed of each group. Symbols represent individual mice. Lines indicate means ± SD (*p<0.05, unpaired t test; data in a-b are pooled from two independently performed experiments). **d-f**. K18-hACE2 transgenic mice received 20 µg of either WT-VHH/12GS-WT-Fc (n = 15) or M2e-VHH-Fc (n = 14) intranasally in a 40 µl volume. Seven hours later, the mice were infected with 8×10^3^ PFU of a clinical SARS-CoV-2 isolate. Animals were monitored for (**d**) weight loss (symbols represent means ± SD, p < 0.0001, 2-way ANOVA) and (**e**) survival (p = 0.0148, Mantel-Cox). (**f**) After 72 h, seven mice of each roup were sacrificed to determine viral titers in their lungs. Symbols represent individual mice. Lines indicate means ± SD. Statistical analysis: **p<0.001 unpaired t test. **g.** Left: Composite overlay showing the locations of VHH72 (grey cartoon with transparent surface, centre-left) and ACE-2 (orange cartoon, top) versus SARS-CoV-2 RBD (cyan cartoon, centre). Tyr369 of SARS-CoV-2 RBD is indicated and shown as purple sticks. The protein-proximal monosaccharides of the ACE-2 N322 N-glycan (clashing with VHH72) are shown as orange sticks; the RBD N343 N-glycan’s protein-proximal monosaccharides are shown as cyan sticks. The emerging RBD variants at residues K417(->N), N439(->K), L452(->R), S477(->N), E484(->K) and N501(->Y) are indicated and shown as yellow sticks. Right: Comparison of VHH72 (rainbow cartoon) in complex with SARS-CoV-1 RBD (cyan cartoon, pdb-entry 6WAQ chains C and D) with a homology model of VHH72 bound to SARS-CoV-2 RBD (cyan cartoon, model obtained from the I-TASSER server), zoomed-in to the zone near VHH72’s S56. VHH72 residues S52, W52a, S53, S56 and V100, SARS-CoV-1 RBD residues Y352, Y356 (purple), N357, S358, T359 and A371, and SARS-CoV-2 RBD residues Y365, Y369 (purple), N370, S371, A372 and P384 are shown as sticks. Figures generated with Pymol (The PyMOL Molecular Graphics System, Open Source Version 2.3. Schrödinger, LLC). RBD Tyr369 assumes a differential preferential conformation between SARS-CoV-1 and SARS-CoV-2, imposed by the P384 in SARS-CoV-2. Accommodating this alteration was the focus of our structure-guided affinity maturation campaign of VHH72. **h.** BLI sensorgrams of humVHH (top) and humVHH_S56A (bottom) (twofold dilution series starting at 100 nM) binding to immobilized monomeric human IgG Fc-fused RBD from SARS-CoV-2. Green and red lines represent double reference-subtracted data and the fit of the data to a 1:1 binding curve is in black. **i.** Binding of humVHH and humVHH_S56A to SARS-CoV-2 spike determined by ELISA (data points are mean ± SD; n=3). **j.** Binding of humVHH and humVHH_S56A to HEK293 cell surface expressed SARS-CoV-2 spike determined by flow cytometry. The graph shows the ratio of the MFI of transfected (GFP^+^) cells over the MFI of non-transfected (GFP^-^) cells. **k.** Dose-dependent inhibition of SARS-CoV-2 RBD binding to the surface of VeroE6 cells determined by flow cytometry. **l,m.** Neutralization of SARS-CoV-2 (**l**) and −1 (**m**) S pseudotyped VSV. The graphs show the mean (n=4 ± SD) GFP fluorescence normalized to the mean GFP fluorescence of non-infected (NI) and infected PBS-treated cells (PBS). GBP: GFP-binding protein = a VHH directed against GFP.

### VHH72 binds a highly conserved region in the RBD of SARS-CoV-2

VHH72 binds to a region in the core of the RBD that is distal from the much more variable RBM^17^. Free energy contribution analysis by FastContact^35^ of snapshots from Molecular Dynamics simulations with the VHH72-RBD complex indicates that the epitope recognized by VHH72 has a prominent two-residue hot-spot, consisting of F377 and K378, which contact VHH72 residues V100 and D100g, respectively (Supplementary Fig. 1a). The epitope is exposed only when the trimeric spike protein has at least one RBD in an ‘up’ conformation (Supplementary Fig. 1b)^17^. In the three-RBD ‘down’ state, the VHH72 contact region belongs to an occluded zone that makes mutual contacts with the adjacent RBDs (Supplementary Fig. 1c), as well as with the helix-turn-helix positioned between heptad repeat 1 and the central helix of the underlying S2 domain. These subtle inter-RBD and inter-S1/S2 contacts allow oscillation between the RBD ‘down’ and the ACE2-engaging ‘up’ positioning^36, 37^. Presumably due to this involvement in spike conformational dynamics, the amino acid residues that contribute to the VHH72 contact region are remarkably conserved in circulating SARS-CoV-2 viruses (Supplementary Fig. 2). Moreover, deep mutational scanning analysis has shown that the VHH72 contact region overlaps with a patch of the RBD in which mutations may severely compromise the fold^38^, further explaining why it is indeed a highly conserved region on the Sarbecoviral RBD.

### Design and selection of a VHH72 variant with increased neutralizing activity

While the prototype VHH72-Fc molecule had promising *in vivo* efficacy, we wished to further enhance its potency, with an eye towards ease of administration at reduced dosage through various routes, including subcutaneous and inhaled. Given the limited available time in this emergency lead development campaign, we decided on a protein modeling based mutant design approach for a VHH72 variant with increased affinity for SARS-CoV-2 RBD. At the start of our investigation, immediately after the SARS-CoV-2 genome was published, however, no SARS-CoV-2 RBD structure was publicly available. Therefore, based on the crystal structure of VHH72 in complex with the SARS-CoV-1 RBD (PDB code: 6WAQ), a model of the SARS-CoV-2 RBD was generated through the I-TASSER server^39^, which was then superimposed by means of the Swiss-PdbViewer^40^ to the SARS-CoV-1 RBD/VHH72 structure (Fig. 1g). Only three residues are different between SARS-CoV-2 and −1 at the VHH72-RBD interface: (1) A372 (T359 in SARS-CoV-1), resulting in the loss of a glycan on N370 (N357 in SARS-CoV-1); (2) N439 (R426 in SARS-CoV-1), resulting in the loss of an ionic interaction with VHH72 residue D61; and (3) P384 (A371 in SARS-CoV-1). Close to P384 is Y369, which is a key contact residue for VHH72 (Supplementary Fig. 1), for which I-TASSER predicted an upward conformation in the SARS-CoV-2 RBD-VHH72 model (Fig. 1g). In the SARS-CoV-1 RBD-VHH72 co-crystal structure, however, the corresponding Y356 points downward and resides in a groove-like depression between two short helices of the RBD. The up-conformation of SARS-CoV-2 Y369 sets it in a mostly hydrophobic small cavity of VHH72, contacting residues S52, W52a, S53, S56 (all in CDR2) and V100 (in CDR3) (Fig. 1g). Molecular dynamics simulations with Gromacs^41^ shows that Y369 can be readily accommodated in that cavity. Interestingly, the later reported cryo-EM or crystal structures of SARS-CoV-2 RBD indeed typically show Y369 in the upward conformation (*e.g*. PDB-entries 6VSB, 6M17, and 6VXX)^42–44^. Presumably Y369, as well as its Y356 counterpart in SARS-CoV-1 RBD, can flip into up or down positions, with the up-position prevailing in SARS-CoV-2 RBD due to a conformational constraint imposed by the nearby P384, which is A371 in SARS-CoV-1.

Guided by molecular modeling, we generated a set of VHH72 variants with point mutations in the residues that line the cavity that accommodates Y369 of SARS-CoV-2 RBD. To ensure that we would only prioritize mutations that enhanced affinity in our intended bivalent Fc-fusion drug context, these variants were rapidly prototyped in parallel as Fc fusions in *Pichia pastoris*, which can produce VHH-Fc fusions much more easily than native hIgG1s. The crude yeast medium was screened for binding to immobilized SARS-CoV-2 RBD by biolayer interferometry (BLI). Introduction of S56A resulted in a slower off-rate in this bivalent context (Supplementary Fig. 3a,b). VHH_S56A/12GS-WT-Fc also displayed higher binding affinity for SARS-CoV-2 spike expressed on the surface of 293T cells than parental WT-VHH/12GS-WT-Fc (Supplementary Fig. 3c).

Next, we humanized the monovalent VHH72 with or without the S56A mutation, based on a sequence comparison with the human IGHV3-JH consensus sequence, and the N-terminal glutamine was replaced by a glutamic acid codon to avoid N-terminal pyroglutamate formation, in order to increase the chemical homogeneity (Supplementary Fig. 4). As with its Fc fusion, we validated that the S56A substitution increased the affinity of the monomeric humanized VHH72 for immobilized SARS-CoV-2 RBD, by approximately seven-fold as determined by BLI (Fig. 1h). Increased affinity of humVHH_S56A for prefusion stabilized SARS-CoV-2 spike was also observed in ELISA and for SARS-CoV-2 spike expressed on the surface of mammalian cells by flow cytometry (Fig. 1i,j). In addition, the mutant prevented the binding of SARS-CoV-2 RBD to ACE2 on the surface of VeroE6 cells seven times more efficiently than humVHH (Fig. 1k). This improved affinity correlated with significantly stronger neutralizing activity of Hum_S56A as determined in a VSV SARS-CoV-2 spike pseudotyped virus neutralization assay (IC_50_ humVHH: 2.550 µg/ml; IC_50_ humVHH_S56A: 0.837 µg/ml) (Fig. 1l). Importantly, humVHH_S56A also neutralized VSV SARS-CoV-1 spike pseudotypes with 10-fold higher potency and bound with higher affinity to SARS-CoV-1 RBD and spike than humVHH (IC_50_ humVHH: 0.491 µg/ml; IC_50_ humVHH_S56A: 0.045 µg/ml) (Fig. 1m and Supplementary Fig. 5).

### VHH72_S56A-Fc constructs with potent SARS-CoV-2 neutralizing activity

We opted to include a human IgG1 with minimal Fc effector functions in our VHH72-Fc designs because there is uncertainty about the possible contribution of IgG effector functions to disease severity in COVID-19 patients^45–47^. To this effect, and as also chosen by several other anti-SARS-CoV-2 antibody developers^48, 49^, we opted to use the well-characterized LALA mutations in the Fc portion, with or without the P329G mutation (LALAPG)^50–52^. In an initial set of experiments, we validated that neither the Gly-Ser linker length between the VHH and the Fc hinge (2 or 14 amino acids), nor the humanization of the VHH nor the introduction of LALAPG mutations in the Fc affected the affinity for SARS-CoV-2 S or its RBD, as determined by BLI, ELISA, flow cytometry and an ACE2 competition assay (Fig. 2a-e and Supplementary Fig. 6b,c). Consistent with this, potency (PRNT_50_) in a plaque neutralization assay using authentic SARS-CoV-2 virus was unaffected by these changes in the linker length or the Fc part (Supplementary Table 1). Subsequently, we built the S56A humanized VHH variants of the WT and LALAPG-Fc-fusions. Consistent with the results obtained with the *Pichia*-produced prototypes, we observed 2-3-fold higher affinity for immobilized bivalent SARS-CoV-2 RBD as determined in BLI, which was further confirmed in flow cytometry-based quantification assays using mammalian cells that express spike on their surface, in hACE2-Fc RBD competition AlphaLISA, and in a VeroE6 cell-based SARS-CoV-2 RBD competition assay (Fig. 2a-e and Supplementary Fig. 6a-e). This enhancement of spike binding affinity resulted in approximately seven-fold enhanced potency in the authentic SARS-CoV-2 neutralization assay (*e.g.* humVHH_S56A/LALAPG-Fc: 0.13 µg/ml compared with humVHH/LALAPG-Fc: 1.22 µg/ml) (Fig. 2f and Supplementary Table 1).

**Figure 2.**
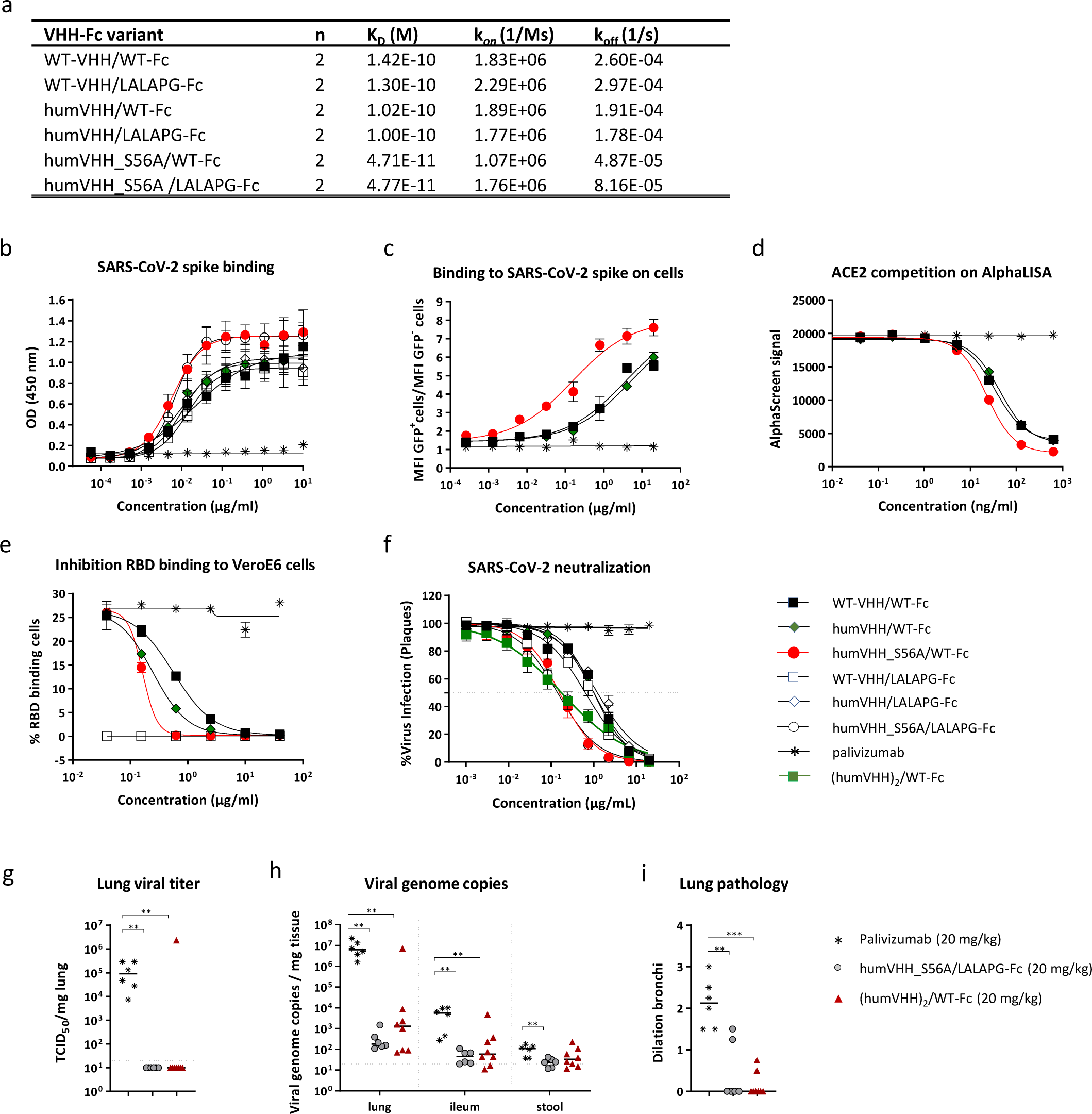
VHH72_S56A-Fc constructs have increased affinity and SARS-CoV-2 neutralizing activity. **a.** Binding affinity of VHH72-Fc variants to immobilized mouse Fc-fused SARS-CoV-2 RBD (RBD-mFc). Apparent kinetics of the 2:2 interaction is based on a global 1:1 fit of the replicate (n = 2) data; values are the averages of replicates. **b.** Binding of the indicated VHH72-Fc constructs (see Supplementary Table 2 for a description) to coated SARS-CoV-2 spike determined by ELISA (data points are mean ± SD; n=3). **c.** Binding of the indicated VHH72-Fc constructs to cell surface expressed SARS-CoV-2 spike determined by flow cytometry. The graph shows the mean (n=2) ratio of the MFI of transfected (GFP^+^) cells over the MFI of non-transfected (GFP^-^) cells. **d.** Inhibition of ACE-2/RBD interaction determined by AlphaLISA (amplified luminescent proximity homogeneous assay). Biotinylated SARS-CoV-2 RBD was loaded on streptavidin coated Alpha Donor beads and human ACE-2-mFc protein was captured on anti-mouse IgG acceptor beads. Interference of the donor-acceptor bead interaction was assessed for serial dilutions of the indicated VHH-Fc constructs. Graph pad Prism was used for curve fitting and IC_50_ determination of triplicate measurements. **e**. Dose-dependent inhibition of SARS-CoV-2 RBD binding to the surface of VeroE6 cells in the presence of the indicated VHH72-Fc constructs as determined by flow cytometry. The graph shows the mean (n=2 ± SD) percentage of cells that bind RBD. **f.** A SARS-CoV-2 plaque reduction neutralization assay was performed with 3 fold serial dilutions of the indicated VHH-Fc fusion constructs. Thirtysix hours after infection, the cells fixed with 3.7% paraformaldehyde and stained with 0.5% crystal violet. Data points in the graph represent the number of plaques and are representative of one experiment that was repeated once. **g-h**. humVHH_S56A/LALAPG-Fc and (humVHH)_2_/WT-Fc at 20 mg/kg protect hamsters against SARS-CoV-2 challenge. Hamsters were intraperitoneally injected with 20 mg/kg of palivizumab, humVHH_S56A/LALAPG-Fc or (humVHH)_2_/WT-Fc and challenged the next day with 2×10^6^ PFU of passage 6 BetaCov/Belgium/GHB-03021/2020. (**g**). Infectious virus in lungs and (**h**) viral RNA in lungs, ileum and stool determined on day 4 after the challenge. (**i**) Severity score of dilated bronchi on day 4 after the challenge.

As an alternative to the S56A point mutation to enhance potency, we further increased binding valency by grafting a tandem repeat of humanized VHH72, separated by a (G_4_S)_3_ linker, onto human IgG1 Fc, resulting in a tetravalent molecule. With this construct we observed a >100 fold higher affinity for SARS-CoV-2 RBD, >100-fold higher-affinity binding to spike-transfected HEK-cells, and a 5-10x stronger virus neutralizing activity compared with its bivalent counterpart (Figure 2f, Supplementary Fig. 6a-e, Supplementary Table 1, and Supplementary Table 2).

Taken together, our affinity improvement engineering campaign hence resulted in two ways of enhancing antiviral potency of our prototype pre-lead by about 5-10 fold each: S56A mutation in the VHH, or tandem repetition of the VHH. We next used the Syrian hamster SARS-CoV-2 challenge model to validate the protective potential of these potency-enhanced VHH-Fc fusion intermediate lead molecules resulting from both methods of affinity enhancement. This model mimics aspects of severe COVID-19 in humans, including high lung virus loads and the appearance of lung lesions^20, 53^. We used the BetaCov/Belgium/GHB-03021/2020 isolate, which is closely related to the first SARS-CoV-2 viruses isolated in Wuhan in the beginning of the new coronavirus pandemic^54^. We assessed the protective potential of a prophylactic dose at 20 mg/kg of either humVHH_S56A/LALAPG-Fc (PRNT_50_ = 0.13 µg/ml) and tetravalent (humWT-VHH72)_2_/WT-Fc (PRNT_50_ = 0.10 µg/ml) formats, administered one day prior to challenge (Fig. 2g-i). No infectious virus was detectable in lung homogenates from any of the VHH72-Fc treated hamsters except for one outlier in the (humWT)_2_/WT-Fc group (Fig. 2g). Compared with control treated animals, a statistically significant reduction in viral RNA levels in the lungs (> 4 log) and ileum (2 log) was observed on day 4 after infection in both VHH72-Fc treated groups, and in stool samples also for humVHH_S56A/LALAPG-Fc (Fig. 2h). Protection was also evident based on µ-computer tomography (µCT) imaging of the lungs on day 4, which showed a significantly reduced incidence of dilated bronchi in the VHH72-Fc treated animals (Fig. 2i). As both methods of affinity enhancement worked well in this experiment, both the bivalent and tetravalent designs were propagated in the subsequent stage of the drug development campaign.

### Generation of a lead therapeutic

Robust expression levels, chemical and physical stability, and absence of atypical posttranslational modifications are important prerequisites for the “developability” of a biologic^55^. Finalizing the design of our molecules for optimal homogeneity, we first truncated the upper hinge of the Fc, in common with most Fc fusions, which avoids heterogeneity due to an unpaired cysteine residue, and compensated for the reduced spacing between the VHH and the Fc by extending the glycine-serine linker to 10 amino acids: (G_4_S)_2_. Next, analysis by mass spectrometry had revealed partial C-terminal lysine removal in CHO-produced product, and we removed this source of charge heterogeneity by removing this lysine codon from the expression construct (Supplementary Fig. 7). Third, to avoid all possibility of N-terminal pyroglutamate formation and its associated charge heterogeneity, we decided to substitute the native N-terminal glutamine residue with an aspartic acid rather than glutamic acid. Analysis of the final designs by intact (reduced) protein and peptide-level LC-MS/MS showed that the VHH/linker/hinge moieties were homogenous and the Fc had CHO-typical N-glycans (Supplementary Fig. 8) and only well-known chemical modifications that are typical for lab-scale transient transfection produced antibodies: low-level glycation at 2 lysine residues (Supplementary Fig. 7 and Supplementary Table 3) and a deamidation site (Supplementary Fig. 9; Supplementary Table 3). Such chemical modifications are limited and controlled for during antibody manufacturing.

Irrespective of the length of the glycine-serine linkers, valency, and Fc types, the VHH72-Fc variants were efficiently produced in transiently transfected ExpiCHO cells with yields as high as 1.2 g/l cell culture after purification (Supplementary Table 4). The resulting humVHH_S56A/LALAPG-Fc/Gen2 construct displayed similar RBD-binding kinetics and SARS-CoV-2 neutralizing activity as humVHH_S56A/LALAPG-Fc (Supplementary Table 1, and Supplementary Fig. 6e). Combining our two potency-enhancing modalities, we introduced the S56A mutation in each of the four VHH72 moieties in the tetravalent format in construct (humVHH_S56A)_2_/LALAPG-Fc/Gen2, which further increased the *in vitro* antiviral potency, reaching a PRNT_50_ value of 0.02 µg/ml, *i.e.* 50-fold improved over the pre-lead parental construct (Supplementary Fig. 6e and Supplementary Table 1). Finally, we also generated hIgG1 LALA variants of the final bivalent sequence design, with or without the S56A VHH mutation (Supplementary Table 1).

All constructs throughout this lead development campaign displayed excellent biophysical properties. Even without any formulation optimization, Dynamic Light Scattering (DLS) analysis of WT-VHH/WT-Fc, for example, at concentrations up to 30 mg/ml in PBS revealed a homogenous sample composition, with only minor levels of aggregates below the limit of quantification (Fig. 3a). All VHH72-Fc constructs displayed a very low tendency to multimerize or aggregate, with >97% of purified VHH72-Fc variant molecules occurring in a SEC-MALS peak with a calculated molar mass of 77-79 kDa, corresponding to a VHH72-Fc monomer (Fig. 3b and Supplementary Table 5), and >95% remaining as such even after 10 days of thermal stress at 40°C (Fig. 3d and Supplementary Table 6). Thermal stability as measured by protein unfolding and aggregation was also not affected by the humanization, S56A, or LALA(PG) substitutions. Unfolding and aggregation of the tetravalent formats (humVHH)_2_/WT-Fc and (humVHH_S56A)_2_/LALAPG-Fc/Gen2, however, started at lower temperatures than the bivalent VHH-Fc variants, whereas aggregation of monomeric VHHs remained very low even at high temperatures (Fig. 3c, Supplementary Fig. 10, and Supplementary Table 6). Prolonged storage at 40°C of constructs humVHH_S56A/LALAPG-Fc/Gen2, humVHH_S56A/LALA-Fc/Gen2, (humVHH_S56A)_2_/LALAPG-Fc/Gen2, and humVHH/LALA-Fc/Gen2 demonstrated high stability with a homogeneous composition of the bivalent constructs, and a minor tendency of tetravalent construct (humVHH_S56A)_2_/LALAPG-Fc/Gen2 to multimerize after 10 days (Fig. 3d-e and Supplementary Table 6). Additionally, in a polyethylene glycol (PEG) precipitation assay which mimics high concentration solubility, the tetravalent molecule displayed less solubility with increasing PEG concentrations (lower PEG midpoint; Supplementary Fig. 11a). Increased apparent hydrophobicity (greater retention time) on a HIC column was also observed for the tetravalent molecule compared to the bivalent molecule indicating a greater probability of aggregation (Supplementary Fig. 11b). The tetravalent molecule also had a more acidic pI that would be potentially less favorable for a manufacturing strategy, requiring greater optimization of ion exchange steps (Supplementary Table 7)^56^.

**Figure 3.**
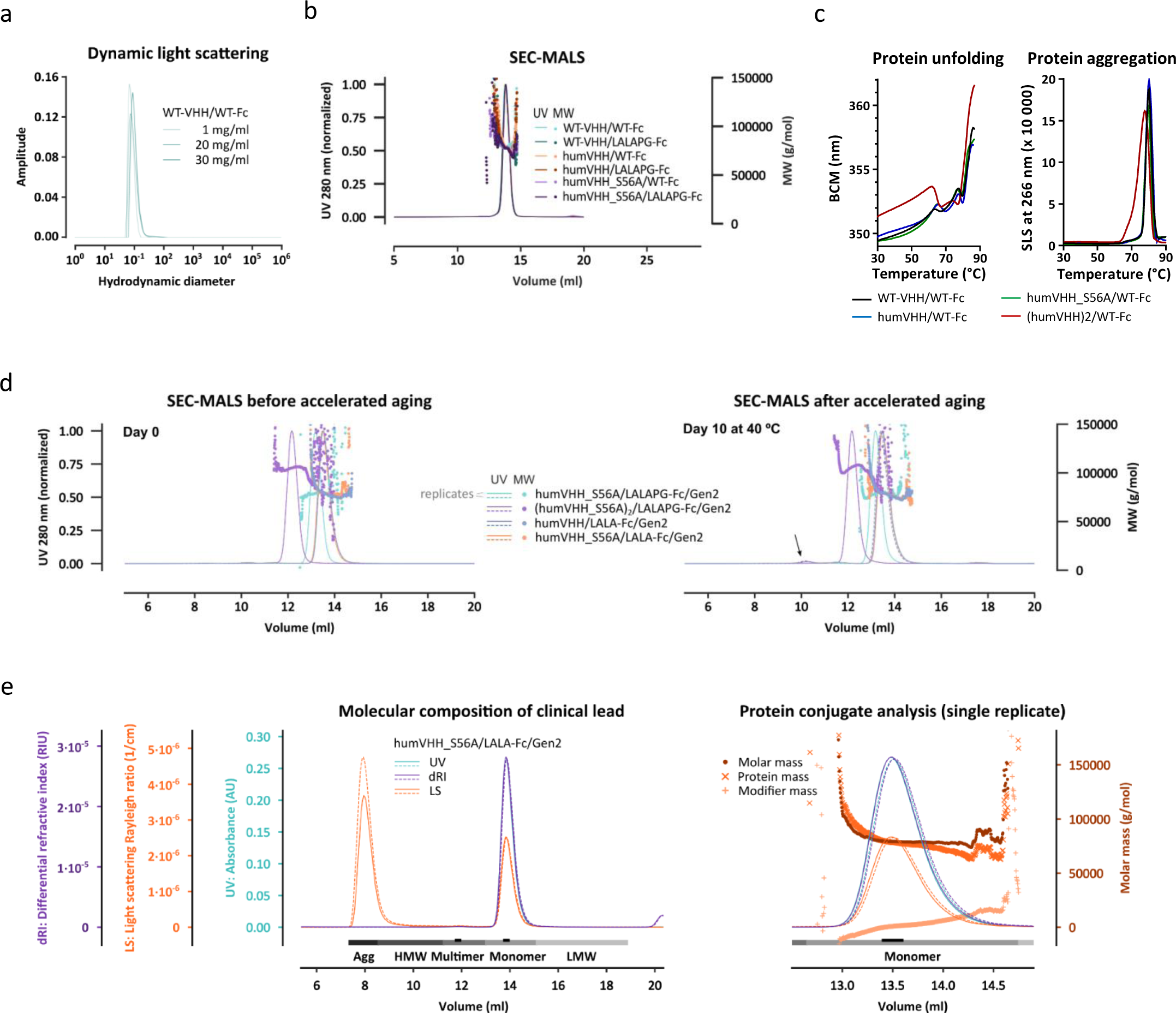
VHH72-Fc constructs have excellent biophysical properties. **a.** DLS of WT-VHH/WT-Fc at 25°C, at 1.0, 20 or 30 mg/ml in 25 mM His and 125 mM NaCl, pH6.0. **b.** SEC-MALS of 2-4 mg/ml samples of the prototype WT-VHH/WT-Fc and variants in which humanization, affinity-increasing (S56A) and LALAPG mutations were introduced (in 25 mM His and 125 mM NaCl, pH6.0). MW = molar mass of peak fraction as determined by light scattering. **c.** Thermostability of the indicated VHH72-Fc constructs. The left-hand graph shows protein unfolding during a thermal ramp measured by intrinsic Trp fluorescence and expressed as barycentric mean (BCM) of the fluorescence intensity at 300 and 400 nm. The right-hand graph shows protein aggregation during a thermal ramp measured by SLS at 260 nm (see also Supplementary Figure 12) **d.** Ten day-storage at 40°C causes no major changes in SEC-MALS profiles of duplicate 1 mg/ml VHH72-Fc variant samples in PBS. MW = molar mass of peak fraction as determined by light scattering. **e.** Complete SEC-MALS profile of clinical lead molecule humVHH_S56A/LALA-Fc/Gen2 run in duplicate (solid and dashed lines). Complete peaks are indicated in dark grey for quantitative analysis, peak apex indicated in light grey for qualitative analysis (results in Supplementary Table 6). The protein conjugate analysis was performed based on the differential extinction coefficients and refractive index values of proteins versus conjugated glycan modifiers.

### VHH72_S56A-Fc constructs protect therapeutically against SARS-CoV-2 challenge infection in hamsters

As a next step we analyzed the *in vivo* efficacy of the S56A-containing fully sequence-optimized bivalent (humVHH_S56A/LALAPG-Fc/Gen2, PRNT IC_50_ = 0.11 µg/ml) and tetravalent ((humVHH_S56A)_2_/LALAPG-Fc/Gen2, PRNT IC_50_ = 0.02 µg/ml) VHH72-Fc constructs against SARS-CoV-2 strain BetaCoV/Munich/BavPat1/2020. This strain carries the D614G mutation in the spike protein, which provides an advantage in fast viral entry and currently dominates the pandemic^57^. The experiments were performed by Viroclinics CRO under the cGLP quality system. Doses of 20, 7 and 2 mg/kg of humVHH_S56A/LALAPG-Fc/Gen2 or (humVHH_S56A)_2_/LALAPG-Fc/Gen2 were administered intraperitoneally 4 hours after challenge, a time point where lung viral titers were already increasing, taken into consideration that the intraperitoneal administration route gives a delay in serum exposure in the first 8 hours. Control animals received 20 mg/kg of palivizumab, a humanized monoclonal antibody directed against the fusion protein of human respiratory syncytial virus, and one group of animals received 20 mg/kg of the bivalent construct one day prior to challenge, as a bridge from our previous experiment using a D614 early pandemic virus, in which we had shown that this dosage was effective in this prophylactic setup (see Fig. 2g-i). Virus replication in the lungs was completely blocked in all but one animal receiving the prophylactic bivalent 20 mg/kg as well as in the therapeutic 20 and 7 mg/kg groups, although 2 out of 6 animals in the therapeutic tetravalent 20 mg/kg group showed residual virus titers (Fig. 4a,b). A clear dose-response relationship was observed, with increased variability in responses at the 2 mg/kg doses (Fig. 4a,b). Gross lung pathology was lowest in the animals that had been treated with 7 mg/kg of the bivalent construct (Fig. 4c). Viral loads in the bronchoalveolar lavage fluid of the hamsters were reduced to background levels in all but 2 animals that had received humVHH_S56A/LALAPG-Fc/Gen2 or (humVHH_S56A)_2_/LALAPG-Fc/Gen2 (Supplementary Fig. 12a). Interestingly, viral titers in the nose and throat of the challenged hamsters were also significantly and dose-dependently reduced compared with the palivizumab control group, indicating that parenteral, post-challenge administration of the VHH-Fc constructs restricts viral replication in the upper and lower respiratory tract of the hamsters (Supplementary Fig. 12b,c).

**Figure 4.**
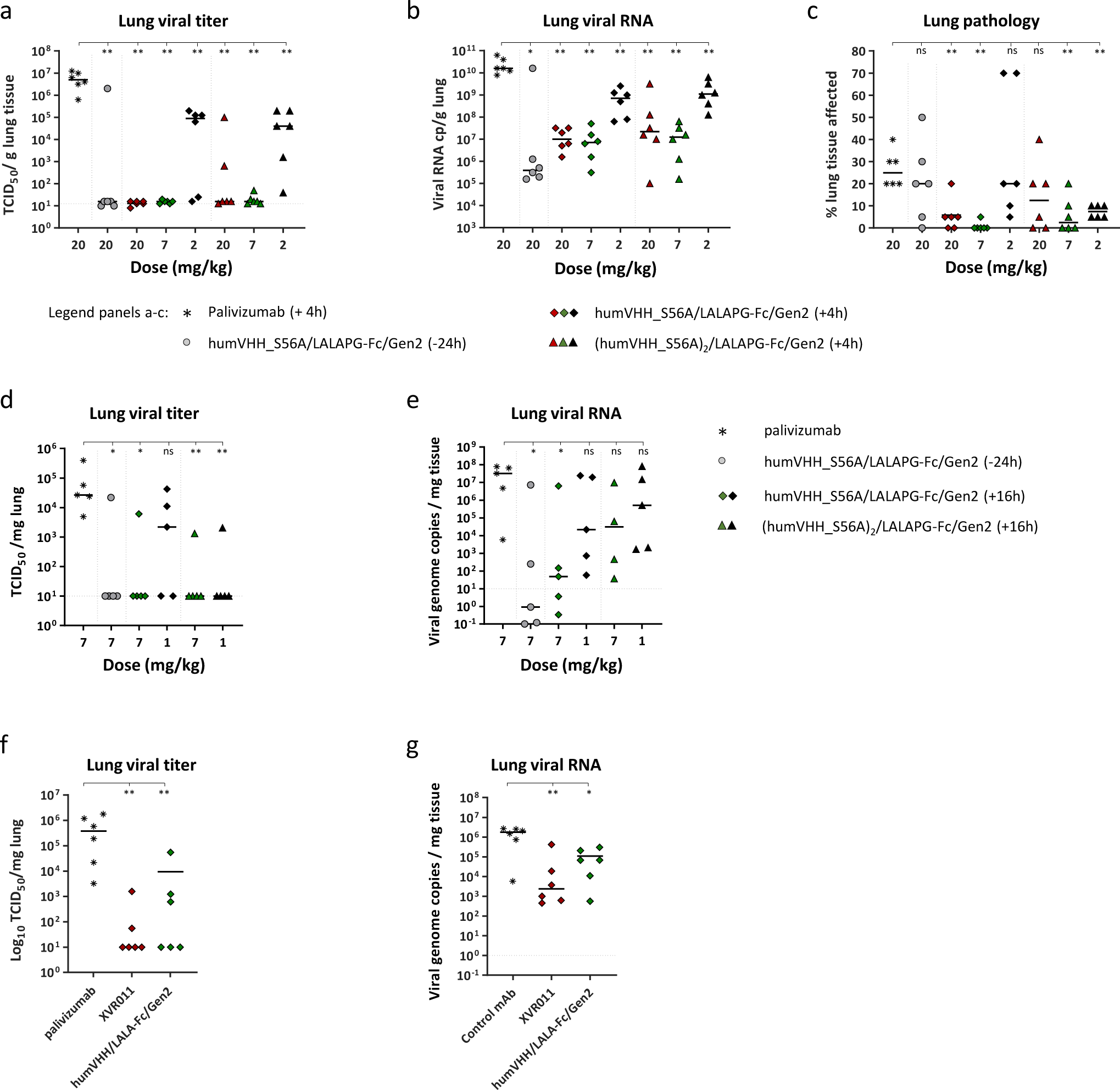
Therapeutic administration of VHH72-Fc constructs restricts SARS-CoV-2 virus replication in Syrian hamsters. **a-c.** Hamsters were challenged with 1×10^4^ PFU of BetaCoV/Munich/BavPat1/2020 and 4 hours later injected intraperitoneally with 20, 7 or 2 mg/kg of bivalent humVHH_S56A/LALAPG-Fc/Gen2 or tetravalent (humVHH_S56A)_2_/LALAPG-Fc/Gen2. The negative control group was treated with 20 mg/kg of palivizumab, injected 4 hours after the challenge infection; hamsters in a prophylactic control group received 20 mg/kg of humVHH_S56A/LALAPG-Fc/Gen2 one day before the challenge. (**a**) Lung virus loads, (**b**) lung viral RNA copies, and (**c**) gross lung pathology determined on day 4 after infection. **d,e**. Hamsters received an intraperitoneal injection of 7 mg/kg of humVHH_S56A/LALAPG-Fc/Gen2 one day prior to challenge or were treated by intraperitoneal injection of 1 or 7 mg/kg of humVHH_S56A/LALAPG-Fc/Gen2 or (humVHH_S56A)_2_/LALAPG-Fc/Gen2 19h after infection with 2×10^6^ PFU of passage 6 BetaCov/Belgium/GHB-03021/2020. Seven mg/kg of palivizumab was used as a negative control treatment. (**d**) Virus load and (**e**) viral RNA levels in the lungs on day 4 after challenge. **f,g**. Hamsters were treated with 4mg/kg of palivizumab, humVHH_S56A/LALA-Fc/Gen2 or humVHH/LALA-Fc/Gen2 injected intraperitoneally 24h after challenge with 2×10^6^ PFU of passage 6 BetaCov/Belgium/GHB-03021/2020. Viral RNA and infectious virus were determined in lung tissue on day 4 after infection. Data were analyzed with the Mann-Whitney U-test using GraphPad Prism software. *, p< 0.05; **, p<0.01; ***, p<0.001.

In a separate experiment, we independently validated this therapeutic dose finding for the bivalent and tetravalent VHH-Fc fusion lead constructs at 7 and 1 mg/kg at a different laboratory (Neyts lab) by intraperitoneal injection, 16h after challenge with a different SARS-CoV-2 isolate (BetaCov/Belgium/GHB-03021/2020 strain) (Fig. 4d). As a prophylactic control, the bivalent humVHH_S56A/LALAPG-Fc/Gen2 construct was administered 1 day prior to challenge at 7 mg/kg. In this study, the infectious virus load in the lungs was also here significantly reduced compared with the control treated animals for both the humVHH_S56A/LALAPG-Fc/Gen2 and (humVHH_S56A)_2_/LALAPG-Fc/Gen2 treated groups at 7 mg/kg, but not at the 1 mg/kg treatment with the bivalent construct. In addition, a strong reduction in genomic viral RNA levels was observed for the 7 mg/kg dose of the bivalent construct in both the therapeutic and prophylactic setting, with the other groups showing higher variability (Fig. 4e).

Finally, we conducted a hamster challenge experiment to evaluate the *in vivo* impact of the S56A mutation in VHH72 in the context of humVHH_S56A/LALA-Fc/Gen2, administered at a dose of 4 mg/kg 24h after challenge with the BetaCov/Belgium/GHB-03021/2020 strain. Treatment with humVHH_S56A/LALA-Fc/Gen2 resulted in a stronger reduction in infectious virus and viral genomic RNA in the lungs compared with humVHH/LALA-Fc/Gen2, providing evidence that the increased *in vitro* affinity and antiviral potency of the S56A mutation in VHH72-Fc also resulted in enhanced protection *in vivo* (Fig. 4f,g). We also determined the concentration of SARS-CoV-2 RBD IgG1-Fc antibodies in hamster serum sampled on day 4 after viral challenge and found a clear inverse correlation between the concentration of SARS-CoV-2 RBD-binding antibodies in serum on day 4 after the challenge and the infectious virus and viral RNA loads in the lungs of the challenged hamsters (Supplementary Fig. 12d,e). Collectively, the results of these stringent hamster challenge experiments executed at two institutions with two distinct isolates, strongly indicate that prophylactic as well as therapeutic administration of humVHH_S56A/LALA-Fc/Gen2 can fully control viral replication at a 4-7 mg/kg dose. At this point, we considered whether to choose the bivalent or tetravalent molecule for further development. While *in vitro*, the tetravalent molecule is clearly more potent, this difference is not apparent *in vivo*. Tetravalency moreover came at a cost of a more complex molecule to manufacture and stably formulate, as described above. We hence prioritized the simpler bivalent design and selected humVHH_S56A/LALA-Fc/Gen2 as our clinical lead molecule, which was termed XVR011.

### XVR011 lacks off-target binding to human proteins and has reduced FcγR binding

For safe use in humans upon systemic administration, antibodies have to have a low propensity for off-target binding to other human membrane/extracellular proteins. To evaluate this for XVR011, we used a human membrane protein microarray assay in which reactivity was probed against fixed HEK293 cells that each overexpress one of 5475 full-length human plasma membrane proteins and cell surface-tethered human secreted proteins and a further 371 human heterodimeric such proteins^58^. Only four proteins were found to potentially show some binding in a high-sensitivity primary screen using XVR011 as primary antibody and an anti-human IgG1 secondary detection antibody: over-expressed proteins FCGR1A, IGHG3, IGF1, and CALHM6, next to the spotted SARS-CoV2 Spike protein, the primary target and positive control. In a targeted confirmation experiment to study binding to these proteins in more detail, we used the clinically well-validated rituximab (anti-CD20) and cells expressing its antigen as a control for potential hIgG1 Fc-mediated interactions, as well as a buffer control instead of primary antibody, to check for interactions with the secondary detection antibody (Supplementary Fig. 13). Apart from the expected XVR011 reactivity with recombinant SARS-CoV-2 spike protein, binding to the primary hits was as low or lower as that of rituximab (which bound CD20-expressing cells) and a robust detection signal was only observed for fixed cells that expressed human immunoglobulin heavy gamma-3 chain IGHG3. However, this reactivity was equally strong with the PBS and rituximab controls, and, therefore due to direct binding of the secondary detection antibody. Based on these results, we conclude that XVR011 is very specific and not polyreactive to human proteins at all, which supports its potential for safe use as a medicine.

Further validating the built-in safety feature, we verified that the introduced LALA mutations in the context of VHH-Fc fusion construct XVR011 resulted in reduced binding to activating Fcγ receptors. We used surface plasmon resonance (SPR) to determine the binding affinity of XVR011 with activating receptors FcγRI, FcγRIIa (H167), FcγRIIa (R167), FcγRIIIa (V176), FcγRIIIa (F176), and FcγRIIIb and the inhibitory receptor FcγRIIb. Compared to rituximab, XVR011 bound to immobilized FcγRI and FcγRIIIa (V176) with a 2300- and 40-fold lower affinity whereas binding to FcγRIIa (H167), FcγRIIa (R167), FcγRIIIa (F176), FcγRIIIb, and FcγRIIb was barely detectible and occurred with an estimated equilibrium dissociation constant (KD) of more than 20 µM (Supplementary Fig. 14 and Supplementary Table 8).

### HumVHH_S56A/LALA-Fc/Gen2 binds to the RBD of a broad range of Sarbecoviruses and neutralizes circulating SARS-CoV-2 variants

To test the Sarbecovirus binding breadth of humVHH_S56A/LALA-Fc/Gen2, we applied a flow cytometry-based yeast surface display method^38^. HumVHH_S56A/LALA-Fc/Gen2 bound to yeast surface displayed RBD of SARS-CoV-2, GD-Pangolin, RaTG13, SARS-CoV-1, LYRa11, and WIV1 (all clade 1) with high affinity. In addition, humVHH_S56A/LALA-Fc/Gen2 could bind the RBD of Rp3 and HKU3-1 (both clade 2) and BM48-31 (clade 3) whereas CB6^50^ and S309^59^, two antibodies also under advanced clinical development, did not bind to these RBDs (Supplementary Fig. 15). Binding of any of the antibodies to the RBDs of the clade 2 viruses Rf1, ZC45 and ZXC42 was undetectable.

**Figure 5.**
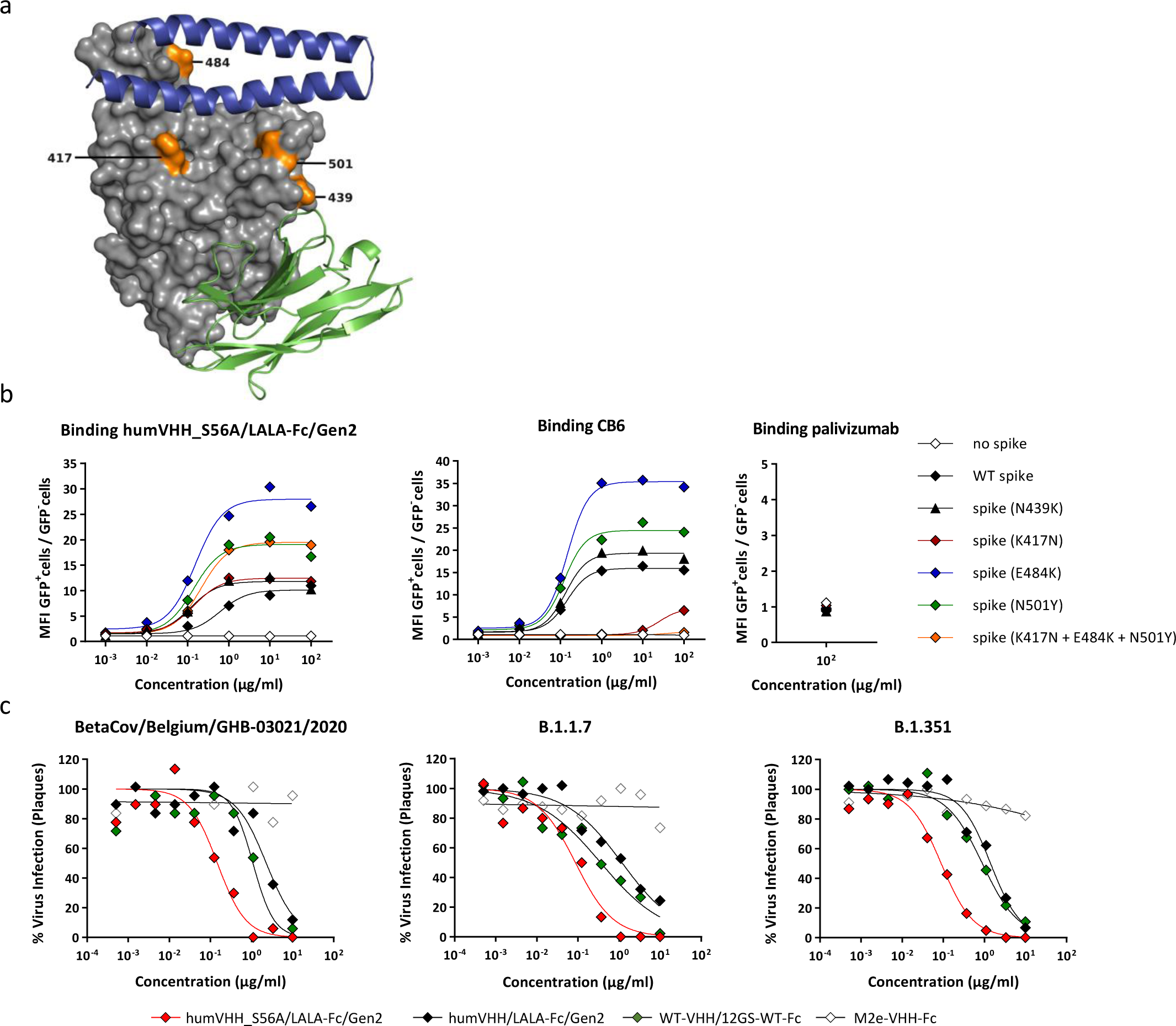
humVHH_S56A/LALA-Fc/Gen2 neutralizes SARS-CoV-2 variants of concern. **a**. Surface view of SARS-CoV-2 RBD (grey) with VHH72 (green cartoon, bottom) and the N-terminal helixes of ACE2 (blue cartoon, top). The RBD-residues K417, N439, E484 and N501 (orange) are indicated. **b.** Binding of humVHH_S56A/LALA-Fc/Gen2 (left), CB6 (middle) and palivizumab (right) to SARS-CoV-1 spike with the RBD replaced by WT, N439K, K417N, E484K, N501Y or (K417N + E484K + N501Y) RBD of SARS-CoV-2, expressed on the surface of 293T cells. Data points represent the ratio of the mean fluorescence intensity (MFI) of transfected (GFP^+^) cells over the MFI of non-transfected (GFP^-^) cells, as determined by flow cytometry. **c**. SARS-CoV-2 plaque reduction neutralization assay with 3 fold serial dilutions of the indicated VHH-Fc fusion constructs using BetaCov/Belgium/GHB-03021/2020, B1.1.7, or B.1.351 variant viruses.

All RBD mutations observed in the current rapidly spreading VOCs are distant from the VHH72 contact region (Fig. 5a). We evaluated binding affinity of our optimized drug lead in a flow cytometry assay, showing equally strong binding of humVHH_S56A/LALA-Fc/Gen2 to wild type as to SARS-CoV-2 RBD N501Y, K417N, E484K, and K417N + E484K + N501Y SARS-CoV-2 RBD mutants expressed on the surface of mammalian cells in the context of the complete spike of SARS-CoV-1 (Fig. 5b)^18^. N501Y is the mutation seen in both the B.1.1.7.^60^ and B.1.351^61^ variants, in which the K417N and E484K are combined with it. Binding of humVHH_S56A/LALA-Fc/Gen2 to RBD variant N439K^62^, in the periphery of the VHH72 contact region and observed 8661 times as of February 18 2021, was also not affected (Fig. 5b). This is unsurprising as this transition to the positively charged Lys mimics the Arg which is present at this position in SARS-CoV-1, to which VHH72 was raised. Fully in line with these results, humVHH_S56A/LALA-Fc/Gen2 showed equal neutralization potency against authentic SARS-CoV-2 BetaCov/Belgium/GHB-03021/2020, a B.1.1.7. variant with N501Y and a B.1.351 variant carrying K417N, E484K, and N501Y mutations (Fig. 5c).

## Discussion

There is an urgent need for safe and effective anti-SARS-CoV-2 drugs that can prevent or cure COVID-19. We report the protein engineering-based drug development of a potent cross-neutralizing, VoC-resistant anti-COVID-19 biologic based on a humanized, *in silico* affinity enhanced VHH72 variant, which, in bivalent and tetravalent VHH-Fc format, demonstrated strong anti-viral efficacy in the hamster challenge model. Compared with conventional human monoclonal antibodies, the VHH-Fc fusion construct is smaller in size (80 kDa versus 150 kDa) and encoded by a single gene, giving advantages with respect to dosing and manufacturability.

Hundreds of human IgGs have so far been described from convalescent SARS-CoV-2 infected patients. Most of the neutralizing antibodies that are being assessed in clinical trials have been selected from such convalescent repertoires. However, the vast majority of neutralizing antibodies in convalescent plasma are targeting the RBM in the RBD and sites on the N-terminal domain (NTD), which are hence experiencing the majority of antibody-mediated selective pressure^15^. Not surprisingly, such antibodies are now affected by mutations in emerging new SARS-CoV-2 VOCs such as in strains classified in the lineages B.1.1.7, B.1.351^63^, and P.1^64^, with the latter two allowing escape from naturally acquired or spike vaccine-induced antibody neutralization and possibly may have arisen in this way. Efforts to develop a pan-Sarbecovirus-neutralizing antibody, such as the VHH72-based biologic described here, are therefore warranted to help protect the society against disease caused by the current, antigenically drifting, SARS-CoV-2 pandemic virus and potentially against future SARS-like coronavirus outbreaks. The VHH72 discovery, determination of its complex structure bound to the SARS-CoV-1 RBD^17^ and demonstration of its antiviral potency as Fc-fusion was first to establish that antibody binding to its non-RBM epitope region of the RBD, while at the same time also occluding ACE2 binding to the RBM, leads to potent neutralization of sarbecoviruses such as SARS-CoV-1 and −2. The functional constraints in the VHH72 binding region (inter-RBD interactions in prefusion spike, interactions with the helix-loop-helix in S2) can explain its strong conservation across the Sarbecoviruses. Meanwhile, a few other mAbs and VHHs which also bind to the VHH72 core epitope (Y369, F377, K378) have been reported^22, 25, 36, 37, 65–67^, with anti-SARS-CoV-2 potency correlating with strength of competition with ACE2 binding. The much lower immunogenicity of the VHH72 binding region in humans than the epitopes on the RBM is also consistent with a recent large-scale serological survey of convalescent SARS-CoV-2 patients, which demonstrated that mAbs with epitopes strongly overlapping that of VHH72 (site II in that study) were competed against much less potently and were present in a smaller proportion of patient sera than was the case for mAbs targeting the ACE2-binding region of the RBD^22^. These results were very similar to what was observed for S309, which is also a SARS-CoV-1/2 cross-neutralizing antibody presently in clinical development^59^. Amongst these binding agents presently described against this ‘cryptic supersite’, XVR011 best combines a very strong potency both *in vitro* and *in vivo*, SARS-CoV-1/2 cross-neutralization, very broad cross-clade sarbecovirus binding, and unaltered strong potency against the currently emerging VoCs, warranting its clinical development. For this, we considered it prudent in patients with progressing COVID-19 disease to mainly rely on a pure virus neutralization mechanism of action, and thus to suppress Fcγ receptor binding of the antibody’s Fc domain. Interestingly, antibodies that bind to VHH72-epitope overlapping epitopes, by nature already have very low complement-dependent cytotoxicity because this RBD region is not sterically accessible in the dominant closed spike conformation of the native spike protein^22^. To silence antibody Fc-mediated effector functions, we settled on the IgG LALA-Fc mutations, which are amongst the best-validated for this purpose^52^. While the field of SARS-CoV-2 antibody development currently sees arguments in both directions with regard to the desired extent of Fc effector functionality in diverse settings (prophylactic, therapeutic, route and frequency of administration), our LALA-Fc choice attempts to strike an optimal balance between safety and efficacy in diverse application modalities, within the limitations of relevant current clinical data, while ascertaining an established regulatory track record and freedom to operate.

It was recently reported that in a therapeutic setting, some human neutralizing antibodies require intact Fc effector functions to control SARS-CoV-2 replication in the K18-hACE2 transgenic mouse and Syrian hamster challenge models^68^, using the LALAPG mutations to remove effector function of those antibodies. In contrast to that report, we found that Fc effector silent humVHH_S56A/LALA(PG)-Fc/Gen2 administered 4, 16 or 24 hours after viral challenge resulted in a very strong reduction of lung viral RNA and infectious virus levels. This suggests that the non-RBM binding mode of neutralization, likely including spike destabilization, perhaps combined with a faster biodistribution compared with a conventional antibody, allows for efficacious viral control. The requirement of effector functionality for therapeutic efficacy appears to be very antibody-dependent even with human antibodies, as a very recent study also demonstrated therapeutic efficacy in the same hamster model of other RBM-binding LALAPG-modified antibodies^69^.

The pharmaceutically fully developed VHH72-based biologic named XVR011, that combines potent neutralizing activity with high stability, broad coverage and silenced Fc effector functionality for enhanced safety, has currently completed cGMP-manufacturing and formal preclinical development. Clinical studies are now being started to evaluate safety and efficacy of rapid administration upon hospitalization of patients, within the first week of COVID-19 symptoms.

## Acknowledgements

We thank Gert Zimmer, Stefan Pöhlmann and Markus Hoffmann for providing reagents to generate VSV pseudotype particles, Jason McLellan and Daniel Wrapp for providing SARS-CoV-2 spike and RBD proteins, and expression plasmids for these proteins, Jesse Bloom for providing the *S. cerevisiae* RBD display library, and Vincent Munster and Michael Letko for providing chimeric SARS-CoV-1 spike expression constructs. We thank Nadja van Boxel and Joana Reis Pedro for excellent technical assistance in the generation of humanized VHHs, stability and functional characterization, and PK assessments. We thank Carolien De Keyzer, Lindsey Bervoets, Elke Maas, Thibault Francken, Tina Van Buyten, Jasper Rymenants and Kathleen Van den Eynde for excellent technical assistance in the hamster challenge studies. We thank Piet Maes for kindly providing the Belgian SARS-CoV-2 strain and Valentijn Vergote and Elisabeth Heylen for facilitating the animal studies. We are grateful to Hanna Hailu and Sarfaraj Topia for material generation, purification and distribution. We thank Oliver Zaccheo and Bruce Carrington for helpful advice in the conducting of the FcγR binding studies. We thank the VIB Protein Service Facility and Savvas Savvides for making MALS and BLI equipment available. We thank the staff of the VIB Flow Core Ghent for their support with flow cytometry experiments. We acknowledge all GISAID contributors for their sharing of sequencing data. We thank the entire teams at VIB-Center for Medical Biotechnology, VIB Innovation and Business, VIB Discovery Sciences and UCB for support and for coping with the effects of prioritization of this Covid19 emergency project. X.Z. received funding of the China Scholarship Council (grant No.201906170033). S.J. is supported by a PhD fellowship of the Fund for Scientific Research Flanders (FWO 1S21918N). This work was supported by FWO projects VIREOS, G0G4920N and G0B1917N, UGent GOA and BOF projects to X.S. and N.C., by VUB grant OZR3571 to N.D., and by the Bundesministerium fuer Bildung und Forschung (BMBF) through the Deutsches Zentrum für Luft- und Raumfahrt (DLR, grant number 01KI2077) to M.S. and P.S.

## Author contributions

B.S., L.v.S., K.R., W.v.B., W.W., S.D., S.D.C., S.V., C.L., H.E., J.B., A.F.O., J.P.C., S.C., D.D.V., G.M., J.C.Z.M., G.M., I.R., K.S., A.V.H., D.V.H., X.Z., L.L., S.J., S.t.H., L.S., L.L., R.B., J.B., D.S., P.S., A.O., I.Rem., C.F., R.A., S.J.F.K., M.E., and J.H. performed wet lab experiments. W.N., F.P., and G.M. performed computational modeling. D.F. compared and interpreted genomic RNA sequences of circulating SARS-CoV-2 viruses. B.S., L.v.S., N.C., and X.S. wrote the manuscript. All authors reviewed and edited the manuscript. H.J.T., K.D., S.J.F.K., D.J., J.R.-P, G.V.V., B.W., L.D., M.S., N.D., D.S., A.H., A.P., K.B., A.T., B.D., C.S., J.N., N.C., and X.S. supervised the work. Funding Acquisition: L.D., M.S., P.S., J.N., N.C., and X.S.

## Methods

### Molecular modeling of the VHH-72 (mutant) interaction with SARS-CoV-2 RBD

Molecular Dynamics simulations were with model-complexes of VHH72 (chain C from pdb-entry 6WAQ) and variants, with the outward-positioned RBD from the cryo-EM structure pdb-entry 6VSB of the SARS-CoV-2 prefusion spike glycoprotein (chain A, residues 335-528). The missing loops at residues 444-448, 455-490 and 501-502 in the cryo-EM RBD were reconstructed from the I-TASSER SARS-CoV-2 RBD model^39^ and the missing residues were added by the use of Swiss-PDBViewer^40^. Simulations of the VHH72(mutant)-RBD complex were with Gromacs version 2020.1^41^ using the Amber ff99SB-ILDN force field^70^ and were run for 5 nanoseconds. After conversion of the trajectory to PDB-format, snapshots were extracted every nanoseconds and were submitted to the FastContact 2.0 server^35^ for binding energy calculations.

### SARS-CoV-2 spike sequence variant analysis

SARS-CoV-2 genome sequences originating from infected human hosts were downloaded from GISAID (N= 571,131 genomes available on February 18 2021). Genomes with invalid DNA character code were removed. Spike coding sequences were retrieved by aligning the genomes to the reference spike sequence annotated in NC_045512.2 (Wuhan-Hu-1 isolate, NCBI RefSeq). For this purpose, pairwise alignments were performed using the R package Biostrings version 2.54.0, a fixed substitution matrix in the “overlap” mode with the following parameters according to Biostrings documentation: 1 and −3 for match and mismatch substitution scores; 5 as gap opening and 2 as gap extension penalties. Incomplete genomes without spike coding sequences, or generating very short or no alignment were removed. Coding sequences with frame-disturbing deletions were also excluded and the remaining open reading frames were *in-silico* translated using Biostrings option to solve “fuzzy” codons containing undetermined nucleotide(s). In the next step, predicted spike protein sequences with undetermined amino acids (denoted as X), derived from poor sequencing results (Ns) were removed. Further, full-length sequences with a single stop codon or lacking a stop signal (due to a possible C-terminal extension) were retained, while proteins with premature stop codon(s) were excluded.

The resulting 440,769 quality-controlled spike protein sequences were aligned using the ClustalOmega algorithm and R package msa version 1.18.0 with default parameters and the BLOSUM65 substitution matrix. R packages seqinr 3.6-1 and BALCONY 0.2.10 were used to calculate amino acid frequencies for all mutations occurring in the dataset at least once. Major and minor allele frequencies and counts were assigned. Effects of individual mutations on RBD yeast-surface display expression (a correlate of fold stability) were derived from Starr *et al*^38^. Data collected for spike protein RBD (positions 333 – 516) was visualized using ggplot2 version 3.3.0.

### E. coli and Pichia pastoris strains

*Escherichia coli* (*E. coli*) MC1061 or DH5α were used for standard molecular biology manipulations. The *Pichia pastoris* (syn. *Komagataella phaffi*) NRRL-Y 11430 OCH1 function-inactivated strain used for VHH-Fc screening (*P. pastoris* OCH1) was obtained by CRISPR-Cas9 engineering^71^. As reported before, the inactivation of the α-1,6-mannosyltransferase encoded by *OCH1* results in secretion of more homogenously glycosylated protein carrying mainly Man8 glycan structure^72^.

### Modular generation of expression plasmids

The expression vectors for all the VHH72-XXX-hFc muteins were generated using an adapted version of the Yeast Modular Cloning toolkit based on Golden Gate assembly^73^. Briefly, coding sequences for the *S. cerevisiae* α-mating factor minus EA-repeats (P3a_ScMF-EAEAdeleted), SARS-VHH72 mutants (P3b_SARS_VHH72-xxx) and human IgG1 hinge-human IgG1 Fc (P4a_(GGGGS)_x2_hIgG1.Hinge-hIgG1.Fc) were codon optimized for expression in *P. pastoris* using the GeneArt (Thermo Fisher Scientific) or IDT proprietary algorithm and ordered as gBlocks at IDT (Integrated DNA Technologies BVBA, Leuven, Belgium). Each coding sequence was flanked by unique part-specific upstream and downstream *Bsa*I-generated overhangs. The gBlocks were inserted in a universal entry vector via *BsmB*I assembly which resulted in different “part” plasmids, containing a chloramphenicol resistance cassette. Plasmids parts were assembled to form expression plasmids (pX-VHH72-xxx-hIgGhinge-hIgGFc) via a Golden Gate *Bsa*I assembly. Each expression plasmid consists of the assembly of 9 parts: P1_ConLS, P2_pGAP, P3a_ScMF-EAEAdeleted, P3b_SARS_VHH72-xxx, P4a_(GGGGS)x2hIgG1.Hinge-hIgG1.Fc, P4b_AOX1tt, P5_ConR1, P6-7_Lox71-Zeo, and P8_AmpR-ColE1-Lox66. Selection was in LB supplemented with 50 µg/ml carbenicillin and 50 µg/ml Zeocin®. All the part and expression plasmids were sequence verified. Transformations of linearized expression plasmids (*Avr*II) were performed using the lithium acetate electroporation protocol as described^74^.

### Protein expression and purification

*P. pastoris* cultures were grown in liquid YPD (1% yeast extract, 2% peptone, 2% D-glucose) or on solid YPD-agar (1% yeast extract, 2% peptone, 2% D-glucose, 2% agar) and selected with 100 µg/ml Zeocin® (InvivoGen). For small scale expression screening, 2-3 single colonies of *P. pastoris* OCH1 transformed with pX-VHH72-xxx-hIgGhinge-hIgGFc were inoculated in 2 ml BMDY (1% yeast extract, 2% peptone, 100 mM KH_2_PO_4_/K_2_HPO_4_, 1.34% YNB, 2% D-glucose, pH 6) or BMGY (same composition but with 1% glycerol replacing the 2% D-glucose) in a 24 deep well block. After 50 hours of expression in a shaking incubator (28°C, 225 rpm), the medium was collected by centrifugation at 1.500 g, 4°C for 5 minutes. Protein expression levels were evaluated on Coomassie-stained SDS-PAGE of crude supernatant. Crude supernatant was used immediately for analytics purposes (biolayer interferometry and mass spectrometry, see below) or stored at −20°C.

VHH72-Fc constructs were produced in ExpiCHO cells (Thermo Fisher Scientific) by transient transfection of the respective expression plasmids. 25 ml cultures with 6 ×10^6^ cells/ml were grown at 37°C and 8% CO_2_ and transfected with 20 µg of pcDNA3.3-based plasmid DNA using ExpiFectamine CHO reagent. Twenty four hours after transfection, 150 µL of ExpiCHO enhancer and 4 mL of ExpiCHO feed was added to the cells, and the cells were further incubated at 32°C and 5% CO_2_. Transfected cells were fed a second time 5 days post-transfection. As soon as cell viability had dropped below 75%, the culture medium was harvested. The VHH72-Fc constructs were purified from the clarified culture medium using a 5 ml MabSelect SuRe column (GE Healthcare). After a wash step with McIlvaine buffer pH 7.2, bound proteins were eluted using McIlvaine buffer pH 3. The eluted protein-containing fractions were neutralized with a saturated Na_3_PO_4_ buffer. These neutralized fractions were then pooled, and loaded onto a HiPrep Desalting column for buffer exchange into storage buffer (25 mM L Histidine, 125 mM NaCl pH 6) or onto a HiLoad 16/600 Superdex 200 pg size-exclusion column (GE-Healthcare) calibrated with phosphate-buffered saline (PBS) or storage buffer. Where data are labeled as ‘XVR011’, protein material manufactured from a stable CHO cell line has been used in the experiments, of identical sequence as humVHH_S56A/LALA-Fc/Gen2 (which is the naming we use for the protein produced in ExpiCHO transient transfection).

Open reading frames corresponding to the light and heavy chains of the hIgG1 anti-SARS-CoV-2 antibody S309 were ordered synthetically at IDT. For both, an optimized Kozak sequence was added upstream of the start codon. The secretion signal of a *Mus musculus* Igκ chain was used to direct secretion of the S309 light chain and of a *Mus musculus* IgG heavy chain to direct secretion of the S309 heavy chain. The carboxy-terminal lysine residue of the S309 heavy chain was omitted. The synthetic DNA fragments were solubilized in ultraclean water at a concentration of 20 ng/µl, A-tailed using the NEBNext-dA-tailing module (NEB), purified using CleanPCR magnetic beads (CleanNA), and inserted in pcDNA3.3-TOPO vector (ThermoFisher). The ORF of positive clones was sequence verified and pDNA of selected clones was prepared using the NucleoBond Xtra Midi kit (Machery-Nagel). Expression in ExpiCHO cells and purification of S309 was performed as described above for the VHH72-Fc constructs except that for S309 the heavy chain and light chain encoding plasmids were mixed in a ratio of 1:2. CB6 (heavy chain GenBank MT470197, light chain GenBank MT470196) was custom produced in a mammalian cell system by Genscript.

### Enzyme-linked immunosorbent assay

Wells of microtiter plates (type II, F96 Maxisorp, Nuc) were coated overnight at 4°C with 100 ng of recombinant SARS-CoV S-2P protein (with foldon), SARS-CoV-1 S-2P protein (with foldon), Fc-tagged SARS-CoV-2 RBD-SD1 or BSA. The coated plates were blocked with 5% milk powder in PBS. Dilution series of the VHHs were added to the wells. Binding was detected by incubating the plates sequentially with either mouse anti-Histidine Tag antibody (MCA1396, Abd Serotec) followed by horseradish peroxidase (HRP)-linked anti-mouse IgG (1/2000, NXA931, GE Healthcare) or Streptavidin-HRP (554066, BD Biosciences) or by HRP-linked rabbit anti-human IgG (A8792, Sigma). After washing, 50 μL of TMB substrate (Tetramethylbenzidine, BD OptETA) was added to the plates and the reaction was stopped by addition of 50 μL of 1 M H_2_SO_4_. The absorbance at 450 nM was measured with an iMark Microplate Absorbance Reader (Bio Rad). Curve fitting was performed using nonlinear regression (Graphpad 8.0).

### Biolayer interferometry (BLI)

The SARS-CoV-2 RBD binding kinetics of VHH72-hIgG1Fc affinity optimized variants in *P. pastoris* supernatant were assessed via biolayer interferometry on an Octet RED96 system (FortéBio). Anti-mouse IgG Fc capture (AMC) biosensors (FortéBio) were soaked in kinetics buffer (10 mM HEPES pH 7.5, 150 mM NaCl, 1 mg/ml bovine serum albumin, 0.05% Tween-20 and 3 mM EDTA) for 20 min. Mouse IgG1 Fc fused SARS-CoV-2 RBD (Sino Biological, 40592-V05H) at 5-15 µg/ml was immobilized on these AMC biosensors to a signal of 0.3-0.8 nm. Recombinant protein concentrations in crude cell supernatants of VHH72-hFc expressing *P. pastoris* OCH1^-^ were estimated based on band intensity on Coomassie-stained SDS-PAGE as compared to a purified VHH-hFc protein. Crude supernatants were diluted 20 to 100-fold in kinetics buffer to an approximate VHH72-hFc affinity mutant concentration of 5-10 nM. Association was measured for 180 s and dissociation for 480 s in similarly diluted supernatant of a non-transformed *P. pastoris* culture. Between analyses, biosensors were regenerated by three times 20 s exposure to regeneration buffer (10 mM glycine pH 1.7). Using ForteBio Data Analysis 9.0 software, data were double reference-subtracted and the decrease of response signal during dissociation was determined to assess the off-rates of the VHH72-Fc variants. To measure the affinity of monovalent VHH72 variants for RBD, monomeric human Fc-fused SARS-CoV-2_RBD-SD1^42^ at 15 µg/ml was immobilized on anti-human IgG Fc capture (AHC) biosensors (FortéBio) to a signal of 0.35-0.5 nm. In an alternative setup, mouse IgG1 Fc fused SARS-CoV-1 RBD (Sino Biological, 40150-V05H) or SARS-CoV-2 RBD (Sino Biological, 40592-V05H) at 15 µg/ml was immobilized on AMC biosensors (ForteBio). Association (120 s) and dissociation (480 s) of two-fold dilution series starting from 200 nM VHH72 variant in kinetics buffer were measured. To assess full binding kinetics of purified bivalent VHH72-hFc variants, bivalent mouse IgG1 Fc fused SARS-CoV-2-RBD (Sino Biological, 40592-V05H) at 15 µg/ml was immobilized on AMC biosensors to a signal of 0.4-0.5 nm. Association (120 s) and dissociation (480 s) of twofold serial dilutions starting from 30 nM VHH72-hFc variants in kinetics buffer were measured. Between analyses of binding kinetics, AHC and AMC biosensors were regenerated by three times 20 s exposure to regeneration buffer (10 mM glycine pH 1.7). Data were double reference-subtracted and aligned to each other in Octet Data Analysis software v9.0 (FortéBio) based on a baseline measurement of a non-relevant VHH-IgG1 Fc fusion protein (for kinetics of VHH72-hFc variants) or kinetics buffer (for kinetics of monovalent VHHs). Association and dissociation of non-saturated curves (both 1:1 kinetics of monovalent VHHs and 2:2 kinetics of VHH-hFc variants) were fit in a global 1:1 model.

### Polyethylene glycol (PEG) aggregation assay

Stock PEG 3350 (Merck, 202444) solutions (w/v) were prepared in PBS pH 7.4 or 50 mM histidine, 250 mM proline pH 5.5. A 1:1.1 serial titration was performed by an Assist plus liquid handling robot (Integra, 4505), to avoid liquid handling issues the stock concentration of PEG 3350 was capped at 40%. To minimize non-equilibrium precipitation, sample preparation consisted of mixing protein and PEG solutions at a 1:1 volume ratio. 35 µL of the PEG 3350 stock solutions was added to a 96 well v bottom PCR plate (A1 to H1) by a liquid handling robot. 35 µL of a 2 mg/ml sample solution was added to the PEG stock solutions resulting in a 1 mg/ml test concentration. This solution was mixed by automated slow repeat pipetting, samples were then incubated at 20°C for 24 h. The sample plate was subsequently centrifuged at 4000 x g for 1 h at 20°C. 50 µL of supernatant was dispensed into a UV-Star®, half area, 96 well, μClear®, microplate (Greiner, 675801). Protein concentrations were determined by UV spectrophotometry at 280 nm using a FLUOstar Omega multi-detection microplate reader (BMG LABTECH). The resulting values were plotted using GraphPad Prism 7.04 and the PEG midpoint score was derived from the midpoint of the sigmoidal dose-response (variable slope) fit.

### Hydrophobic interaction chromatography (HIC) assay

Apparent hydrophobicity was assessed using a hydrophobic interaction chromatography (HIC) assay employing a Dionex ProPac HIC-10 column, 100 mm×4.6 mm (Thermo Fisher 063655), containing a stationary phase consisting of a mixed population of ethyl and amide functional groups bonded to silica. All separations were carried out on an Agilent 1200 HPLC equipped with a fluorescence detector. The column temperature was maintained at 20°C throughout the run and the flow rate was 0.8 ml/min. The mobile phases used for the HIC method were (A) 0.8 M ammonium sulfate and 50 mM phosphate pH 7.4, and (B) 50 mM phosphate pH 7.4. Following a 2 min hold at 0% B, the column was loaded with 15 µl of sample at 2 mg/ml, and bound protein was eluted using a linear gradient from 0 to 100% B in 45 min and the column was washed with 100% B for 2 min and re-equilibrated in 0% B for 10 min prior to the next sample. The separation was monitored by absorbance at 280 nm.

### Experimental isoelectric point (pI) measurement

An iCE3^TM^ whole-capillary imaged capillary isoelectric focusing (cIEF) system (ProteinSimple) was used to experimentally determine pI. Samples were prepared by mixing the following: 30 µl sample (from a 1 mg/ml stock in HPLC grade water), 35 µL of 1% methylcellulose solution (ProteinSimple, 101876), 4 µl pH 3-10 pharmalytes (ProteinSimple, 042-848), 0.5 µl of 4.65, 0.5 µl 9.77 synthetic pI markers (ProteinSimple, 102223 and 102219), and 12.5 µl of 8 M urea solution (Sigma Aldrich®). HPLC grade water was used to make up the final volume to 100 µl. Samples were focused for 1 min at 1.5 kV, followed by 5 min at 3 kV, 280 nm images of the capillary were taken using the Protein Simple software. The resulting electropherograms were analysed using iCE3 software and pI values were assigned (linear relationship between the pI markers).

### Mass spectrometry analysis of proteins

Intact VHH72-Fc protein (10 µg) was first reduced with tris(2-carboxyethyl)phosphine (TCEP; 10 mM) for 30 min at 37°C, after which the reduced protein was separated on an Ultimate 3000 HPLC system (Thermo Fisher Scientific, Bremen, Germany) online connected to an LTQ Orbitrap XL mass spectrometer (Thermo Fischer Scientific). Briefly, approximately 8 µg of protein was injected on a Zorbax 300SB-C18 column (5 µm, 300Å, 1×250mm IDxL; Agilent Technologies) and separated using a 30 min gradient from 5% to 80% solvent B at a flow rate of 100 µl/min (solvent A: 0.1% formic acid and 0.05% trifluoroacetic acid in water; solvent B: 0.1% formic acid and 0.05% trifluoroacetic acid in acetonitrile). The column temperature was maintained at 60°C. Eluting proteins were directly sprayed in the mass spectrometer with an ESI source using the following parameters: spray voltage of 4.2 kV, surface-induced dissociation of 30 V, capillary temperature of 325 °C, capillary voltage of 35 V and a sheath gas flow rate of 7 (arbitrary units). The mass spectrometer was operated in MS1 mode using the orbitrap analyzer at a resolution of 100,000 (at m/z 400) and a mass range of 600-4000 m/z, in profile mode. The resulting MS spectra were deconvoluted with the BioPharma Finder^TM^ 3.0 software (Thermo Fischer Scientific) using the Xtract deconvolution algorithm (isotopically resolved spectra). The deconvoluted spectra were manually annotated.

### Peptide mapping by mass spectrometry

VHH72-Fc protein (15 µg) was diluted with 50 mM triethylammonium bicarbonate (pH 8.5) to a volume of 100 µl. First, protein disulfide bonds were reduced with dithiothreitol (DTT; 5 mM) for 30 min at 55°C and alkylated with iodoacetamide (IAA; 10 mM) for 15 min at room temperature (in the dark). The protein was then digested with LysC endoproteinase (0.25 µg; NEB) for 4 hours at 37°C, followed by sequencing grade trypsin (0.3 µg; Promega) for 16 hours at 37°C. After digestion, trifluoroacetic acid was added to a final concentration of 1%. Prior to LC-MS analysis, the samples were desalted using the Pierce^TM^ C18 Spin Columns (Thermo Fischer Scientific). First, spin columns were activated with 400 µl 50% acetonitrile (2x) and equilibrated with 0.5% trifluoroacetic acid in 5% acetonitrile (2x), after which samples were slowly added on top of the C18 resin. The flow through of each sample was reapplied on the same spin column for 4 times to maximize peptide binding to the resin. After washing the resin with 200 µl of 0.5% trifluoroacetic acid in 5% acetonitrile (2x), peptides were eluted with 2 times 20 µl 70% acetonitrile. Desalted peptide samples were dried and resuspended in 50 µl 0.1% trifluoroacetic acid in 2% acetonitrile.

For the LC-MS/MS analysis, 5 µl of the desalted peptide samples was injected on an in-house manufactured C18 column (ReprosilPur C18 (Dr. Maisch), 5 µm, 0.25×200mm IDxL) and separated using a 30 min gradient from 0% to 70% solvent B at a flow rate of 3 µl/min (solvent A: 0.1% formic acid and 0.05% trifluoroacetic acid in water; solvent B: 0.1% formic acid and 0.05% trifluoroacetic acid in 70% acetonitrile). The column temperature was maintained at 40°C. Eluting proteins were directly sprayed in the LTQ Orbitrap XL mass spectrometer with an ESI source using the following parameters: spray voltage of 4.2 kV, capillary temperature of 275 °C, capillary voltage of 35 V and a sheath gas flow rate of 5 (arbitrary units). The mass spectrometer was operated in data-dependent mode, automatically switching between MS survey scans and MS/MS fragmentation scans of the 3 most abundant ions in each MS scan. Each MS scan (m/z 250-3000) was followed by up to 3 MS/MS scans (isolation window of 3 Da, CID collision energy of 35%, activation time of 30 ms) that fulfill predefined criteria (minimal signal of 5000 counts, exclusion of unassigned and single charged precursors). Precursor ions were excluded from MS/MS selection for 60 sec after two selections within a 30 sec time frame.

The resulting MS/MS spectra were analyzed with the BioPharma Finder^TM^ 3.0 software (Thermo Fischer Scientific) and mapped onto the appropriate protein sequence. For peptide identification, the following parameters were used: maximum peptide mass of 7000 Da, mass accuracy of 5 ppm and a minimum confidence of 0.80. Cysteine carbamidomethylation was set as a fixed modification. Deamidation of asparagine and glutamine, pyroglutamate formation of N-terminal glutamine, glycation of lysine, and oxidation of methionine and tryptophan were set as variable modifications. The search for glycosylation modifications was enabled (CHO-specific). The maximum number of variable modifications per peptide was set at 3.

### SEC-MALS

To determine the molecular mass and aggregation behavior of VHH72-hFc variants, the protein was analyzed by size exclusion chromatography multi-angle laser light scattering (SEC-MALS). For each analysis, 100 µl sample filtered through 0.1 µm Ultrafree-MC centrifugal filters (Merck Millipore) was injected onto a Superdex 200 Increase 10/300 GL Increase SEC column (GE Healthcare) equilibrated with sample buffer, coupled to an online UV detector (Shimadzu), a mini DAWN TREOS (Wyatt) multi-angle laser light scattering detector and an Optilab T-rEX refractometer (Wyatt) at 298 K. The refractive index (RI) increment value (dn/dc value) at 298 K and 658 nm was calculated using SEDFIT v16.1^75^ and used for the determination of the protein concentration and molecular mass. Glycoprotein conjugate analysis was performed using a value of 0.140 ml/g for dn/dc_glycan_ (average carbohydrate dn/dc^76^) and a UV extinction coefficient of zero to account for the negligible absorbance of glycans at 280 nm. Data analysis was carried out using the ASTRA 7.3.2 software.

### Physical and chemical stability testing

Dynamic light scattering was performed using the Uncle instrument (Unchained Labs; Pleasanton, CA, USA). To evaluate the homogeneity and possible aggregate formation or WT-VHH/WT-Fc at 25°C, 10 µL of sample at 1.0, 20 or 30 mg/ml in 25 mM His and 125 mM NaCl, pH6.0 was added to the sample cuvette. Laser and attenuator controls were set at Auto while 10 acquisitions were run per data point with an acquisition time of 10 s for each. Intrinsic tryptophan-fluorescence was monitored upon temperature-induced protein unfolding in the Uncle instrument. Also here, 10 µL of sample at 1 mg/ml was applied to the sample cuvette, and a linear temperature ramp was initiated from 25 to 95°C at a rate of 0.5°C/min, with a pre-run incubation for 180 s. The barycentric mean (BCM) and static light scattering (SLS at 266 nm and 473 nm) signals were plotted against temperature in order to obtain melting temperatures (T_m_) and aggregation onset temperatures (T_agg_), respectively.

Accelerated temperature stress experiments were performed to accelerate the possible formation of soluble and/or insoluble protein aggregates as well as chemical isoforms, *i.e.* asparagine/glutamine deamidation, aspartate/glutamate isomerization, N-terminal pyro-glutamate and truncated species. Stability at elevated temperature was assessed by subjecting duplicate 200 µl 1 mg/ml protein samples (0.02% sodium azide added and 0.22 µm filtered) in 1.5 ml polypropylene Safe-Lock tubes (Eppendorf) to 10 days of storage at 40°C while shaking at 200 rpm in a thermomixer (Eppendorf Thermomixer Comfort 5355).

### Flow cytometric analysis of binding to HEK293S cells expressing the SARS-CoV-1 or SARS-CoV-2 spike protein

To investigate the binding of monomeric VHH72 and VHH72-Fc constructs to spike proteins on the surface of mammalian cells by flow cytometry we used expression plasmids containing the coding sequence of the SARS-CoV-1 and SARS-CoV-2 spikes or of a SARS-CoV-1 spike protein in which the RBD was replaced by that of SARS-CoV-2 as described by Letko *et al*.^14^. The latter was used as a template to generate expression plasmids of the K417N, N439K, E484K, N501Y, and the combination of K417N + E484K + N501Y spike variants by QuickChange site-directed mutagenesis (Agilent) according to the manufacturer’s instructions. Two days after transfecting HEK293T cells or HEK293S cells with spike expression plasmids each combined with a GFP expression plasmid, the cells were collected, washed once with PBS and fixed with 1% PFA for 30 minutes. Binding of human monoclonal antibodies palivizumab, CB6, and S309 and VHH72-Fc or variants thereof was detected with an AF633 conjugated goat anti-human IgG antibody (Invitrogen). Binding of monomeric VHHs to SARS-CoV-1 or SARS-CoV-2 S was detected with a mouse anti-HisTag antibody (AbD Serotec) and an AF647 conjugated donkey anti-mouse IgG antibody (Invitrogen). Following 3 washes with PBS containing 0.5% BSA, the cells were analyzed by flow cytometry using an BD LSRII flow cytometer (BD Biosciences). Binding was calculated as the mean AF633 fluorescence intensity (MFI) of GFP expressing cells (GFP^+^) divided by the MFI of GFP negative cells (GFP^-^). The binding curves were fitted using nonlinear regression (Graphpad 8.0).

### RBD competition assay on Vero E6 cells

SARS-CoV-2 RBD fused to murine IgG Fc (Sino Biological) at a final concentration of 0.4 μg/ml was incubated with a dilution series of monovalent VHH or VHH-Fc proteins and incubated at room temperature for 20 min followed by an additional 10 min incubation on ice. VeroE6 cells grown at sub-confluency were detached by cell dissociation buffer (Sigma) and trypsin treatment. After washing once with PBS, the cells were blocked with 1% BSA in PBS on ice. All remaining steps were also performed on ice. The mixtures containing RBD and VHHs or VHH-Fc fusions were added to the cells and incubated for 1 h. Subsequently, the cells were washed 3 times with PBS containing 0.5% BSA and stained with an AF647 conjugated donkey anti-mouse IgG antibody (Invitrogen) for 1 h. Following three additional washes with PBS containing 0.5% BSA, the cells were analyzed by flow cytometry using a BD LSRII flow cytometer (BD Biosciences).

### Inhibition of ACE-2/RBD interaction by AlphaLISA immunoassay

Dose-dependent inhibition of the interaction of SARS-CoV-2 RBD protein with the ACE-2 receptor was assessed in a competition AlphaLISA (amplified luminescent proximity homogeneous assay). In brief, 2019-nCoV S protein RBD that was biotinylated through an Avi-tag (AcroBiosystems, Cat nr. SPD-C82E9) was loaded on streptavidin coated Alpha Donor beads (Perkin Elmer, Cat nr. 6760002). Human ACE-2-mFc protein (Sino Biological, Cat nr. 10108-H05H) was captured on anti-mouse IgG (Fc specific) acceptor beads (Perkin Elmer, Cat nr. AL105C). Serial dilutions of antibodies and VHH-Fc (final concentration ranging between 100nM - 0,001 nM) were made in assay buffer (PBS containing 0.5% BSA and 0.05% Tween-20), and mixed with biotinylated RBD protein (final concentration 1 nM) in white low binding 384well microtitre plates (F-bottom, Greiner Cat nr 781904). As isotype control antibody, palivizumab, was included. Subsequently, recombinant human ACE-2-Fc (final concentration 0.2 nM) was added to the mixture. After an incubation for 1 hour at room temperature, donor and acceptor beads were added to a final concentration of 20 µg/ml for each in a final volume of 25 µl for an additional incubation of 1 hour at room temperature in the dark. Interaction between beads was assessed after illumination at 680 nm and reading at 615 nm on an Ensight instrument. Graph pad Prism was used for curve fitting and IC_50_ determination of triplicate measurements.

### SARS-CoV pseudovirus neutralization assay

To generate replication-deficient VSV pseudotyped viruses, HEK293T cells, transfected with SARS-CoV-1 S or SARS-CoV-2 S were inoculated with a replication deficient VSV vector containing eGFP and firefly luciferase expression cassettes^77, 78^. After a 1 h incubation at 37°C, the inoculum was removed, cells were washed with PBS and incubated in media supplemented with an anti-VSV G mAb (ATCC) for 16 hours. Pseudotyped particles were then harvested and clarified by centrifugation^17^. For the VSV pseudotype neutralization experiments, the pseudoviruses were incubated for 30 min at 37°C with different dilutions of purified VHH or VHH-Fc fusions or with GFP-binding protein (GBP: a VHH specific for GFP). The incubated pseudoviruses were subsequently added to subconfluent monolayers of VeroE6 cells. Sixteen hours later, the cells were lysed using passive lysis buffer (Promega). The transduction efficiency was quantified by measuring the GFP fluorescence in the prepared cell lysates using a Tecan infinite 200 pro plate reader. As indicated in the legends the GFP fluorescence was normalized using either the GFP fluorescence of non-infected cells and infected cells treated with PBS or the lowest and highest GFP fluorescence value of each dilution series. The IC_50_ was calculated by non-linear regression curve fitting, log(inhibitor) vs. response (four parameters).

### SARS-CoV-2 plaque reduction neutralization test (PRNT)

The PRNT results shown in Figure2f, Supplementary Fig. 6e, and Supplementary Table 1 were performed with SARS-CoV-2 strain BetaCov/Belgium/GHB-03021/2020 (EPI ISL 407976; 2020-02-03) used from passage P6 grown on VeroE6 cells^20^. SARS-CoV-2 viruses belonging to the VoC UK and South African lineages B.1.1.7 (hCoV-19/Belgium/rega-12211513/2020; EPI_ISL_791333, 2020-12-21) and B.1.351 (hCoV-19/Belgium/rega-1920/2021; EPI_ISL_896474, 2021-01-11) were each isolated from nasopharyngeal swabs taken from travelers returning to Belgium in December 2020 and January 2021, respectively, and have recently been described^79^. B.1.1.7 was from a healthy subject and B.1.351 from a patient with respiratory symptoms. Passage 2 B.1.1.7. and B.1.351 virus stocks were grown on Vero E6 cells and median tissue culture infectious doses (TCID_50_) defined by titration using the Spearman-Kärber method^80^. Passage 0 and 2 B.1.1.7 and B.1.351 virus preparations had identical genome sequences.

Dose-dependent neutralization of distinct VHH-Fc constructs was assessed by mixing the VHH-Fc constructs at different concentrations (three-fold serial dilutions starting from a concentration of 20 µg/ml), with 100 PFU SARS-CoV-2 in DMEM supplemented with 2% FBS and incubating the mixture at 37°C for 1h. VHH-Fc-virus complexes were then added to VeroE6 cell monolayers in 12-well plates and incubated at 37°C for 1h. Subsequently, the inoculum mixture was replaced with 0.8% (w/v) methylcellulose in DMEM supplemented with 2% FBS. After 3 days incubation at 37°C, the overlays were removed, the cells were fixed with 3.7% PFA, and stained with 0.5% crystal violet. Half-maximum neutralization titers (PRNT_50_) were defined as the VHH-Fc concentration that resulted in a plaque reduction of 50% across 2 or 3 independent plates.

### Human membrane protein microarray assay

Retrogenix Ltd (Derbyshire, UK) performed the specificity testing of XVR011 using a proprietary cell microarray technology^58^. First, a pre-screen was undertaken to determine the levels of background binding of the test antibody to untransfected HEK293 cells, and to slides spotted with gelatin +/− SARS-CoV-2 spike protein. Slides were spotted with expression vectors encoding both ZsGreen1 and human CD20 or EGFR, and used to reverse-transfect HEK293 cells. After fixation, slides were spotted with 10% gelatin ± SARS-CoV-2 FL Spike-6His (Peak Proteins Ltd, UK; lot # 20200623-01-b1). XVR011 at concentrations of 1, 2.5, or 10 µg/mL, 1 µg/mL Rituximab biosimilar (positive control) or PBS alone, was then added to the slides. Binding to target-expressing cells and untransfected cells was assessed using an AlexaFluor647 (AF647) labelled anti-human IgG Fc detection antibody, followed by fluorescence imaging. Binding to soluble spotted SARS-CoV-2 FL Spike-6His was seen at all concentrations tested and for the full library screen, XVR011 was applied at a concentration of 10 µg/ml. For Library screening, 5,475 expression vectors, encoding both ZsGreen1 and a full-length human plasma membrane protein or a cell surface-tethered human secreted protein, were individually arrayed in duplicate across 16 microarray slides (‘slide-sets’). Vectors encoding a further 371 human heterodimers were co-arrayed across a further microarray slide. An expression vector (pIRES-hEGFR-IRES-ZsGreen1) was spotted in quadruplicate on every slide, and was used to ensure that a minimal threshold of transfection efficiency had been achieved or exceeded on every slide. This minimal threshold had been defined previously. Human HEK293 cells were used for reverse transfection/expression. An additional slide was screened containing SARS-CoV-2 FL Spike-6His spotted in gelatin on top of fixed, untransfected HEK293 cells. Test antibody was added to each slide after cell fixation, giving a final concentration of 10 µg/ml. Detection of binding was performed by using the same fluorescent secondary antibody as used in the Pre-screen. Two replicate slides were screened for each of the slide-sets. Fluorescent images were analyzed and quantitated (for transfection) using ImageQuant software. A protein ‘hit’ is defined as a duplicate spot showing a raised signal compared to background levels. This is achieved by visual inspection using the images gridded on the ImageQuant software. Hits were classified as ‘strong, medium, weak or very weak’, depending on the intensity of the duplicate spots. In a subsequent Confirmation/Specificity screen, all 4 vector library hits, and 2 control receptors (CD20 and EGFR) were over-expressed in HEK293 cells and SARS-CoV-2 FL Spike-6His were spotted onto two slides in gelatin after cell fixation. Confirmatory slides were then probed with XVR011 at 10 µg/ml, Rituximab biosimilar at 1 µg/ml or PBS, followed by Alexafluor647 conjugated anti-human IgG Fc detection antibody.

### Surface plasmon resonance analysis of Fcγ Receptor binding

Surface plasmon resonance (SPR) analyses of XVR011 binding to purified recombinant FcγRI, FcγRIIa (H167), FcγRIIa (R167), FcγRIIIa (V176), FcγRIIIa (F176), FcγRIIIb, and FcγRIIb were performed at Biaffin GmbH & Co KG (Kassel, Germany) on a Biacore T200 instrument (GE Healthcare). His-tagged recombinant human FcγRI, FcγRIIa (H167), FcγRIIa (R167), FcγRIIb, FcγRIIIa (V176), FcγRIIIa (F176), and FcγRIIIb were purchased from Acro Biosystems. Rituximab (Roch Diagnostics), a CD20-specific chimeric mouse-human monoclonal antibody with a human IgG1 Fc, was used as a control for Fcγ receptor binding. Anti-His monoclonal antibody was immobilized on S sensor chip CM3 by covalent coupling and used to capture the His-tagged Fcγ receptors. Ligand capturing concentrations were as follows. FcγRI: 0.04 µg/ml / 0.08 µg/ml, FcγRIIA (H167): 0.05 µg/ml / 0.1 µg/ml, FcγRIIA (R167): 0.1 µg/ml / 0.2 µg/ml, FcγRIIB: 0.05 µg/ml / 0.1 µg/ml, FcγRIIIA (V176) : 0.04 µg/ml / 0.1 µg/ml, FcγRIIIA (F176) : 0.03 µg/ml / 0.12 µg/ml, and FcγRIIIB: 0.06 µg/ml / 0.15 µg/ml. Serially diluted XVR011 and Rituximab, diluted in 10 mM HEPES (pH 7.4), 150 mM NaCl, 3 mM EDTA, 0.05% Tween 20, were then injected on FcγR capture surfaces and analyzed in multi cycle kinetic mode (9-10 concentrations) at 50 µL/min (association time: 1 min; dissociation time: 5 to 10 min). The sensor chips were regenerated with 10 mM of Glycine pH 1.5 (two consecutive 30 s injections) followed by 3 M of guanidine hydrochloride (two consecutive 30 s injections). Data processing (*e.g.* double referencing including blank run subtraction) and evaluation was performed by steady state analysis using Biacore T200 Evaluation Software version 3.1. Statistical evaluation (mean, SD) was conducted using Excel (Microsoft).

### Flow cytometric analysis of antibody binding to Sarbecovirus RBD displayed on the surface of *Saccharomyces cerevisiae*

A pool of plasmids, based on the pETcon yeast surface display expression vector, that encode the RBDs of a set of SARS-CoV2 homologs was generously provided by Dr. Jesse Bloom^38^. This pool was transformed to *E. coli* TOP10 cells by electroporation at the 10 ng scale and plated onto low salt LB agar plates supplemented with carbenicillin. Single clones were selected, grown in liquid low salt LB supplemented with carbenicillin and miniprepped. Selected plasmids were Sanger sequenced with primers covering the entire RBD coding sequence and the process was repeated until every desired RBD homolog had been picked up as a sequence-verified single clone. Additionally, the CDS of the RBD of SARS-CoV2 was ordered as a yeast codon-optimized gBlock and cloned into the pETcon vector by Gibson assembly. The plasmid was transformed into *E. coli*, prepped and sequence-verified as described above. DNA of the selected pETcon RBD plasmids was transformed to *Saccharomyces cerevisiae* strain EBY100 according to the protocol by Gietz and Schiestl^81^ and plated on yeast drop-out medium (SD agar -trp -ura). Single clones were selected and verified by colony PCR for correct insert length. A single clone of each RBD homolog was selected and grown overnight in 10 ml of liquid repressive medium (SRaf -ura -trp) at 28°C. These precultures were then back-diluted to 50 ml of liquid inducing medium (SRaf/Gal -ura -trp) at an OD_600_ of 0.67/ml and grown for 16 hours before harvest. After washing in PBS, the cells were fixed in 1% PFA, washed twice with PBS, blocked with 1% BSA and stained with dilution series of anti-RBD antibodies or palivizumab. Binding of the antibodies was detected using Alexa fluor 633 conjugated anti-human IgG antibodies (Invitrogen). Expression of the surface-displayed myc-tagged RBDs was detected using a FITC conjugated chicken anti-myc antibody (Immunology Consultants Laboratory, Inc.). Following 3 washes with PBS containing 0.5% BSA, the cells were analyzed by flow cytometry using an BD LSRII flow cytometer (BD Biosciences). The binding curves were fitted using nonlinear regression (Graphpad 8.0).

### Mouse challenge experiments

Transgenic (K18-hACE2)2Prlmn mice were originally purchased from The Jackson Laboratory. Locally bred hemizygous 10-14-week-old animals of both sexes were used for experiments. Infection experiments were performed in accordance with the guidelines of the Federation for Laboratory Animal Science Associations and the national animal welfare body. All experiments were in compliance with the German animal protection law and approved by the animal welfare committee of the Regierungspräsidium Freiburg (permit G-20/91).

WT-VHH/12GS-WT-Fc treatments were performed either by injecting approximately 500 µl of a 250 µg/ml solution of WT-VHH/12GS-WT-Fc into the peritoneum of non-anesthetized mice to reach a dose of 5 mg/kg or by applying 40 µl of the WT-VHH/12GS-WT-Fc at 1 mg/ml to the nostrils of isoflurane-anesthetized mice. Infection of isoflurane-anesthetized mice with 8 x 10^3^ PFU of a clinical SARS-CoV-2 isolate (Muc-IMB-1/2020) was done by applying 40 µl samples to the nostrils. Infected mice were monitored for weight loss and clinical signs of disease for 14 days. Mice were sacrificed by cervical dislocation on day 3 post infection to determine infectious virus in lungs by plaque assay using Vero E6 cells. Infected mice were euthanized and scored dead if they lost 25 % of their initial body weight or if showing severe signs of disease. All infection experiments were performed under BSL3 conditions.

### Hamster challenge experiments

The hamster infection model of SARS-CoV-2 performed in the Neyts lab, including the associated analytical procedures, has been described before^82^. In brief, female Syrian hamsters (*Mesocricetus auratus*) of 6-8 weeks’ old were anesthetized with ketamine/xylazine/atropine and inoculated intranasally with 50 µL containing 2×10^6^ TCID50 SARS-CoV-2 BetaCov/Belgium/GHB-03021/2020. Animals were treated once by intraperitoneal injection, either 1 day prior or 16-24h post SARS-CoV-2 challenge. Hamsters were monitored daily for appearance, behavior and weight. At day 4 pi, hamsters were euthanized by i.p. injection of 500 μL Dolethal (200mg/ml sodium pentobarbital, Vétoquinol SA). Lung, ileum and stool were collected and viral RNA and infectious virus were quantified by RT-qPCR and end-point virus titration, respectively. Serum samples were collected at day 4 pi for PK analysis. Micro-CT data of *in vivo* hamster lungs were acquired using dedicated small animal micro-CT scanners, either using the X-cube (Molecubes, Ghent, Belgium) or the Skyscan 1278 (Bruker Belgium, Kontich, Belgium), as described earlier ^82^. Visualization and quantification of reconstructed micro-CT data were performed with DataViewer and CTan software (Bruker Belgium). Housing conditions and experimental procedures were approved by the ethics committee of animal experimentation of KU Leuven (license P065-2020).

The hamster model performed at Viroclinics was as follows. Male 14-15 weeks old Syrian hamsters (*Mesocricetus auratus*), weighing 106 to 158 grams were obtained from Janvier. The animals were housed for 7 days in individually ventilated cages (2 hamsters per cage) under BSL-2 conditions prior to transfer to a BSL-3 animal house. Six animals were used per group. Palivizumab (20 mg/kg), humVHH_S56A/LALAPG-Fc/Gen2 (20, 7, and 2 mg/kg) and (humVHH_S56A)_2_/LALAPG-Fc/Gen2 (20, 7 and 2 mg/kg) were administered by intraperitoneal injection in a volume of 1ml per 100g of hamster body mass 4 hours after the challenge. One group of hamsters received 20 mg/kg of humVHH_S56A/LALAPG-Fc/Gen2 by intraperitoneal injection 24 hours before challenge infection. All animals were infected intranasally with 10^4^ TCID_50_ of SARS-CoV-2 BetaCoV/Munich/BavPat1/2020 (Passage 3, grown on VeroE6 cells) in a total volume of 0.1 ml. On day 4 after infection all animals were euthanized by exsanguination under anesthesia.

Ethics approval for the hamster study performed at Viroclinics was registered under number: 277002015283-WP11. On day 2 after infection, throat swabs were collected in 1.5 ml virus transport medium, aliquoted and stored. Upon necropsy, broncho alveolar lavage was performed on the left lung with 1ml of PBS and tissue samples were collected and stored in 10% formalin for histopathology and immunohistochemistry (left lung and left nasal turbinate) and frozen for virological analysis (right lung and right nasal turbinate). For virological analysis, tissue samples were weighed, homogenized in infection medium and centrifuged briefly before titration. Serum samples on day 4 (approximately 500 μl per animal) post infection were collected during euthanization and immediately transferred to appropriate tubes containing a clot activator. Throat swabs, BAL and tissue homogenates were used to detect viral RNA. To this end, RNA was isolated (nucleic acid purification on the MagNA Pure 96; Roche Life Science), reverse transcribed and Taqman PCR (on the 7500 RealTime PCR system; Applied Biosystems) was performed using specific primers (E_Sarbeco_F: 5’ACAGGTACGTTAATAGTTAATAGCGT3’ and E_Sarbeco_R: 5’ATATTGCAGCAGTACGCACACA3’) and probe (E_Sarbeco_P1: 5’ACACTAGCCATCCTTACTGCGCTTCG3’) specific for betacoronavirus E gene. Quadruplicate 10-fold serial dilutions were used to determine the virus titers in confluent layers of Vero E6 cells. To this end, serial dilutions of the samples (throat swabs, BAL and tissue homogenates) were made and incubated on Vero E6 monolayers for 1 hour at 37 degrees. Vero E6 monolayers were then washed and incubated for 4-6 days at 37 degrees after which plates were stained and scored using the vitality marker WST8 (colorimetric readout). To this end, WST-8 stock solution was prepared and added to the plates. Per well, 20 μL of this solution (containing 4 μL of the ready-to-use WST-8 solution from the kit and 16 μL infection medium, 1:5 dilution) was added and incubated 3-5 hours at RT. Subsequently, plates were measured for optical density at 450 nm (OD450) using a microplate reader and visual results of the positive controls (cytopathic effect (cpe)) were used to set the limits of the WST-8 staining (OD value associated with cpe). Viral titers (TCID_50_/ml or/g) were calculated using the method of Spearman-Karber^80^.

### Quantification of RBD-binding VHH-Fcs in serum samples of challenged hamsters

Concentrations of XVR011 and preleads in hamster serum samples were quantified in a competition AlphaLISA (described above). This assay detects the inhibition of the interaction of SARS-CoV-2 RBD protein with monovalent humVHH72_S56A. This homogeneous assay without wash steps in a closed system is considered advantageous for testing samples from virus challenged animals. In standards, spiked controls and diluted samples were mixed with 3 nM VHH72-h1 (S56A)-Flag3-His3 and 2.5 nM biotinylated SARS-CoV-2 RBD protein in white low binding 384-well microtitre plates (F-bottom, Greiner Cat nr 781904). After an incubation for 1 hour at room temperature, donor and acceptor beads were added to a final concentration of 20 µg/ml for an additional incubation of 1 hour at room temperature in the dark. Interaction between beads was assessed after illumination at 680 nm and reading at 615 nm on an Ensight instrument. Standard curves were generated by 1.7-fold serial dilutions of the respective compound in pooled hamster serum diluted in alphascreen assay buffer (PBS containing 0.5% BSA and 0.05% Tween-20). Concentrations were back-calculated by 4PL interpolation using Graphpad Prism (9.0).

## Data availability

The main data supporting the findings of this study are available in the paper and its Supplementary information. The associated raw data are too numerous to be readily shared publicly and can be made available from the corresponding authors upon reasonable request.

## References

1. Zhu, N. et al. A Novel Coronavirus from Patients with Pneumonia in China, 2019. N. Engl. J. Med. 382, 727–733 (2020).

2. Shrock, E. et al. Viral epitope profiling of COVID-19 patients reveals cross-reactivity and correlates of severity. Science 370, (2020).

3. Polack, F. P. et al. Safety and Efficacy of the BNT162b2 mRNA Covid-19 Vaccine. N. Engl. J. Med. (2020) doi:10.1056/NEJMoa2034577.

4. Voysey, M. et al. Safety and efficacy of the ChAdOx1 nCoV-19 vaccine (AZD1222) against SARS-CoV-2: an interim analysis of four randomised controlled trials in Brazil, South Africa, and the UK. Lancet Lond. Engl. 397, 99–111 (2021).

5. Tenforde, M. W. et al. Influenza vaccine effectiveness in inpatient and outpatient settings in the United States, 2015 - 2018. Clin. Infect. Dis. Off. Publ. Infect. Dis. Soc. Am. (2020) doi:10.1093/cid/ciaa407.

6. Beaudoin-Bussières, G. et al. Decline of Humoral Responses against SARS-CoV-2 Spike in Convalescent Individuals. mBio 11, (2020).

7. Greaney, A. J. et al. Comprehensive mapping of mutations to the SARS-CoV-2 receptor-binding domain that affect recognition by polyclonal human serum antibodies. bioRxiv 2020.12.31.425021 (2021) doi:10.1101/2020.12.31.425021.

8. Lucas, C. et al. Kinetics of antibody responses dictate COVID-19 outcome. medRxiv 2020.12.18.20248331 (2020) doi:10.1101/2020.12.18.20248331.

9. Weinreich, D. M. et al. REGN-COV2, a Neutralizing Antibody Cocktail, in Outpatients with Covid-19. N. Engl. J. Med. (2020) doi:10.1056/NEJMoa2035002.

10. Chen, P. et al. SARS-CoV-2 Neutralizing Antibody LY-CoV555 in Outpatients with Covid-19. N. Engl. J. Med. (2020) doi:10.1056/NEJMoa2029849.

11. Libster, R. et al. Early High-Titer Plasma Therapy to Prevent Severe Covid-19 in Older Adults. N. Engl. J. Med. (2021) doi:10.1056/NEJMoa2033700.

12. Pyzik, M. et al. The Neonatal Fc Receptor (FcRn): A Misnomer? Front. Immunol. 10, 1540 (2019).

13. Baum, A. et al. Antibody cocktail to SARS-CoV-2 spike protein prevents rapid mutational escape seen with individual antibodies. Science (2020) doi:10.1126/science.abd0831.

14. Eurosurveillance editorial team. Updated rapid risk assessment from ECDC on the risk related to the spread of new SARS-CoV-2 variants of concern in the EU/EEA - first update. Euro Surveill. Bull. Eur. Sur Mal. Transm. Eur. Commun. Dis. Bull. 26, (2021).

15. Wang, P. et al. Increased Resistance of SARS-CoV-2 Variants B.1.351 and B.1.1.7 to Antibody Neutralization. bioRxiv 2021.01.25.428137 (2021) doi:10.1101/2021.01.25.428137.

16. Diamond, M. et al. SARS-CoV-2 variants show resistance to neutralization by many monoclonal and serum-derived polyclonal antibodies. Res. Sq. (2021) doi:10.21203/rs.3.rs-228079/v1.

17. Wrapp, D. et al. Structural Basis for Potent Neutralization of Betacoronaviruses by Single-Domain Camelid Antibodies. Cell 181, 1004–1015.e15 (2020).

18. Letko, M., Marzi, A. & Munster, V. Functional assessment of cell entry and receptor usage for SARS-CoV-2 and other lineage B betacoronaviruses. Nat. Microbiol. 5, 562–569 (2020).

19. International Committee on Taxonomy of Viruses Executive Committee. The new scope of virus taxonomy: partitioning the virosphere into 15 hierarchical ranks. Nat. Microbiol. 5, 668–674 (2020).

20. Boudewijns, R. et al. STAT2 signaling restricts viral dissemination but drives severe pneumonia in SARS-CoV-2 infected hamsters. Nat. Commun. 11, 5838 (2020).

21. Cai, Y. et al. Distinct conformational states of SARS-CoV-2 spike protein. Science 369, 1586–1592 (2020).

22. Piccoli, L. et al. Mapping Neutralizing and Immunodominant Sites on the SARS-CoV-2 Spike Receptor-Binding Domain by Structure-Guided High-Resolution Serology. Cell 183, 1024–1042.e21 (2020).

23. Chi, X. et al. Humanized single domain antibodies neutralize SARS-CoV-2 by targeting the spike receptor binding domain. Nat. Commun. 11, 4528 (2020).

24. Custódio, T. F. et al. Selection, biophysical and structural analysis of synthetic nanobodies that effectively neutralize SARS-CoV-2. Nat. Commun. 11, 5588 (2020).

25. Koenig, P.-A. et al. Structure-guided multivalent nanobodies block SARS-CoV-2 infection and suppress mutational escape. Science (2021) doi:10.1126/science.abe6230.

26. Hanke, L. et al. An alpaca nanobody neutralizes SARS-CoV-2 by blocking receptor interaction. Nat. Commun. 11, 4420 (2020).

27. Esparza, T. J., Martin, N. P., Anderson, G. P., Goldman, E. R. & Brody, D. L. High affinity nanobodies block SARS-CoV-2 spike receptor binding domain interaction with human angiotensin converting enzyme. Sci. Rep. 10, 22370 (2020).

28. Schoof, M. et al. An ultrapotent synthetic nanobody neutralizes SARS-CoV-2 by stabilizing inactive Spike. Science 370, 1473–1479 (2020).

29. Huo, J. et al. Neutralizing nanobodies bind SARS-CoV-2 spike RBD and block interaction with ACE2. Nat. Struct. Mol. Biol. 27, 846–854 (2020).

30. Li, W. et al. High Potency of a Bivalent Human VH Domain in SARS-CoV-2 Animal Models. Cell 183, 429–441.e16 (2020).

31. Xiang, Y. et al. Versatile and multivalent nanobodies efficiently neutralize SARS-CoV-2. Science 370, 1479–1484 (2020).

32. Moreau, G. B. et al. Evaluation of K18-hACE2 Mice as a Model of SARS-CoV-2 Infection. Am. J. Trop. Med. Hyg. 103, 1215–1219 (2020).

33. McCray, P. B. et al. Lethal infection of K18-hACE2 mice infected with severe acute respiratory syndrome coronavirus. J. Virol. 81, 813–821 (2007).

34. Winkler, E. S. et al. SARS-CoV-2 infection of human ACE2-transgenic mice causes severe lung inflammation and impaired function. Nat. Immunol. 21, 1327–1335 (2020).

35. Camacho, C. J. & Zhang, C. FastContact: rapid estimate of contact and binding free energies. Bioinforma. Oxf. Engl. 21, 2534–2536 (2005).

36. Zhou, D. et al. Structural basis for the neutralization of SARS-CoV-2 by an antibody from a convalescent patient. Nat. Struct. Mol. Biol. (2020) doi:10.1038/s41594-020-0480-y.

37. Liu, H. et al. Cross-Neutralization of a SARS-CoV-2 Antibody to a Functionally Conserved Site Is Mediated by Avidity. Immunity 53, 1272–1280.e5 (2020).

38. Starr, T. N. et al. Deep Mutational Scanning of SARS-CoV-2 Receptor Binding Domain Reveals Constraints on Folding and ACE2 Binding. Cell 182, 1295–1310.e20 (2020).

39. Yang, J. & Zhang, Y. I-TASSER server: new development for protein structure and function predictions. Nucleic Acids Res. 43, W174–181 (2015).

40. Guex, N. & Peitsch, M. C. SWISS-MODEL and the Swiss-PdbViewer: an environment for comparative protein modeling. Electrophoresis 18, 2714–2723 (1997).

41. Abraham, M. J. et al. GROMACS: High performance molecular simulations through multi-level parallelism from laptops to supercomputers. SoftwareX 1–2, 19–25 (2015).

42. Wrapp, D. et al. Cryo-EM structure of the 2019-nCoV spike in the prefusion conformation. Science 367, 1260–1263 (2020).

43. Yan, R. et al. Structural basis for the recognition of SARS-CoV-2 by full-length human ACE2. Science 367, 1444–1448 (2020).

44. Walls, A. C. et al. Structure, Function, and Antigenicity of the SARS-CoV-2 Spike Glycoprotein. Cell 181, 281–292.e6 (2020).

45. Zohar, T. & Alter, G. Dissecting antibody-mediated protection against SARS-CoV-2. Nat. Rev. Immunol. 20, 392–394 (2020).

46. Arvin, A. M. et al. A perspective on potential antibody-dependent enhancement of SARS-CoV-2. Nature 584, 353–363 (2020).

47. Schlothauer, T. et al. Novel human IgG1 and IgG4 Fc-engineered antibodies with completely abolished immune effector functions. Protein Eng. Des. Sel. PEDS 29, 457–466 (2016).

48. DeFrancesco, L. COVID-19 antibodies on trial. Nat. Biotechnol. 38, 1242–1252 (2020).

49. Yang, L. et al. COVID-19 antibody therapeutics tracker: a global online database of antibody therapeutics for the prevention and treatment of COVID-19. Antib. Ther. 3, 205–212 (2020).

50. Shi, R. et al. A human neutralizing antibody targets the receptor-binding site of SARS-CoV-2. Nature 584, 120–124 (2020).

51. Xu, D. et al. In vitro characterization of five humanized OKT3 effector function variant antibodies. Cell. Immunol. 200, 16–26 (2000).

52. Wines, B. D., Powell, M. S., Parren, P. W., Barnes, N. & Hogarth, P. M. The IgG Fc contains distinct Fc receptor (FcR) binding sites: the leukocyte receptors Fc gamma RI and Fc gamma RIIa bind to a region in the Fc distinct from that recognized by neonatal FcR and protein A. J. Immunol. Baltim. Md 1950 164, 5313–5318 (2000).

53. Chan, J. F.-W. et al. Simulation of the clinical and pathological manifestations of Coronavirus Disease 2019 (COVID-19) in golden Syrian hamster model: implications for disease pathogenesis and transmissibility. Clin. Infect. Dis. Off. Publ. Infect. Dis. Soc. Am. (2020) doi:10.1093/cid/ciaa325.

54. Spiteri, G. et al. First cases of coronavirus disease 2019 (COVID-19) in the WHO European Region, 24 January to 21 February 2020. Euro Surveill. Bull. Eur. Sur Mal. Transm. Eur. Commun. Dis. Bull. 25, (2020).

55. Jain, T. et al. Biophysical properties of the clinical-stage antibody landscape. Proc. Natl. Acad. Sci. U. S. A. 114, 944–949 (2017).

56. Jarasch, A. et al. Developability assessment during the selection of novel therapeutic antibodies. J. Pharm. Sci. 104, 1885–1898 (2015).

57. Korber, B. et al. Tracking Changes in SARS-CoV-2 Spike: Evidence that D614G Increases Infectivity of the COVID-19 Virus. Cell (2020) doi:10.1016/j.cell.2020.06.043.

58. Freeth, J. & Soden, J. New Advances in Cell Microarray Technology to Expand Applications in Target Deconvolution and Off-Target Screening. SLAS Discov. Adv. Life Sci. R D 25, 223–230 (2020).

59. Pinto, D. et al. Cross-neutralization of SARS-CoV-2 by a human monoclonal SARS-CoV antibody. Nature 583, 290–295 (2020).

60. Leung, K., Shum, M. H., Leung, G. M., Lam, T. T. & Wu, J. T. Early transmissibility assessment of the N501Y mutant strains of SARS-CoV-2 in the United Kingdom, October to November 2020. Euro Surveill. Bull. Eur. Sur Mal. Transm. Eur. Commun. Dis. Bull. 26, (2021).

61. Zahradník, J. et al. SARS-CoV-2 RBD in vitro evolution follows contagious mutation spread, yet generates an able infection inhibitor. bioRxiv 2021.01.06.425392 (2021) doi:10.1101/2021.01.06.425392.

62. Thomson, E. C. et al. Circulating SARS-CoV-2 spike N439K variants maintain fitness while evading antibody-mediated immunity. Cell (2021) doi:10.1016/j.cell.2021.01.037.

63. Tegally, H. et al. Emergence and rapid spread of a new severe acute respiratory syndrome-related coronavirus 2 (SARS-CoV-2) lineage with multiple spike mutations in South Africa. medRxiv 2020.12.21.20248640 (2020) doi:10.1101/2020.12.21.20248640.

64. SARS-CoV-2 reinfection by the new Variant of Concern (VOC) P.1 in Amazonas, Brazil - SARS-CoV-2 coronavirus / nCoV-2019 Genomic Epidemiology. Virological https://virological.org/t/sars-cov-2-reinfection-by-the-new-variant-of-concern-voc-p-1-in-amazonas-brazil/596 (2021).

65. Lv, Z. et al. Structural basis for neutralization of SARS-CoV-2 and SARS-CoV by a potent therapeutic antibody. Science 369, 1505–1509 (2020).

66. Zhang, C. et al. Development and structural basis of a two-MAb cocktail for treating SARS-CoV-2 infections. Nat. Commun. 12, 264 (2021).

67. Yuan, M. et al. Structural basis of a shared antibody response to SARS-CoV-2. Science 369, 1119–1123 (2020).

68. Winkler, E. S. et al. Human neutralizing antibodies against SARS-CoV-2 require intact Fc effector functions and monocytes for optimal therapeutic protection. BioRxiv Prepr. Serv. Biol. (2020) doi:10.1101/2020.12.28.424554.

69. Andreano, E. et al. Extremely potent human monoclonal antibodies from COVID-19 convalescent patients. Cell (2021) doi:10.1016/j.cell.2021.02.035.

70. Lindorff-Larsen, K. et al. Improved side-chain torsion potentials for the Amber ff99SB protein force field. Proteins 78, 1950–1958 (2010).

71. Weninger, A., Hatzl, A.-M., Schmid, C., Vogl, T. & Glieder, A. Combinatorial optimization of CRISPR/Cas9 expression enables precision genome engineering in the methylotrophic yeast Pichia pastoris. J. Biotechnol. 235, 139–149 (2016).

72. Krainer, F. W. et al. Knockout of an endogenous mannosyltransferase increases the homogeneity of glycoproteins produced in Pichia pastoris. Sci. Rep. 3, 3279 (2013).

73. Lee, M. E., DeLoache, W. C., Cervantes, B. & Dueber, J. E. A Highly Characterized Yeast Toolkit for Modular, Multipart Assembly. ACS Synth. Biol. 4, 975–986 (2015).

74. Wu, S. & Letchworth, G. J. High efficiency transformation by electroporation of Pichia pastoris pretreated with lithium acetate and dithiothreitol. BioTechniques 36, 152–154 (2004).

75. Zhao, H., Brown, P. H. & Schuck, P. On the distribution of protein refractive index increments. Biophys. J. 100, 2309–2317 (2011).

76. Tsunashima, Y., Moro, K., Chu, B. & Liu, T. Y. Characterization of group C meningococcal polysaccharide by light-scattering spectroscopy. III. Determination of molecular weight, radius of gyration, and translational diffusional coefficient. Biopolymers 17, 251–265 (1978).

77. Berger Rentsch, M. & Zimmer, G. A vesicular stomatitis virus replicon-based bioassay for the rapid and sensitive determination of multi-species type I interferon. PloS One 6, e25858 (2011).

78. Hoffmann, M. et al. SARS-CoV-2 Cell Entry Depends on ACE2 and TMPRSS2 and Is Blocked by a Clinically Proven Protease Inhibitor. Cell 181, 271–280.e8 (2020).

79. Abdelnabi, R. et al. Comparative infectivity and pathogenesis of emerging SARS-CoV-2 variants in Syrian hamsters. bioRxiv 2021.02.26.433062 (2021) doi:10.1101/2021.02.26.433062.

80. Kärber, G. Beitrag zur kollektiven Behandlung pharmakologischer Reihenversuche. Naunyn-Schmiedebergs Arch. Für Exp. Pathol. Pharmakol. 162, 480–483 (1931).

81. Gietz, R. D. & Schiestl, R. H. High-efficiency yeast transformation using the LiAc/SS carrier DNA/PEG method. Nat. Protoc. 2, 31–34 (2007).

82. Kaptein, S. J. F. et al. Favipiravir at high doses has potent antiviral activity in SARS-CoV-2-infected hamsters, whereas hydroxychloroquine lacks activity. Proc. Natl. Acad. Sci. U. S. A. 117, 26955–26965 (2020).

