## Supplemental Figures and Tables for "Drug development of an affinity enhanced, broadly neutralizing heavy chain-only antibody that restricts SARS-CoV-2 in rodents" (from another version of this preprint: version 1, 2021-03-08)

Supplementary Figure 1

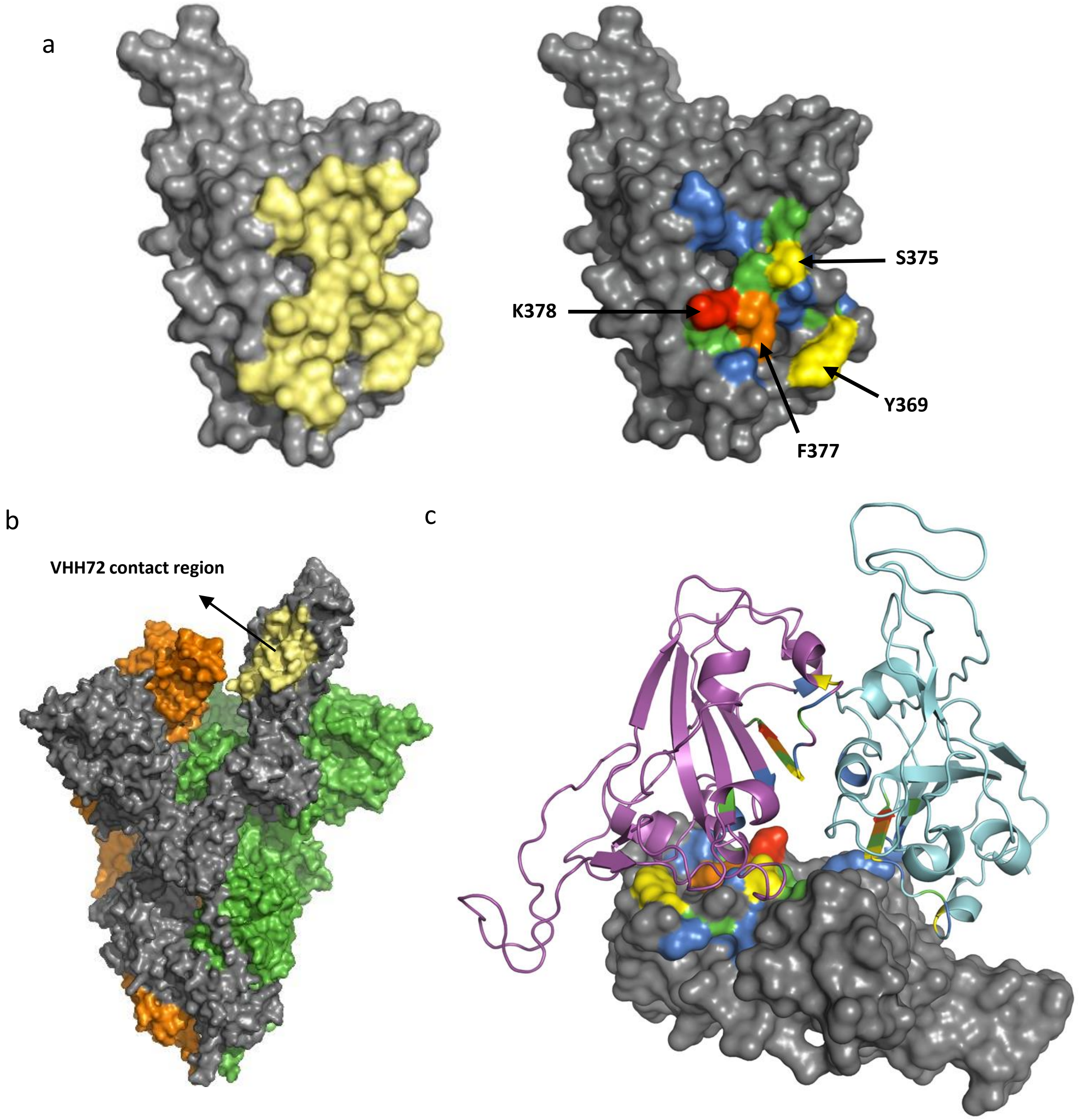


**Supplementary Figure 1. The VHH72 contact region in SARS-CoV-2 RBD. a.** Left: RBD as surface-view with the VHH72 contact region indicated in yellow, for which PDBePISA predicts residues 368-379, 381-385, 404, 405, 407, 408, 435-437, 503, 504 and 508. Right: relative contribution of the contact residues based on electrostatic plus desolvation free energy (kcal/mol) per residue calculated by FastContact. F377 and K378 represent a prominent hot-spot for VHH72 binding, as does Y369, the preferential orientation of which appears different between SARS-CoV-1 and SARS-CoV-2, which was the focus of our affinity maturation campaign. Red: -9.8 (K378); orange: -4.3 (F377); yellow: -2.21 (S375) and -2.01 (Y369); green: -2.0 to -1.0 (A372, T376, C379 and Y508); blue: -1.0 to -0.5 (S371, F374, P384, V407, R408, T436). **b**. Location of the contact region of VHH72 (light yellow) on a SARS-CoV-2 pre-fusion spike protein with one RBD in 'up' position (protomer in grey). Shown with a surface view of the spike protein from pdb-entry 7DD8, originally complexed with a Fab of mouse monoclonal antibody 3C1. **c**. The contact region of VHH72 is occluded in the RBD-closed state of SARS-CoV-2 spike pre-fusion protein. Apex-view of intact wild-type SARS-CoV-2 pre-fusion closed state spike trimer pdb-entry 6XR8, showing only the three RBDs. Chain A, grey-surface; chain B, cyan-cartoon; chain C, magenta-cartoon. The residues at the VHH72 binding contact region are shown with the same FastContact contribution colors as above.

Supplementary Figure 2

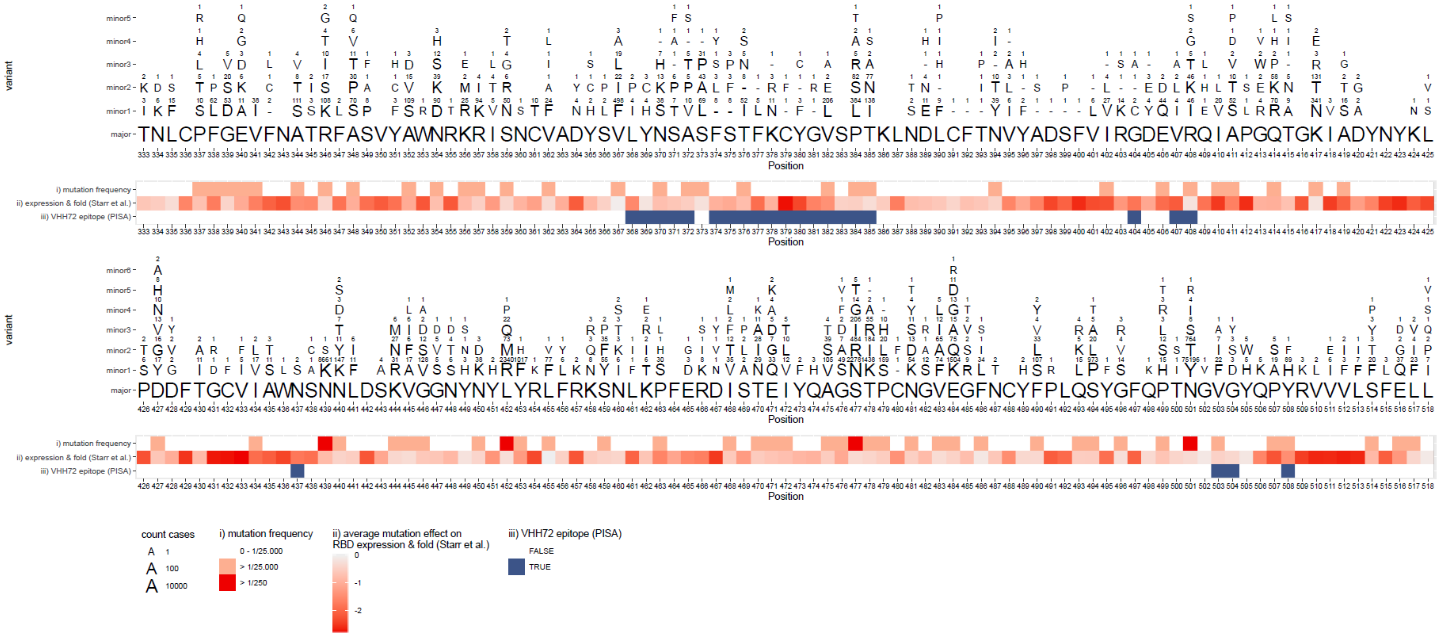


**Supplementary Figure 2. The contact region of VHH72 is highly conserved in circulating SARS-CoV-2 viruses.** The amino acid sequence of SARS-CoV-2 RBD (spike protein amino acid positions 333-516 of Wuhan-Hu-1 isolate) is shown with all missense mutations, detected at least once in 440,769 SARS-CoV-2 genomes analyzed, depicted above each residue. Variants are ordered vertically at each position, according to frequency represented by data point size and the number of observed cases. i) Mutation frequency (white: low mutation frequency < 1/25.000, pink: intermediate mutation frequency, red: high mutation frequency > 1/250), ii) average mutation effect on the RBD yeast surface display expression as a proxy for fold stability, according to Starr *et al.*[38] (white: no effect; red: strong negative effect), iii) the VHH72 contact region according to PISA calculations.

Supplementary Figure 3

a

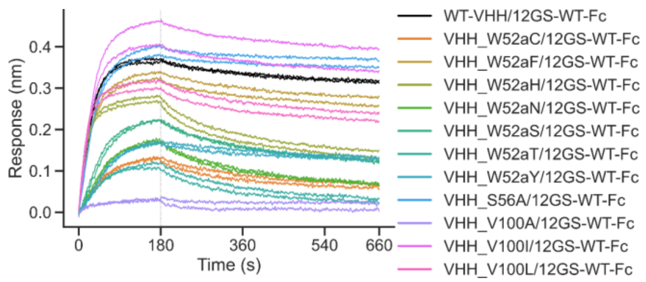


b

| Variant | Affinity mutation | | | n | % signal decrease during 480s dissociation |
|---|---|---|---|---|---|
| | position | original | mutation | | averaged after double reference subtraction |
| WT-VHH/12GS-WT-Fc | - | - | - | 24 | 13.0 |
| VHH_W52aC/12GS-WT-Fc | 52a | W | C | 2 | 52.8 |
| VHH_W52aF/12GS-WT-Fc | 52a | W | F | 2 | 19.1 |
| VHH_W52aH/12GS-WT-Fc | 52a | W | H | 2 | 49.1 |
| VHH_W52aN/12GS-WT-Fc | 52a | W | N | 2 | 60.5 |
| VHH_W52aS/12GS-WT-Fc | 52a | W | S | 2 | 45.0 |
| VHH_W52aT/12GS-WT-Fc | 52a | W | T | 2 | 75.5 |
| VHH_W52aY/12GS-WT-Fc | 52a | W | Y | 2 | 23.2 |
| VHH_S56A/12GS-WT-Fc | 56 | S | A | 2 | 7.8 |
| VHH_V100A/12GS-WT-Fc | 100 | V | A | 2 | 51.9 |
| VHH_V100I/12GS-WT-Fc | 100 | V | I | 2 | 15.3 |
| VHH_V100L/12GS-WT-Fc | 100 | V | L | 2 | 25.7 |

c

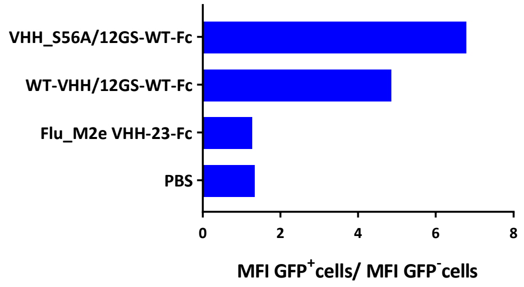


**Supplementary Figure 3. Selection of VHH72_S56A-Fc. a.** Association (180 s) and dissociation (480 s) of duplicate WT-VHH/12GS-WT-Fc and VHH-mutants in diluted crude *Pichia pastoris* supernatant on SARS-CoV-2 RBD-mFc coated biosensors. **b.** Signal decrease during the dissociation phase of the indicated VHH/12GS-WT-Fc variants expressed in crude *Pichia pastoris* supernatant. **c.** The S56A substitution increases the affinity of VHH/12GS-WT-Fc for the SARS-CoV-2 spike protein expressed on the surface of mammalian cells. HEK293T cells were cotransfected with a GFP and a SARS-CoV-2 spike expression vector and then stained with 2-fold diluted crude *Pichia pastoris* supernatant expressing the indicated VHH-Fc proteins and analyzed by flow cytometry. Flu_M2e VHH-23-Fc is directed against the influenza A virus matrix 2 protein and was included as a negative control. The bars in the graph represent the ratio of the mean fluorescence intensity (MFI) of the GFP-expressing cells over the MFI of the GFP negative cells.

Supplementary Figure 4

```
VHH72:        QVQLQESGGGLVQAGGSLRLSCAASGRTFSEYAMGWFRQAPGKEREFVATISWSGGSTYYTDSVKGRFTISRDNAKNTVYLQMNSLKPDDTAVYYCAAAGLGTVVSEWDYDYDYWGQGTQVTVSS
humVHH:       EVQLVESGGGLVQPGGSLRLSCAASGRTFSEYAMGWFRQAPGKEREFVATISWSGGSTYYTDSVKGRFTISRDNAKNTVYLQMNSLRPEDTAVYYCAAAGLGTVVSEWDYDYDYWGQGTLVTVSS
humVHH_S56A:  EVQLVESGGGLVQPGGSLRLSCAASGRTFSEYAMGWFRQAPGKEREFVATISWSGGATYYTDSVKGRFTISRDNAKNTVYLQMNSLRPEDTAVYYCAAAGLGTVVSEWDYDYDYWGQGTLVTVSS

IGHV3-JH cons: EVQLVESGGGLVQPGGSLRLSCAASGFTFSSYAMHWVRQAPGKGLEWVSVISSDGSSTYYADSVKGRFTISRDNSKNTLYLQMNSLRAEDTAVYYCAR...............WGQGTLVTVSS
```

**Supplementary Figure 4. Humanization of VHH72.** Amino acid residue sequence of VHH72, humVHH, humVHH_S56A, and the human IGHV3-JH consensus sequence. Boxed amino acids indicate CDR1, -2, and -3 in VHH72, humVHH, and humVHH_S56A. Amino acid residues in red represent the mutations in the framework regions that were introduced to generate the humVHH variant. Substitution S56A is marked in purple.

Supplementary Figure 5

a

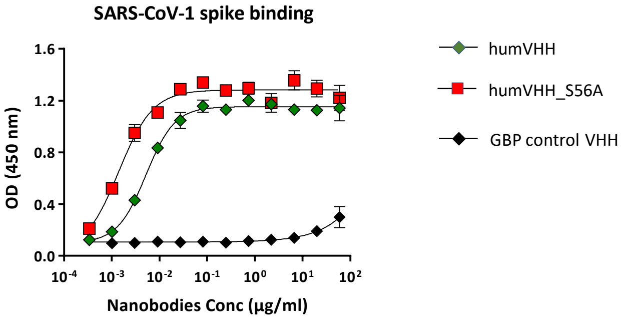


b

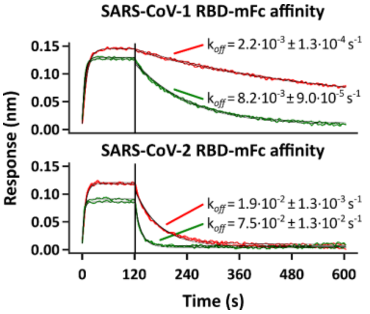


**Supplementary Figure 5. HumVHH_S56A has increased affinity for SARS-CoV-1 spike and RBD. a.** Binding to SARS-CoV-1 spike determined by ELISA (data points are mean ± SD; n=3). **b.** BLI sensorgram of humVHH and humVHH_S56A (each 200 nM) binding to immobilized RBD from SARS-CoV-1 and SARS-CoV-2 fused to mouse Fc. Red (humVHH_S56A) and green (humVHH) curves represent double reference-subtracted data and a partial local fit of the data to a 1:1 binding curve is colored black. Dissociation constants indicated ($k_{off}$) are mean and standard deviation of duplicate measures. GBP: GFP-binding protein = a VHH directed against GFP.

Supplementary Figure 6

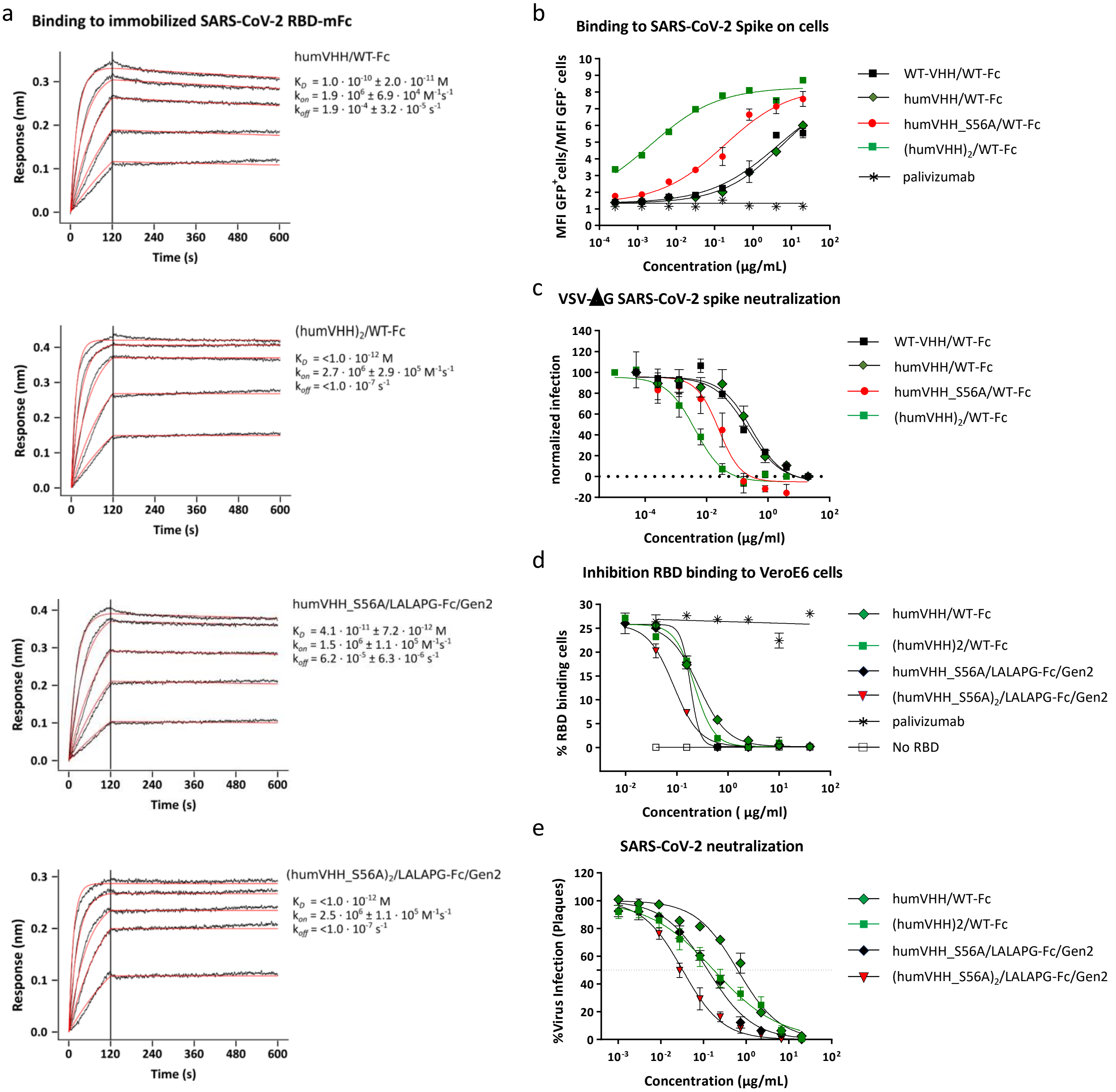

**Supplementary Figure 6. Tetravalent VHH72-Fc has increased affinity for SARS-CoV-2 RBD**. **a.** Binding affinity of bivalent (humVHH/WT-Fc and humVHH_S56A/LALAPG-Fc/Gen2) versus tetravalent ((humVHH)$_2$/WT-Fc and (humVHH_S56A)$_2$/LALAPG-Fc/Gen2) VHH72-Fc variants (twofold dilution series starting at 30 nM) to immobilized mouse Fc fused SARS-CoV-2 RBD (RBD-mFc). Black lines represent double reference-subtracted data and the fit of the data to a 1:1 binding curve is colored red. Replicate BLI analyses of which a single representative replicate is shown. Kinetics parameters are average of duplicate experiments. **b.** Binding of the indicated VHH72-Fc constructs to mammalian cell surface expressed SARS-CoV-2 spike determined by flow cytometry. The graph shows the mean (n=2) ratio of the MFI of transfected (GFP+) cells over the MFI of non-transfected (GFP-) cells. Data are the same as in Figure 2c except that tetravalent (humVHH)$_2$/WT-Fc is included here. **c.** VSV SARS-CoV-2 spike pseudotype virus neutralization assay with the indicated VHH72-Fc constructs. The graph shows the mean (n=3 ± SEM) GFP fluorescence normalized to the lowest and highest value of each dilution series. Data for WT-VHH/WT-Fc, humVHH/WT-Fc, and humVHH_S56A/WT-Fc are the same as shown in Figure 2g. **d.** Dose-dependent inhibition of SARS-CoV-2 RBD binding to the surface of VeroE6 cells in the presence of the indicated VHH72-Fc constructs as determined by flow cytometry. The graph shows the mean (n=2 ± SD) percentage of cells that bind RBD. Data for humVHH/WT-Fc, PBS and no RBD are the same as in Figure 2e. **e.** SARS-CoV-2 plaque reduction neutralization assay with 3 fold serial dilutions of the indicated VHH-Fc fusion constructs. Thirty-six hours after infection, the cells fixed with 3.7% paraformaldehyde and stained with 0.5% crystal violet. Data points in the graph represent the number of plaques and are representative of one experiment that was repeated once. Data for humVHH/WT-Fc and (humVHH)$_2$/WT-Fc are the same as in Figure 2f.

### Supplementary Figure 7

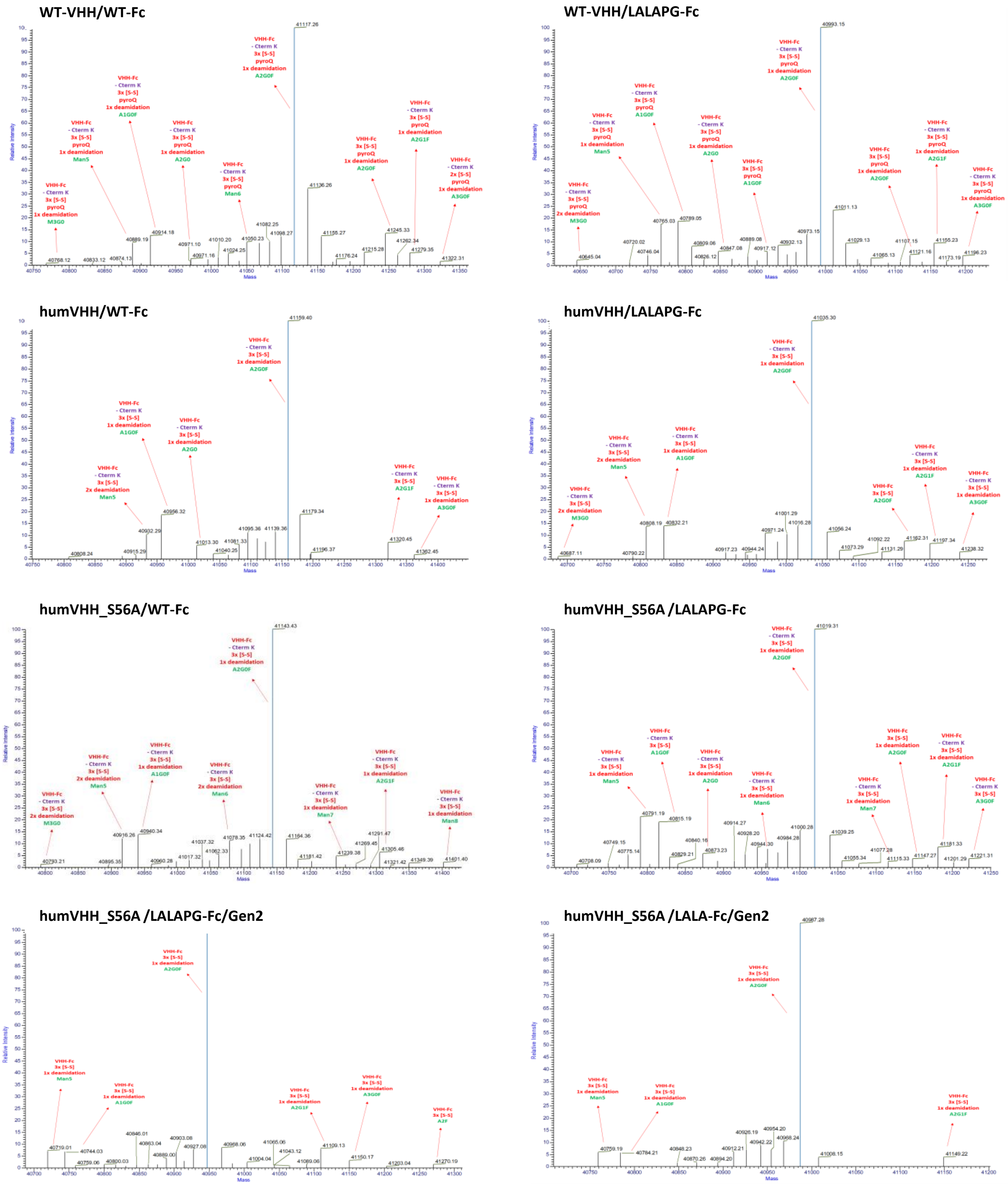


**Supplementary Figure 7. Intact protein mass spectrometry analysis of different VHH72-Fc constructs.** Proteins were first reduced, then separated with reversed phase liquid chromatography, and finally analyzed with an Orbitrap mass spectrometer. The panels depict the deconvoluted spectra of the different protein constructs.

Supplementary Figure 8

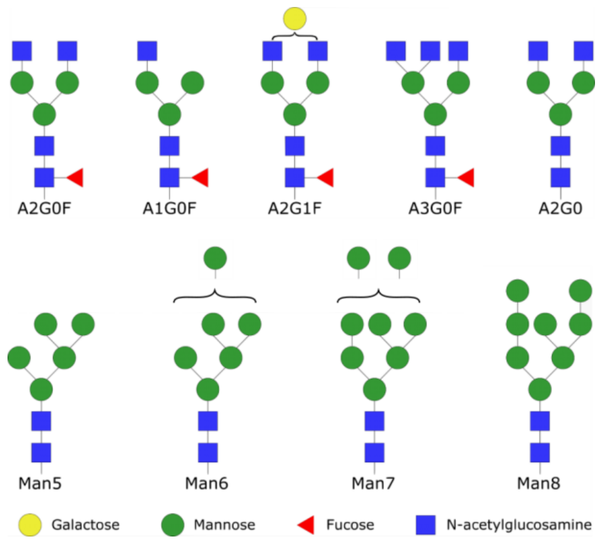


**Supplementary Figure 8. Glycan codes and corresponding glycan composition of the detected N-glycans.**

Supplementary Figure 9

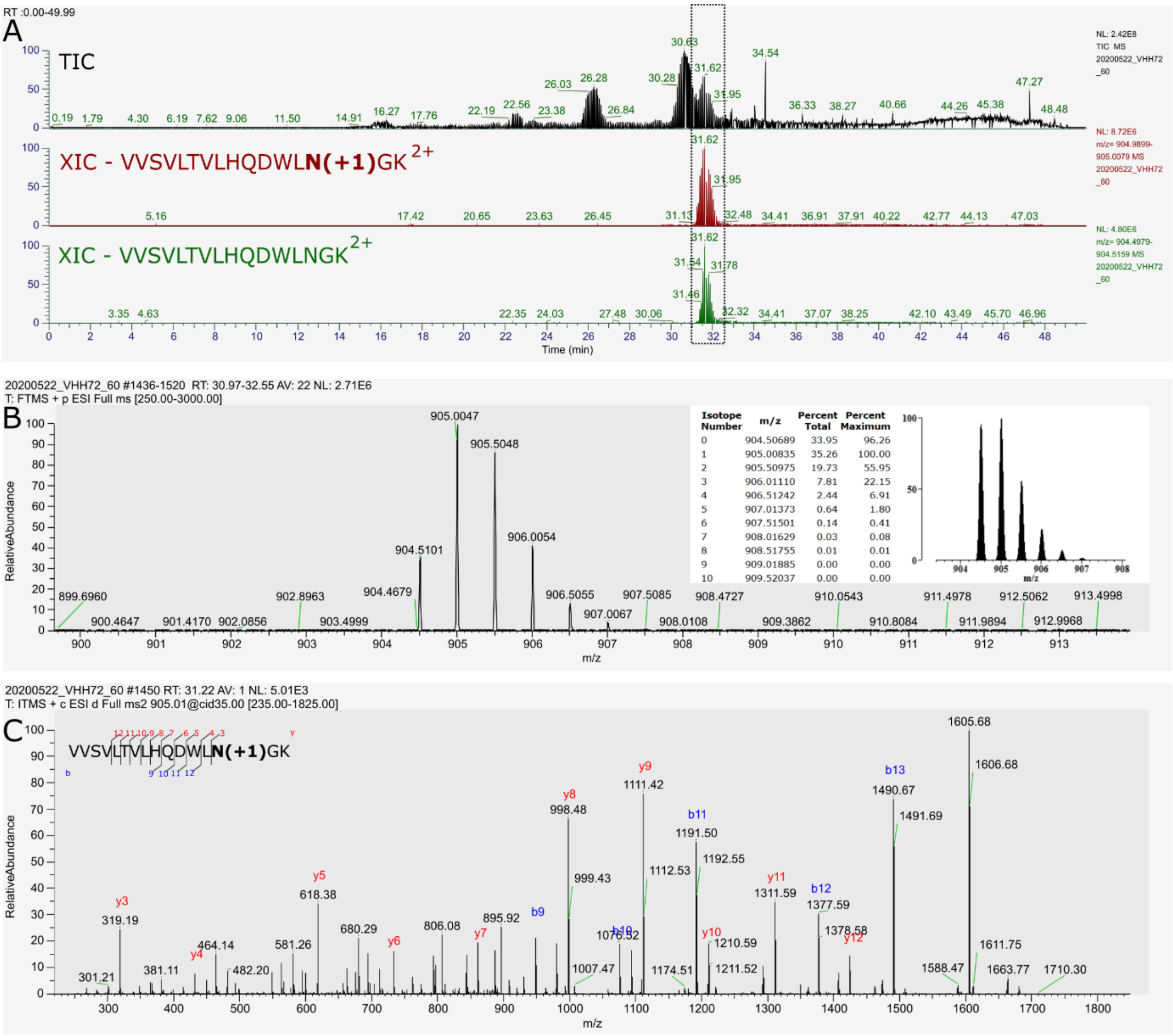


**Supplementary Fig. 9: N227 deamidation analysis of humVHH_S56A/WT-Fc. a.** The upper chromatogram shows the total ion current (TIC) of the digested humVHH_S56A/WT-Fc. The two panels below show the extracted ion chromatograms (XIC) of the tryptic peptide with deamidated N227 (middle) or non-modified N227 (bottom). XICs have a mass tolerance of 10 ppm. **b.** Summation of the time window indicated by the dotted line on panel A, zoomed in on the double charged peptide containing N227. The expected isotopic envelope of the non-modified peptide is shown on the right, generated by *MS-Isotope in ProteinProspector 6.2.3.* The distorted experimental isotope envelope is indicative for deamidation. **c.** CID MS2 spectrum of 905.01 Da precursor peak. Fragment ions match with the theoretical y and b ion series of the N227 deamidated peptide.

Supplementary Figure 10

a

**SEC-MALS of humVHH_S56A/LALAPG-Fc/Gen2 after 0 days**

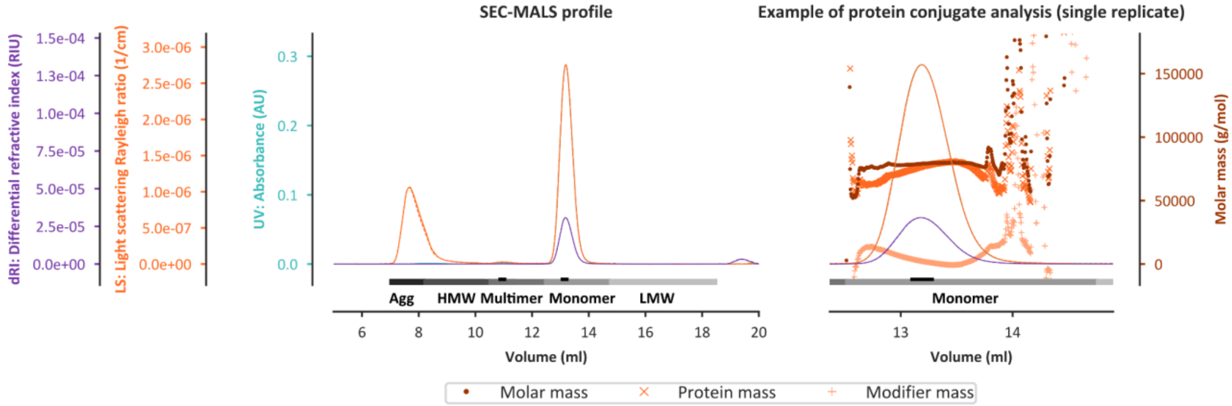


**SEC-MALS of humVHH_S56A/LALAPG-Fc/Gen2 after 10 days at 40 °C**

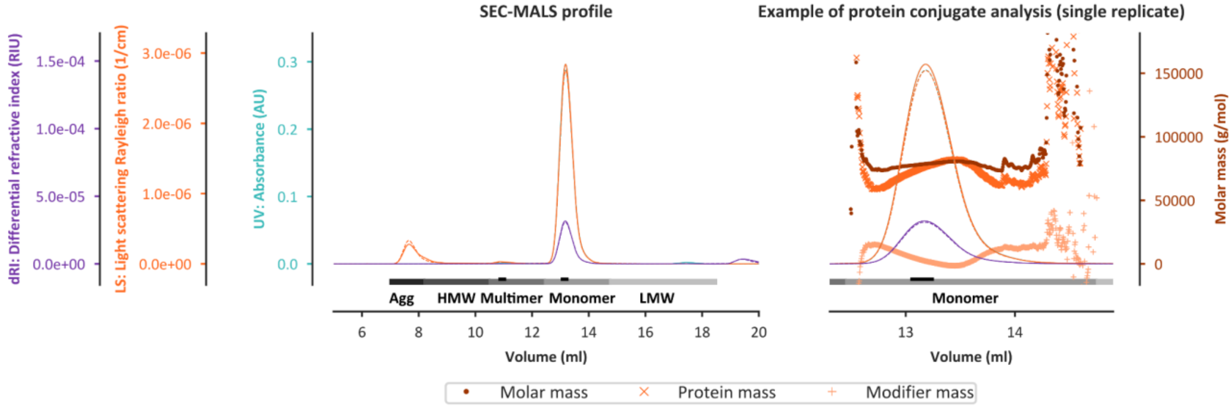


b

**SEC-MALS of (humVHH_S56A)$_2$/LALAPG-Fc/Gen2 after 0 days**

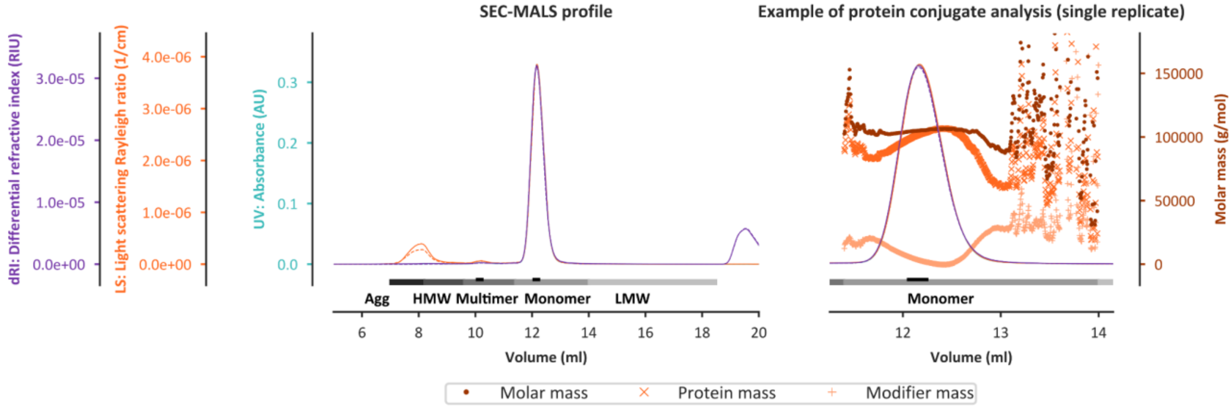


**SEC-MALS of (humVHH_S56A)$_2$/LALAPG-Fc/Gen2 after 10 days at 40 °C**

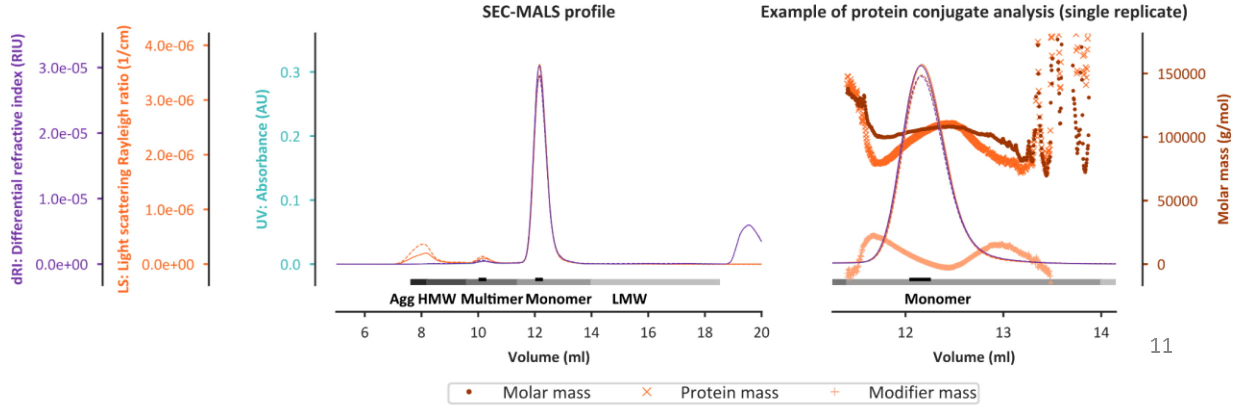

Supplementary Figure 10 - continued

c

**SEC-MALS of humVHH/LALA-Fc/Gen2 after 0 days**

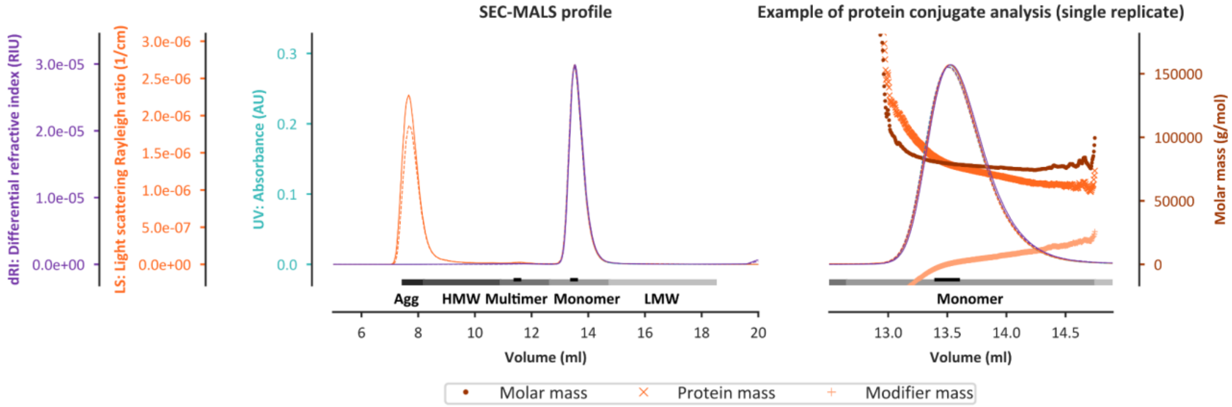


**SEC-MALS of humVHH/LALA-Fc/Gen2 after 10 days at 40 °C**

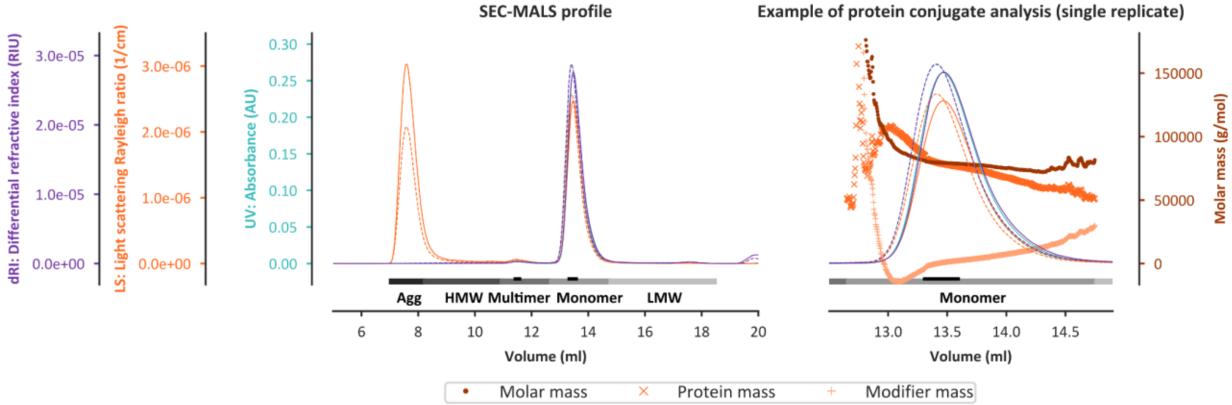


d

**SEC-MALS of humVHH_S56A/LALA-Fc/Gen2 after 0 days**

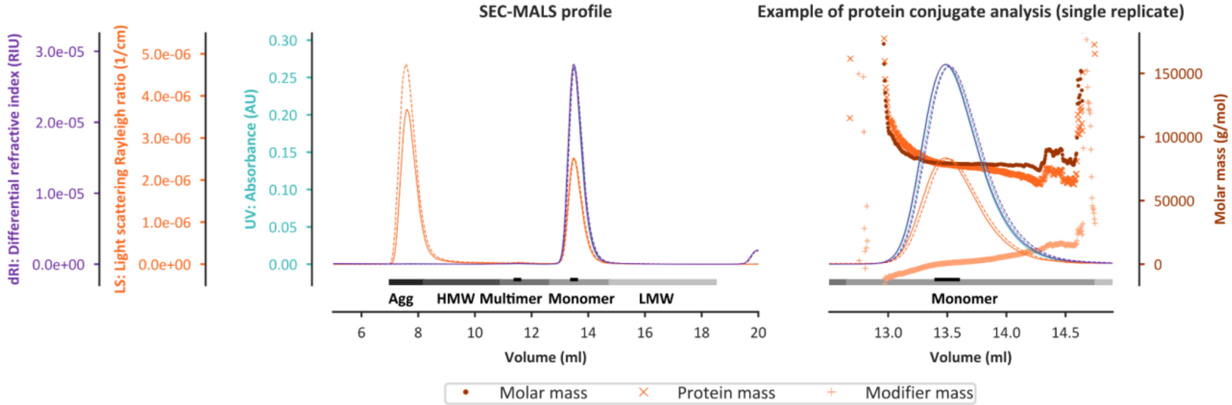


**SEC-MALS of humVHH_S56A/LALA-Fc/Gen2 after 10 days at 40 °C**

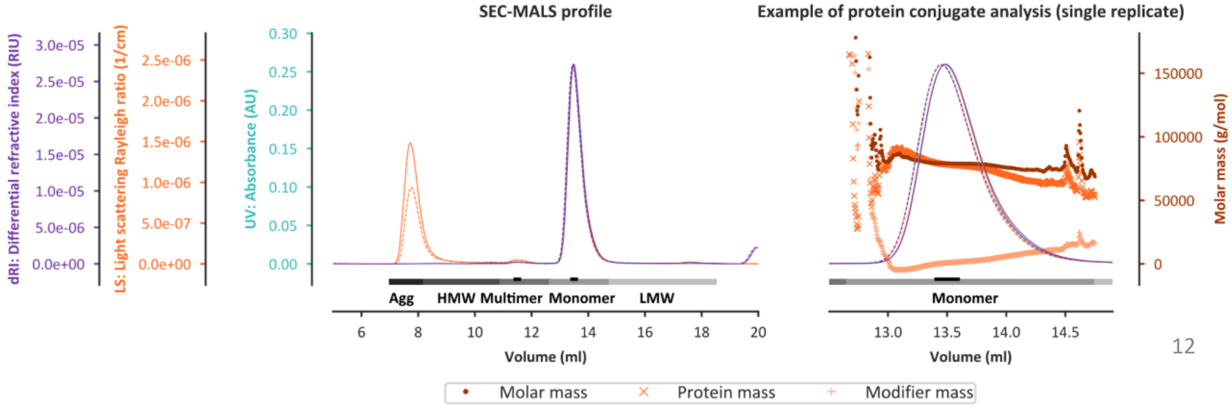

**Supplementary Fig. 10. VHH72-Fc constructs are stable in accelerated stress tests.** Ten day-storage at 40°C causes no major changes in SEC-MALS profiles of 1 mg/ml of **a.** humVHH_S56A/LALAPG-Fc/Gen2, **b.** (humVHH_S56A)2/LALAPG-Fc/Gen2, **c.** humVHH/LALA-Fc/Gen2, and **d.** humVHH_S56A /LALA-Fc/Gen2 in PBS. Complete SEC-MALS profiles of independent duplicates (solid and dashed lines). Peaks for quantitative analysis are indicated in grey below the spectra, peak apex indicated in black for qualitative analysis (results in Supplementary Table 7). The protein conjugate analysis was performed based on the differential extinction coefficients and refractive index values of proteins versus conjugated glycan modifiers.

Supplementary Figure 11

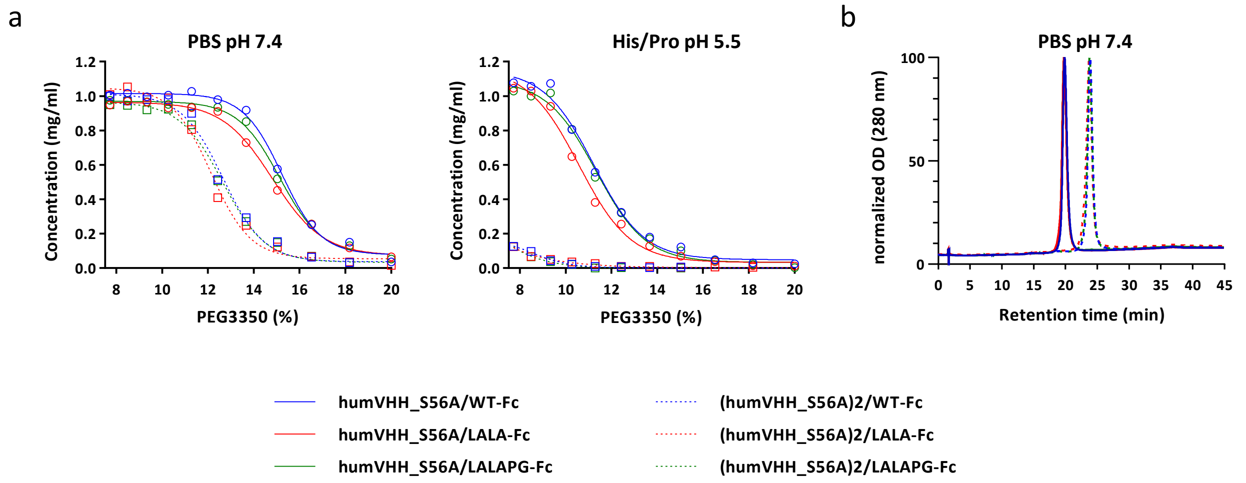


**Supplementary Figure 11. Tetravalent (humVHH_S56A)$_2$/Fc constructs show reduced solubility and greater apparent hydrophobicity compared to the bivalent constructs. a.** PEG aggregation assay to assess solubility of bivalent versus tetravalent VHH-Fc constructs in PBS pH 7.4 (left) or a 50 mM histidine and 250 mM proline buffer pH 5.5 (right). **b**. Hydrophobic interaction chromatography of bivalent versus tetravalent VHH-Fc constructs in 50 mM phosphate and 0.8M ammonium sulfate pH 7.4, eluted in a 45 min linear gradient to zero ammonium sulfate.

Supplementary Figure 12

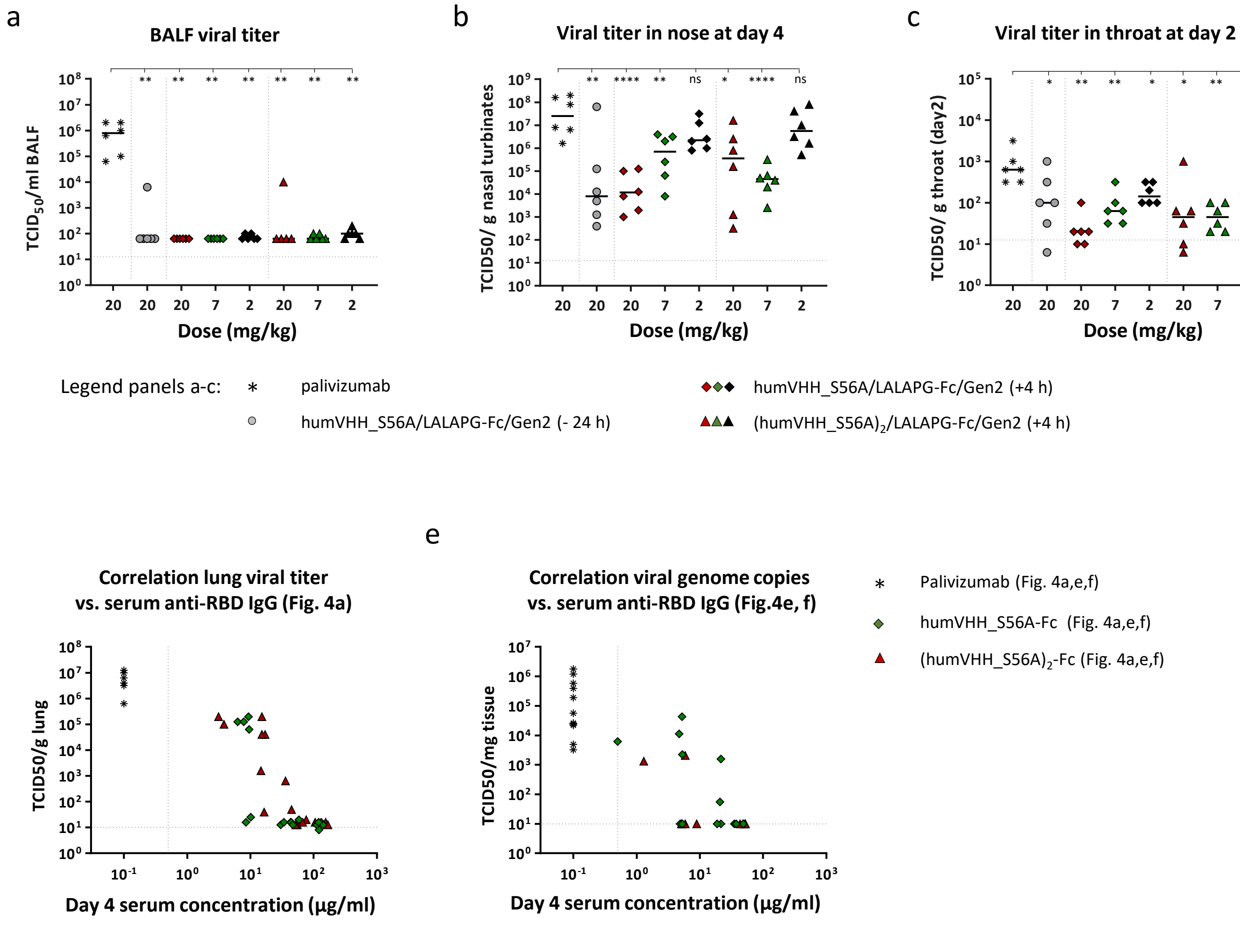


**Supplementary Figure 12. Therapeutic administration of VHH72-Fc constructs reduces the SARS-CoV-2 viral burden in BALF, nose and throat of Syrian hamsters. a-c.** Hamsters were challenged with 1x$10^4$ PFU of BetaCoV/Munich/BavPat1/2020 and 4 hours later injected intraperitoneally with 20, 7 or 2 mg/kg of humVHH_S56A/LALAPG-Fc/Gen2 or $(humVHH\_S56A)_2$/LALAPG-Fc/Gen2. The negative control group was treated with 20 mg/kg of palivizumab, injected 4 hours after the challenge infection; hamsters in a prophylactic control group received 20 mg/kg of humVHH_S56A/LALAPG-Fc/Gen2 one day before the challenge. (**a**) Infectious virus titers in bronchoalveolar lavage and (**b**) nasal turbinates determined on day 4 after infection. (**c**) Viral titers in the throat sampled on day 2 after the challenge. **d, e**. Correlation between the serum concentration of RBD-binding antibodies and infectious virus in the lungs sampled on day 4 after challenge. Data in panels (**d**) and (**e**) are compiled from respectively the therapeutic experiments shown in Fig.4a and Fig.4e and f. Data were analyzed with the Mann-Whitney U-test using GraphPad Prism software. *, $p< 0.05$; **, $p<0.01$; ***, $p<0.001$; ****, $p<0.0001$.

Supplementary Figure 13

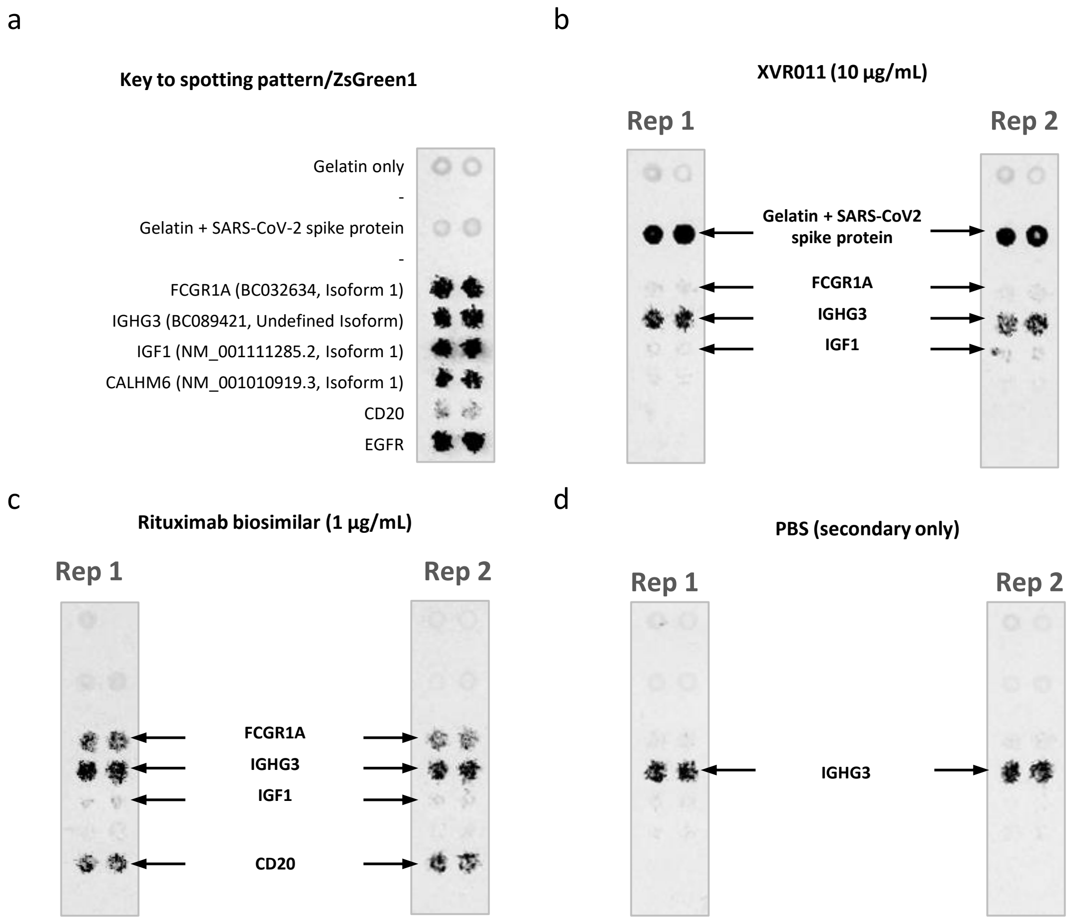


**Supplementary Figure 13. humVHH_S56A/LALA-Fc/Gen2 (= XVR011) is not poly-reactive.** XVR011 at 10 µg/ml was assessed for binding against 5475 human plasma membrane proteins and cell surface tethered secreted proteins plus 371 human heterodimers expressed in transfected HEK293 cells. The confirmation screen for the initial hits of the library screen is shown. **a**. Key to the microarray spotting pattern. Gelatin only and gelatin + SARS-CoV-2 spike protein were spotted. The other spots represent cells that were transfected with an expression plasmid for ZsGreen1 and the indicated respective cDNAs. **b**. Reactivity of XVR011 at 10 µg/ml followed by AlexaFluor647 labelled anti-human IgG Fc detection antibody, followed by fluorescence imaging. **c**. Reactivity of Rituximab 1 µg/ml with the microarray with detection as in (**b**). **d**. Reactivity with the microarray of the secondary antibody (PBS instead of primary antibody). Rep: replicate.

### Supplementary Figure 14

a

Rituximab binding to FcγRI

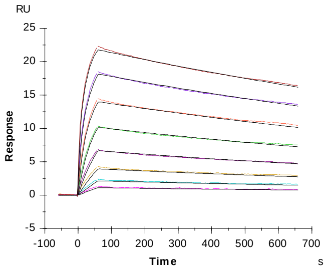


XVR011 binding to FcγRI

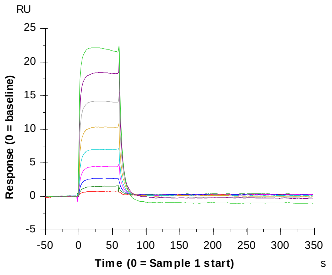


b

Rituximab binding to FcγRIIa (H167)

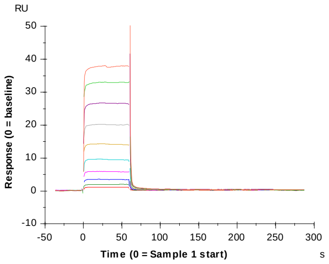


XVR011 binding to FcγRIIa (H167)

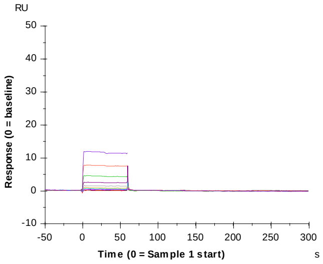


c

Rituximab binding to FcγRIIa (R167)

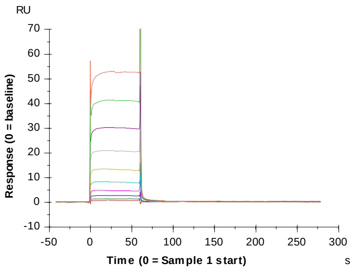


XVR011 binding to FcγRIIa (R167)

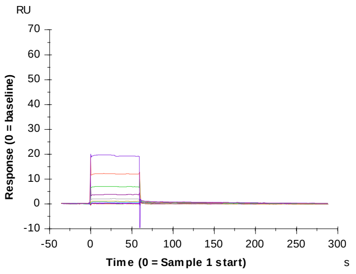


d

Rituximab binding to FcγRIIb

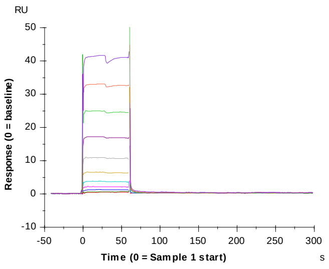


XVR011 binding to FcγRIIb

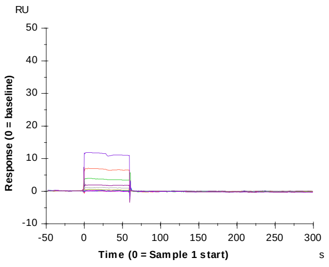

Supplementary Figure 14 cont.

e

Rituximab binding to FcγRIIIa (V176) XVR011 binding to FcγRIIIa (V176)

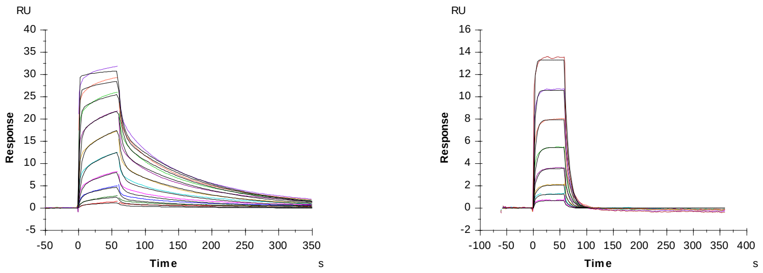


f

Rituximab binding to FcγRIIIa (F176) XVR011 binding to FcγRIIIa (F176)

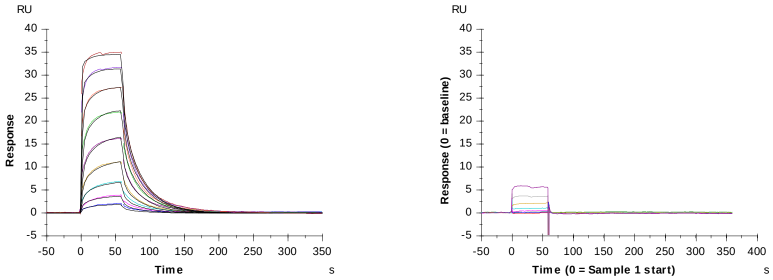


g

Rituximab binding to FcγRIIIb XVR011 binding to FcγRIIIb

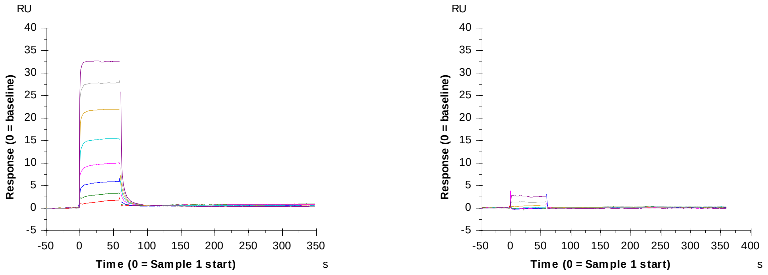


**Supplementary Fig. 14. Surface plasmon resonance (SPR) sensorgrams for the binding of XVR011 and rituximab to immobilized Fcγ receptors. a-g.** Binding of rituximab (left) and XVR011 (right) to (**a**) FcγRI, (**b**) FcγRIIa (H167), (**c**) FcγRIIa (R167), (**d**) FcγRIIb, (**e**) FcγRIIIa (V176), (**f**) FcγRIIIa (F176), and (**g**) to FcγRIIIb.

Supplementary Figure 15

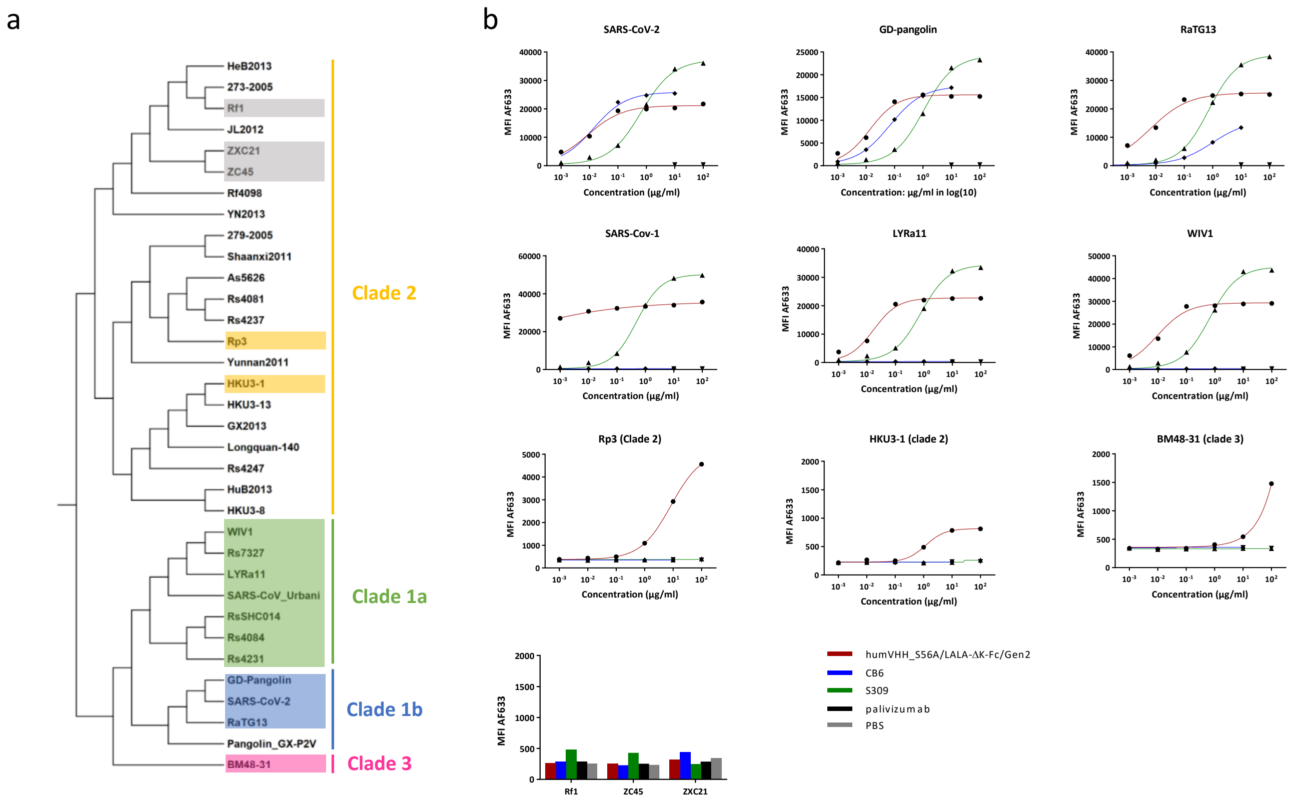


**Supplementary Figure 15. humVHH_S56A/LALA-Fc/Gen2 binds to the RBD of a diverse range of Sarbecoviruses. a.** Cladogram (UPGMA method) based on the RBD of SARS-CoV-1-related, SARS-CoV-2-related and clade 2 and clade 3 Bat SARS-related Sarbecoviruses. The colored boxes indicate the RBD variants that are bound by humVHH_S56A/LALA-Fc/Gen2 as determined by flow cytometry of either yeast cells that display the indicated RBD variants or HEK293T cells that express SARS-CoV-1 spike proteins in which the RBD is substituted by the indicated RBD variants. The grey boxes indicate the RBD variants for which no binding of humVHH_S56A/LALA-Fc/Gen2 could be observed. **b.** Analysis of the binding of humVHH_S56A/LALA-Fc/Gen2, S309, CB6 and palivizumab to *Saccharomyces cerevisiae* cells that display the RBD of the indicated Sarbecoviruses. The graphs show the MFI of AF633 conjugated anti-human IgG that was used to detect the binding of dilution series of the tested antibodies to *S. cerevisiae* cells that express the RBD derived from the indicated Sarbecoviruses.

**Supplementary Table 1. Overview of the VHH72-Fc constructs and their $PRNT_{50}$ SARS-CoV-2 neutralization values**.

| # | Short name | VHH72-Fc variant | $PRNT_{50}$ (µg/ml) |
|---|---|---|---|
| 1 | WT-VHH/12GS-WT-Fc | VHH72-GS$(G_4S)_2$-hIgG1hinge-hIgG1Fc | 0.68 |
| 2 | WT-VHH/WT-Fc | VHH72-GS-hIgG1hinge-hIgG1Fc | 1.01 |
| 3 | WT-VHH/LALAPG-Fc | VHH72-GS-hIgG1hinge-hIgG1Fc_LALAPG | 0.57 |
| 4 | humVHH/WT-Fc | VHH72_h1-GS-hIgG1hinge-hIgG1Fc | 0.94 |
| 5 | $(humVHH)_2$/WT-Fc | VHH72_h1-$(G_4S)_3$-VHH72_h1-GS-hIgG1hinge-hIgG1Fc | 0.10 |
| 6 | humVHH/LALAPG-Fc | VHH72_h1-GS-hIgG1hinge-hIgG1Fc_LALAPG | 1.22 |
| 7 | humVHH_S56A/WT-Fc | VHH72_h1_S56A-GS-hIgG1hinge-hIgG1Fc | 0.12 |
| 8 | humVHH_S56A/LALAPG-Fc | VHH72_h1_S56A-GS-hIgG1hinge-hIgG1Fc_LALAPG | 0.13 |
| 9 | humVHH/LALA-Fc/Gen2 | VHH72_h1_E1D-$(G_4S)_2$-hIgG1hinge_EPKSCdel-hIgG1Fc_LALA_K447del | ND |
| 10 | humVHH_S56A /LALAPG-Fc/Gen2 | VHH72_h1_E1D_S56A-$(G_4S)_2$-hIgG1hinge_EPKSCdel-hIgG1Fc_LALAPG_K477del | 0.11 |
| 11 | $(humVHH\_S56A)_2$/LALAPG-Fc/Gen2 | VHH72_h1_E1D_S56A-$(G_4S)_3$-VHH72_h1_E1D_S56A-GS-hIgG1hinge_EPKSCdel-hIgG1Fc_LALAPG_K477del | 0.02 |
| 12 | humVHH_S56A/LALA-Fc/Gen2 | VHH72_h1_E1D_S56A-$(G_4S)_2$-hIgG1hinge_EPKSCdel-hIgG1Fc_LALA_K477del | 0.09 |

PRNT: plaque reduction neutralization assay

ND: not determined.

Design 12 (humVHH_S56A/LALA-Fc/Gen2) was chosen for the development of the drug candidate that was cGMP manufactured and formulated as XVR011.

**Supplementary Table 2**. **Binding affinity of VHH72-Fc variants to immobilized mouse Fc-fused SARS-CoV-2 RBD (RBD-mFc).** Apparent kinetics of the 2:2 interaction is based on a global 1:1 fit of the replicate (n = 2) data; values are the averages of the replicates.

| # | VHH-Fc variant | $K_D$ (M) | $k_{on}$ (1/Ms) | $k_{off}$ (1/s) |
|---|---|---|---|---|
| 5 | $(humVHH)_2$/WT-Fc | <1.0E-12 | 2.65E+06 | <1.0E-07 |
| 10 | humVHH_S56A /LALAPG-Fc/Gen2 | 4.08E-11 | 1.52E+06 | 6.16E-05 |
| 11 | $(humVHH\_S56A)_2$/LALAPG-Fc/Gen2 | <1.0E-12 | 2.48E+06 | <1.0E-07 |

**Supplementary Table 3. Peptide mapping results of the different VHH72-Fc constructs.** LysC-Trypsin codigested protein was analyzed with LC-MS/MS, and peptides and peptide modifications detected with BioPharma Finder™.

| # | VHH-Fc variant | Sequence coverage | Modifications | | | |
|---|---|---|---|---|---|---|
| | | | Pyro-Glu | N-glycosylation | Glycation | Deamidation |
| 2 | WT-VHH/WT-Fc | 84.4% | Q1 (98.13%) | N209 | K238 | N227 |
| 3 | WT-VHH/LALAPG-Fc | 82.5% | Q1 (92.28%) | N209 | K160, K200 | N227 |
| 4 | humVHH/WT-Fc | 83% | / | N209 | K160, K238 | N227 |
| 6 | humVHH/LALAPG-Fc | 83.3% | / | N209 | K160, K238 | N227 |
| 7 | humVHH_S56A/WT-Fc | 85.2% | / | N209 | K238 | N227 |
| 8 | humVHH_S56A /LALAPG-Fc | 84.7% | / | N209 | K160, K238 | N227 |
| 10 | humVHH_S56A /LALAPG-Fc/Gen2 | 83.9% | / | N212 | / | N230 |
| 12 | humVHH_S56A /LALA-Fc/Gen2 | 80.1 | / | N212 | / | N230 |

**Supplementary Table 4**. **Expression levels of VHH72-Fc constructs in transiently transfected ExpiCHO cells.**

| # | VHH-Fc variant | ExpiCHO-S yield (mg/l) |
|---|---|---|
| 1 | WT-VHH/12GS-WT-Fc | 638 |
| 2 | WT-VHH/WT-Fc | 403 |
| 3 | WT-VHH/LALAPG-Fc | 773 |
| 4 | humVHH/WT-Fc | 768 |
| 5 | $(humVHH)_2$/WT-Fc | 803 |
| 6 | humVHH/LALAPG-Fc | 1033 |
| 7 | humVHH_S56A/WT-Fc | 1215 |
| 8 | humVHH_S56A /LALAPG-Fc | 806 |

**Supplementary Table 5. VHH72-Fc fusion proteins have very low aggregation propensity in PBS, also after introduction of humanizing and affinity-increasing mutations, both in wildtype human IgG1-Fc context and after LALAPG modification.**

| ID | VHH72-Fc variant | Quantitative analysis of total profile | | | | | Qualitative analysis of monomer peak | |
|---|---|---|---|---|---|---|---|---|
| | | Peak quantitation (%) | | | | | Molar mass (kDa) | |
| | | Monomer | Multimer | HMW | LMW | Aggregates | Predicted* | MALS |
| 2 | WT-VHH/WT-Fc | 97.8 | 0.5 | 0.2 | 1.2 | 0.2 | 79.7 | 79.3 |
| 3 | WT-VHH/LALAPG-Fc | 98.4 | 0.4 | 0.1 | 1.0 | 0.1 | 79.4 | 78.2 |
| 4 | humVHH/WT-Fc | 98.8 | 0.4 | 0.0 | 0.9 | 0.0 | 79.7 | 78.9 |
| 6 | humVHH/LALAPG-Fc | 98.9 | 0.6 | 0.0 | 0.5 | 0.0 | 79.5 | 77.9 |
| 7 | humVHH_S56A/WT-Fc | 97.4 | 0.6 | 0.2 | 1.7 | 0.1 | 79.7 | 78.8 |
| 8 | humVHH_S56A /LALAPG-Fc | 97.2 | 0.4 | 0.4 | 1.9 | 0.2 | 79.5 | 77.7 |

SEC-MALS analysis of purified VHH72-Fc samples. The peak corresponding to the molar weight of an assembled bivalent VHH-Fc protein was indicated as 'monomer peak'. Peak quantitation (%) is based on the refraction signal. HMW = high molecular weight species; LMW = low molecular weight species. *Molar mass as predicted by the ExPASy ProtParam tool.

**Supplementary Table 6. Bivalent and tetravalent VHH72-Fc constructs occur as >96% monomers upon 10 days stress testing at 40 °C in PBS (without formulation).**

| **Sample** | | | | **Quantitative analysis of total profile** | | | | | | | | | | **Qualitative analysis of monomer peak** | | |
|---|---|---|---|---|---|---|---|---|---|---|---|---|---|---|---|---|
| **#** | **VHH72-Fc variant** | **Days at 40 °C** | **n** | **Peak quantitation (%)** | | | | | **Polydispersity** | | | | | Molar mass (kDa) | | Hydrodynamic radius rh(Q)w (nm) |
| | | | | Monomer | Multimer | HMW | LMW | Agg | Monomer | Multimer | HMW | LMW | Agg | Predicted* | MALS | |
| **10** | humVHH_S56A /LALAPG-Fc/Gen2 | 0 | 2 | 98.1 | 0.2 | 0.6 | 0.9 | 0.3 | 1.006 | 1.297 | 1.288 | 1.536 | 1.964 | 79.1 | 77.9 | 4.3 |
| | | 10 | 2 | 98.7 | 0.4 | 0.1 | 0.9 | 0.0 | 1.003 | 1.972 | 1.990 | 13.247 | 1.989 | | 77.7 | 4.1 |
| **11** | $(humVHH\_S56A)_2$/LALAPG-Fc/Gen2 | 0 | 2 | 97.5 | 1.4 | 0.5 | 0.6 | 0.2 | 1.007 | 1.160 | 1.202 | 3.380 | 1.366 | 107.2 | 104.3 | 5.3 |
| | | 10 | 2 | 95.1 | 2.6 | 0.6 | 1.7 | 0.1 | 1.008 | 1.187 | 1.485 | 15.592 | 1.454 | | 104.7 | 5.1 |
| **9** | humVHH/LALA-Fc/Gen2 | 0 | 2 | 98.2 | 0.5 | 0.3 | 1.2 | 0.0 | 1.011 | 2.218 | 2.579 | 2.465 | 1.554 | 79.2 | 79.3 | 4.3 |
| | | 10 | 2 | 95.1 | 1.5 | 0.7 | 2.7 | 0.2 | 1.005 | 1.249 | 2.646 | 2.251 | 1.896 | | 78.8 | 4.3 |
| **12** | humVHH_S56A/LALA-Fc/Gen2 | 0 | 2 | 97.9 | 0.6 | 0.3 | 1.1 | 0.2 | 1.017 | 1.734 | 2.106 | 1.902 | 7.982 | 79.1 | 79.3 | 4.5 |
| | | 10 | 2 | 96.1 | 1.1 | 0.6 | 2.1 | 0.1 | 1.003 | 1.172 | 2.069 | 2.345 | 1.922 | | 78.8 | 4.3 |

SEC-MALS analysis of purified VHH72-Fc samples. The peak corresponding to the molar weight of an assembled bivalent VHH-Fc protein was indicated as 'monomer peak'. Peak quantitation (%) is based on the refraction signal. Qualitative analysis of the monomer peak was performed on the 200 µl peak elution fraction, as indicated in black in Figure 3e. Values reported are the averages of duplicate measurements of two independently stress tested samples. HMW = high molecular weight species; LMW = low molecular weight species; agg = aggregates. *Molar mass as predicted by the ExPASy ProtParam tool.

**Supplementary Table 7**. **Biophysical characteristics of bivalent versus tetravalent VHH72-Fc variants.**

| Short name | VHH72-Fc variant | HIC retention time (min) | PEG assay (% midpoint) PBS pH 7.4 | His/Pro pH 5.5 | pI Theoretical* | cIEF |
|---|---|---|---|---|---|---|
| humVHH_S56A/WT-Fc | VHH72_h1_E1D_S56A-$(G_4S)_2$-hIgG1hinge-hIgG1Fc | 19.9 | 15.2 | 11.3 | 6.0 | 6.3 |
| humVHH_S56A/LALA-Fc | VHH72_h1_E1D_S56A-$(G_4S)_2$-hIgG1hinge-hIgG1Fc_LALA | 19.8 | 14.8 | 10.7 | 6.0 | 6.3 |
| humVHH_S56A/LALAPG-Fc | VHH72_h1_E1D_S56A-$(G_4S)_2$-hIgG1hinge-hIgG1Fc_LALAPG | 19.8 | 15.1 | 11.4 | 6.0 | 6.3 |
| $(humVHH\_S56A)_2$/WT-Fc | VHH72_h1_E1D_S56A-$(G_4S)_3$-VHH72_h1_S56A-GS-hIgG1hinge-hIgG1Fc | 23.9 | 12.7 | <7.7 | 5.5 | 5.9 |
| $(humVHH\_S56A)_2$/LALA-Fc | VHH72_h1_E1D_S56A-$(G_4S)_3$-VHH72_h1_S56A-GS-hIgG1hinge-hIgG1Fc_LALA | 23.7 | 12.1 | <7.7 | 5.5 | 5.9 |
| $(humVHH\_S56A)_2$/LALAPG-Fc | VHH72_h1_E1D_S56A-$(G_4S)_3$-VHH72_h1_S56A-GS-hIgG1hinge-hIgG1Fc_LALAPG | 23.8 | 12.6 | <7.7 | 5.5 | 5.9 |

A greater hydrophobic interaction chromatography (HIC) retention time indicates increased apparent hydrophobicity. Solubility at increasing polyethylene glycol (PEG) concentrations was determined in two buffer systems (PBS pH 7.4 and 50 mM histidine, 250 mM proline, pH 5.5), the midpoint of the sigmoidal PEG dose-response fit is indicated as '% midpoint'. A higher PEG midpoint score suggests greater solubility at high concentrations. Approximately 90% of the tetravalent sample formulated in 50 mM histidine, 250 mM proline pH 5.5 buffer aggregated at the lowest PEG3350 concentration (7.7 %), therefore a midpoint could not be determined. The isoelectic point (pI) was determined by capillary isoelectric focusing (cIEF). *Theoretical pI as calculated using the MPCT tool (ABIM, France).

**Supplementary Table 8. Binding affinity of XVR011 and Rituximab to immobilized recombinant human Fcγ receptors determined by SPR.**

| Fcγ receptor | XVR011 $K_D$ (M) ± SD (n=3) | Rituximab $K_D$ (M) ± SD (n=3) |
|---|---|---|
| FcγRI | 1.76 ± 0.01 x $10^{-6}$ | $K_{D1}$: 7.73 x $10^{-10}$; $K_{D2}$: 4.76 ± 0.27 x $10^{-9}$* |
| FcγRIIa (H167) | > 20 μM | 1.28 ± 0.14 x $10^{-6}$ |
| FcγRIIa (R167) | > 20 μM | 2.90 ± 0.30 x $10^{-6}$ |
| FcγRIIb | > 20 μM | 5.00 ± 0.43 x $10^{-6}$ |
| FcγRIIIa (V176) | 5.66 ± 0.15 x $10^{-6}$ | 1.43 ± 0.11 x $10^{-7}$ |
| FcγRIIIa (F176) | > 20 μM | 6.98 ± 0.79 x $10^{-7}$ |
| FcγRIIIb | > 20 μM | 1.76 ± 0.11 x $10^{-6}$ |

* equilibrium dissociation constants ($K_{D1\ an}$ $K_{D2}$) of rituximab interacting with FcγRI, fitted to a heterogenous ligand model.

All other data were fitted to a 1:1 binding model.

Listed are mean values ± SD of n=3 independent experiments.
